# Two Small Proteins Enable N_2_O Reduction in Acidic Environments

**DOI:** 10.64898/2025.12.16.694761

**Authors:** Xinhui Wang, Leying Meng, Hu Li, Yifang Zhang, Minghou Li, Baoyu Xiang, Jing Zhu, Chen Li, Chen Yang, Siyu Yu, Xiaoyan Pang, Jianwen Wei, Menghui Zhang, Lars R Bakken, Åsa Frostegård, Lin Zhang, Xiaojun Zhang

## Abstract

Denitrifying bacteria are major contributors to nitrous oxide (N_2_O) emissions, which rise with global soil acidification, likely because low pH impairs maturation of the periplasmic enzyme N_2_O reductase (NosZ) in most bacteria. While some bacteria can reduce N_2_O under acidic conditions, the underlying mechanism remain unclear. Here, we isolated bacteria from acidic soils and identified strains that efficiently reduce N_2_O at low pH. They carry two novel proteins, NosP and NosQ, along with NosZ clade I. Functional analyses of deletion mutants demonstrated that NosP and NosQ are essential for the maturation of NosZ at low pH, but are not required at circumneutral pH. Cryo-EM analysis indicates that NosPQ forms a protective complex with NosZ, safeguarding its maturation under acidic conditions. Metagenomic analysis shows that *nosPQ* genes are widely distributed across diverse bacterial taxa and soil types. Our findings reveal a key molecular adaptation mechanism for N_2_O reduction in acidic environments and provide valuable insights for developing efficient N_2_O mitigation strategies.

## MAIN

The year 2024 marked a critical turning point in the trajectory of climate change, with global temperatures exceeding 1.5□ above pre-industrial levels for the first time^1^. This landmark event highlights the urgent need to mitigate greenhouse gas emissions^2,3^, and a key priority for the agricultural sector is to reduce the nitrous oxide (N_2_O) emission as it is a major contributor to the sector’s global warming impact ^4^. Driven by increasing demands for food, energy, and waste management, N_2_O emissions are projected to rise, posing a significant challenge to climate mitigation efforts^5,6^. Agricultural soils, especially those subjected to intensive nitrogen fertilization, are major anthropogenic sources of N_2_O^7,8^, with soil acidification further exacerbating these emissions^9,10^.

Nitrous oxide reductase (NosZ), the only known enzyme responsible for catalyzing the reduction of N_2_O to N_2_^11^, plays a key role in curbing N_2_O emissions. Long before NosZ was identified, Delwich and Wijler^12^ demonstrated that N_2_O reduction in soil is hampered by soil acidification. This inhibition of N_2_O reduction at low pH has since been confirmed through numerous studies of soils and cultures of denitrifying bacteria^13–15^, and was long attributed to a direct inhibition of NosZ. A more plausible explanation is that the majority of denitrifying bacteria fail to synthesize functional NosZ at low pH, most likely because the maturation of the enzyme in the periplasm is hampered by pH<6. This hypothesis is derived from a series of observations—conducted on both pure cultures and soil microbial communities^13,15–17^—which demonstrate that functional NosZ synthesis fails at low pH, even though *nosZ* gene transcription occurs. The enzyme is, however, functional at low pH if it has been synthesized at circumneutral pH, albeit not at an optimal rate^13,18^. The maturation of the NosZ apoprotein, which is produced in the cytoplasm, takes place in the periplasm where the pH is similar to the external environment. The maturation process includes the insertion of Cu ions into the active sites Cu_A_ and Cu_Z_, a step that requires several maturation proteins^19,20^. Some bacteria have apparently evolved mechanisms to synthesize functional NosZ at low pH, however. Evidence for this includes observations of N_2_O reduction in several acidic soils across diverse ecosystems^21–27^, and the discovery of bacteria capable of N_2_O reduction at acidic pH^28–30^, including one that carries a novel clade III NosZ ^31^. Deciphering the mechanism for such adaptation would enhance our understanding of nitrogen cycling in acidic environments and open avenues for mitigating N_2_O emissions from acid soils.

Here, we report the isolation of diverse bacteria capable of reducing N_2_O at acidic pH. Further investigations proved that these bacteria carry previously uncharacterized genes in their *nos* gene clusters, encoding auxiliary proteins which were evidently essential for maturation of NosZ clade I under acidic conditions. These genes were detected in metagenomes from globally distributed acidic soils, suggesting their potentially important role in N_2_O reduction at acidic pH.

### N_2_O reduction in low pH soils

Reduction of N_2_O in soils is influenced by a complex interplay of environmental and biological factors^32^. Acidification, as explained above, has been identified as a key repressor, accounting for the observed inverse correlation between N□O emissions and soil pH^9,33,34^. Despite the prevailing understanding that microbial N_2_O reduction is inhibited under acidic conditions, N_2_O reduction in strongly acidic soils has been reported across various ecosystems^22–25,35^. In the present study we collected two acidic soils (Figure 1a), one forest soil (pH 4.4) and one paddy soil (pH 4.8), and demonstrated their capabilities of N_2_O reduction (Figure 1b, Extended Data Fig. 1a, Supplementary Fig. 1). Hypothetically, N_2_O reduction in such acidic soils may be attributed to the synthesis of functional NosZ in near-neutral microhabitats within the soil matrix. To test whether N_2_O can be reduced when the soil aggregates were destroyed, we dispersed 1 g of the paddy soil in 30 mL of acidic (pH 4.5) soil extract-glucose (SEG) medium, effectively eliminating any near-neutral micro-niches. The slurry was incubated under anoxic conditions with N_2_O and stirred vigorously. The glucose-induced N_2_O reduction rate in the slurries, calculated per g soil, was 3,700 nmol N_2_O g^-1^h^-1^ (Extended Data Fig. 1b), which is two orders of magnitude higher than in the non-amended intact soil. These results suggest that this acidic soil contains bacteria capable of synthesizing NosZ at very low pH. Interestingly, the soil also appeared to harbour bacteria that require circumneutral pH for synthesizing functional NosZ, as evidenced by the immediate enhancement of N_2_O reduction following liming (Add Extended Data Fig. 1c, Supplementary Results chapter 2.1), which is consistent with previous literature reports^36^.

**Fig. 1.**
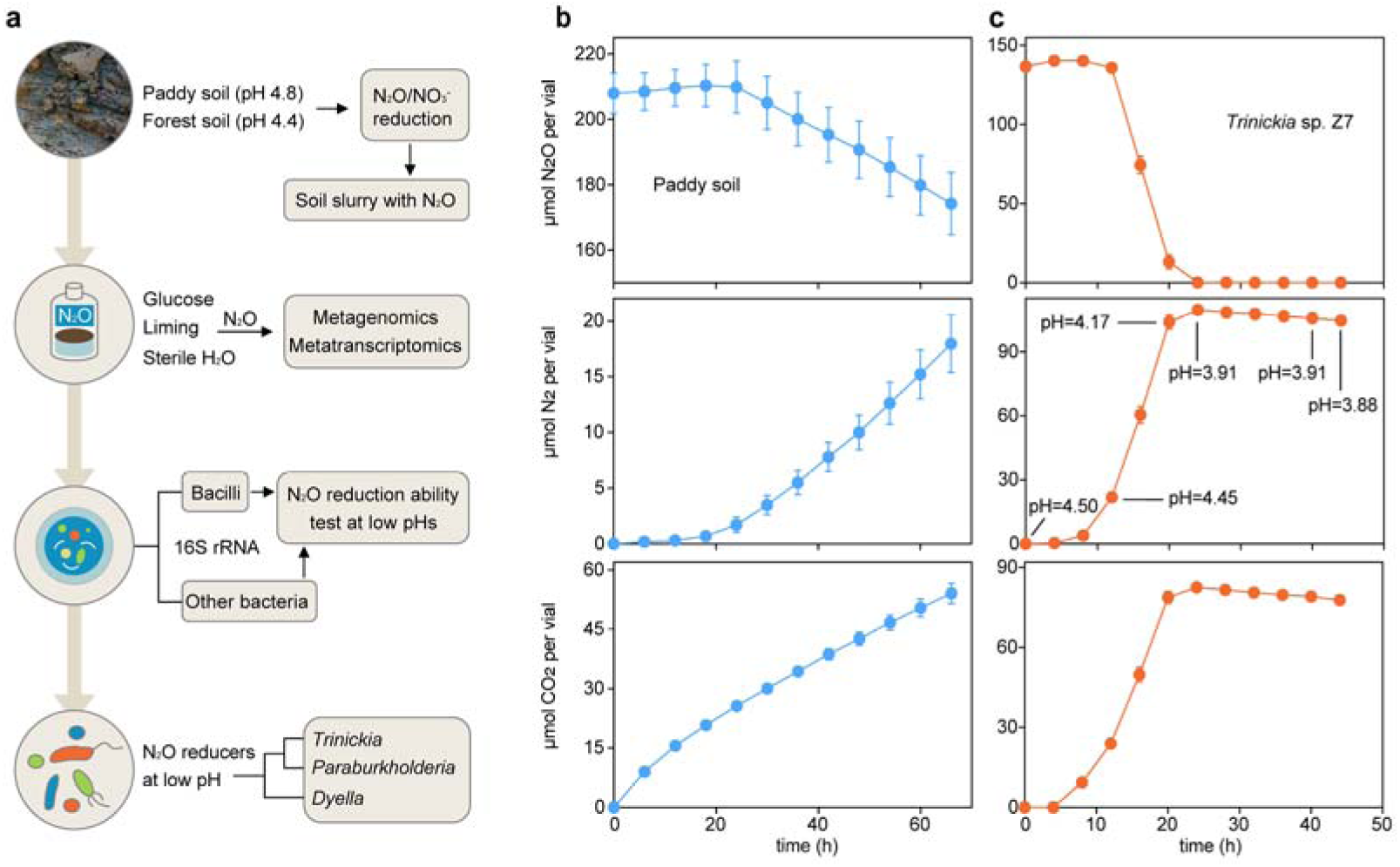
Discovery of acid-tolerant N_2_O reducing bacteria. (**a**) Workflow illustrating the enrichment, isolation and identification of acid-tolerant N_2_O reducing bacteria. Two strongly acidic soils were first found to be able to reduce N_2_O. Different amendments were then tried on paddy soil and the microbial community was analyzed through metagenomics and metatranscriptomics. Diverse isolates were obtained using various isolation methods and assessed for their N_2_O reducing capability at low pH. Finally, the pure isolates belonging to the genera *Trinickia*, *Paraburkholderia*, *Dyella* were identified as acid-tolerant N_2_O reducing bacteria through strictly-controlled pH experiments. (**b**) N_2_O reduction in 30 g (wet weight; ww) of intact paddy soil. Gas kinetics were monitored during anoxic incubation in gas tight vials containing 5 mL of N_2_O in the headspace. (**c**) N_2_O reduction by *Trinickia* sp. Z7 in acidic non-buffered modified Sistrom’s medium with initial pH=4.5. The cells were previously grown in acidic (pH 4.2) Sistrom’s medium under oxic conditions with vigorous stirring (500 rpm). Regular transfers of these cells to new medium (>23 generations) were performed to eliminate the possibility that the inoculum carried previously synthesized N_2_O reductase that could have matured under non-acidic conditions. Subsequently, 300 μL was taken from these precultures and inoculated into 30 mL of Sistrom’s medium and incubated under continuous stirring (500 rpm). Each vial was supplemented with 3 mL of N_2_O in the headspace and pH measurements were carried out in replicates. Each treatment was measured in three replicate vials (n=3). Bars represent SEM.

### Isolation of acid-tolerant N_2_O reducing bacteria

We searched the metagenomes and metatranscriptomes originating from the paddy soil incubations (Extended Data Fig. 1c), sampled at different time points, to identify organisms which could possibly synthesize functional NosZ in this soil. These anoxic incubation with N_2_O included unamended paddy soil (S), paddy soil amended with glucose (G), and paddy soil amended with lime raising the pH to 7.8 (C). Detailed results from the meta-genome and transcriptome analyses are provided in Supplementary Results chapter 2.2 & 2.3.

Meta-omics analyses identified a cluster of Firmicutes that showed extensive *nosZ* transcription in the paddy soil (Extended Data Fig. 2). To selectively isolate such endospore forming bacteria, we performed a heat treatment of a soil suspension followed by dilution plating (aerobic incubation) on four different media (Supplementary Results chapter 2.4). This yielded 40 sequenced and taxonomically distinct isolates belonging to *Bacillus* and *Neobacillus*. All carried *nosZ*, and all were capable of reducing N_2_O at circumneutral pH, but none reduced N_2_O at low pH.

Co-abundance network analysis of the *nosZ* gene revealed an interesting bacterial cluster with *nosZ* gene whose abundance increased in the acidic soil during incubation with N_2_O (Supplementary Fig. 6). Metagenomic binning identified assembled genomes of *Trinickia* and *Noviherbaspirillum* as particularly interesting candidates for targeted isolation. To increase the chances of obtaining as wide a diversity as possible of these *nosZ*-containing, facultative anaerobic organisms, we first performed dilution plating under oxic conditions, as suggested by Lycus et al^30^. Plating on three different agar media resulted in 104 isolates. Of these, seven *Trinickia* and one *Dyella* strains were able to reduce N_2_O at both neutral and low pH, whereas one Paraburkholderia strain was only able to reduce N_2_O at acidic pH (Supplementary Results chapter 2.4).

### Validation of functional NosZ under low pH conditions

To verify that N_2_O reduction at low pH originates from NosZ assembled under acidic conditions, meticulous precautions were taken to ensure that the inoculum did not contain NosZ that had been pre-synthesized at circumneutral pH (Supplementary Results chapter 2.5). The inocula were pre-grown in low-pH, buffered medium for >23 generations. Continuous and vigorous stirring maintained strictly oxic conditions and prevented cell aggregation; this was essential, since aggregates may develop higher local pH than the bulk medium. Using such acidic inocula, functional NosZ production at low pH was demonstrated for the isolated strain *Trinickia* sp. Z7 incubated in media covering a wide pH range (Fig. 1c, Extended Data Fig. 3a and 4, Supplementary Fig. 10). Two other isolates, *Dyella* sp. L6 (Extended Data Fig. 3b) and *Paraburkholderia* sp. L5 (Extended Data Fig. 5), were also able to reduce N_2_O under strict acidic pH conditions. In addition, we tested five strains retrieved from culture collections (Supplementary Table 2), selected because they carry two additional genes *nosP* and *nosQ* (Fig 2) in their *nos* gene clusters. All these strains were evidently able to synthesize functional NosZ at low pH (Extended Data Fig. 5 & 6).

**Fig. 2.**
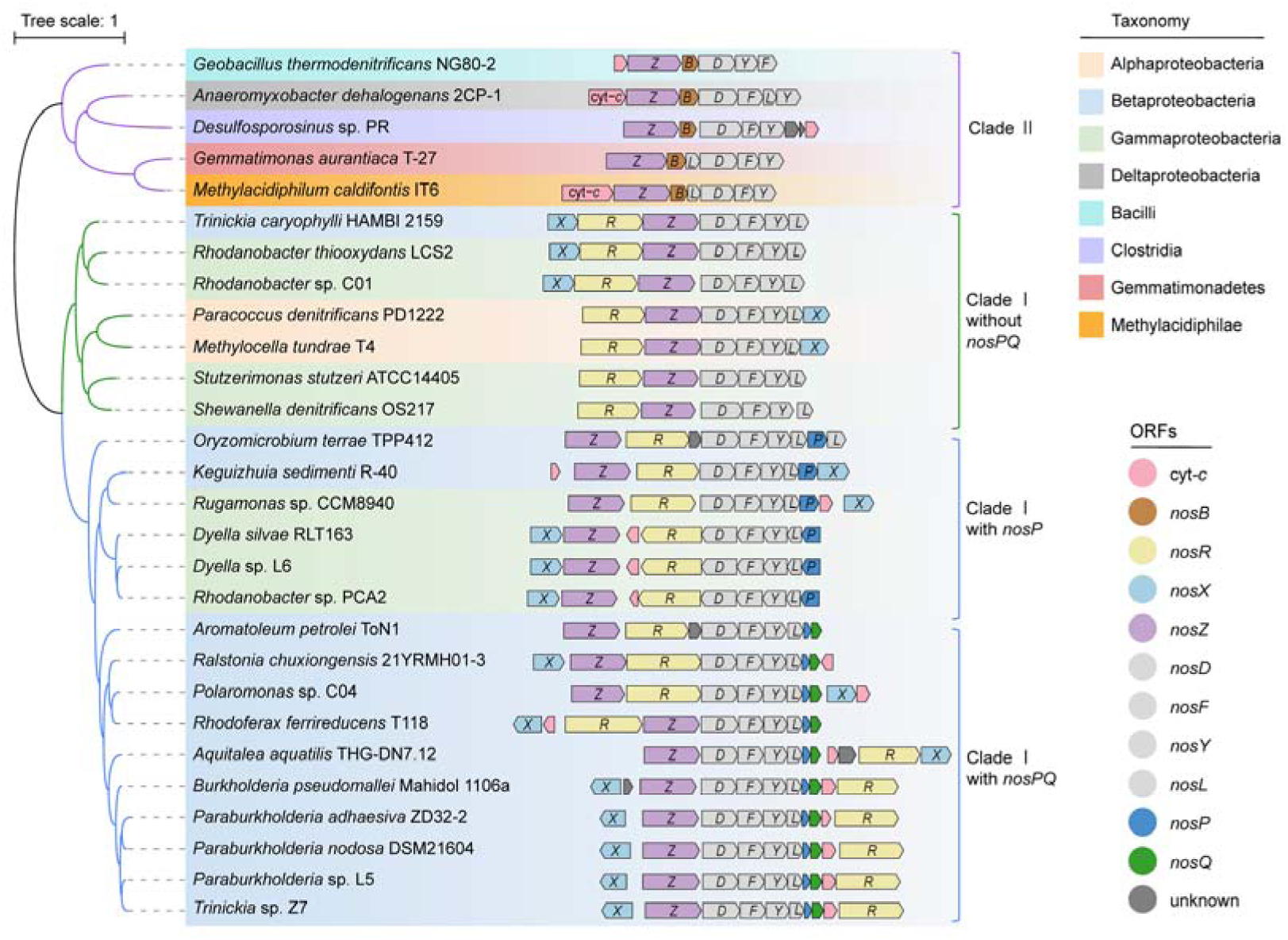
Maximum-likelihood phylogenetic tree and *nos* gene cluster organization in bacteria carrying clade I and clade II nitrous oxide reductases. The reference genomes were obtained from NCBI, including the recently reported acid-tolerant N_2_O reducers and some of the *nosPQ*-containing genomes. Putative *nos*-cluster genes were predicted using the genome annotation approaches detailed in the Supplementary Methods section. The phylogenetic tree was constructed with IQ-TREE using aligned NosZ protein sequences from each genome and visualized on iTOL. The colored arrows in the *nos* gene clusters represent different functional genes and indicate orientation and approximate length.

### The *nos* gene clusters in acid-tolerant N_2_O reducers

Genome analysis showed that the isolates *Trinickia* sp. Z7 and *Paraburkholderia* sp. L5 both carry the gene of *nosZ* clade I ^37^ with typical *nos* gene clusters (Fig. 2), including *nosX* (two copies in *Trinickia* sp. Z7), *nosZ*, *nosD*, *nosF*, *nosY*, *nosL*, and *nosR*. Interestingly, three additional genes were localized between *nosL* and *nosR* in the *nos* gene clusters of both organisms. These include a *c*-type cytochrome gene and two previously uncharacterized genes, designated as *nosP* and *nosQ* in this study (Supplementary Results chapter 2.6). Multiple sequence alignments and structure predictions of Nos proteins from *Trinickia* and *Paraburkholderia* revealed key structural motifs for metal coordination, such as the seven histidine ligands for Cu_Z_ center in NosZ, the Cu^+^ binding sites in NosD and NosL^19^. The *nos* gene cluster of the isolate *Dyella* sp. L6 also harbors *nosP* and *nosQ*, but in this strain they are fused into a single open reading frame (ORF), which we here designate as *nosP*, located downstream of *nosL*. Transcriptome sequencing of cultures incubated at pH 4.6 (gas kinetics shown in Extended Data Fig. 3a) confirmed that *Trinickia* sp. Z7 transcribed most *nos* genes, including *nosP*, during preculturing under oxic, acidic conditions, except for *nosQ* and *nosR* (Extended Data Fig. 7). When shifting to N_2_O reduction, transcription of all *nos* genes, including *nosP* and *nosQ*, increased significantly, except for the *nosX-1 and nosX-2* genes.

A genome analysis of the currently reported acid-tolerant N_2_O-reducing bacteria, *Desulfosporosinus* sp. PR^28^, *Methylocella tundrae* T4 and *Methylacidiphilum caldifontis* IT6 ^29^, *Rhodanobacter* sp. C01^30^, showed that they all lack *nosP* and *nosQ*, and have significantly different *nos* gene cluster arrangements and NosZ protein sequences compared to the *nosPQ* containing bacteria identified in this study (Fig. 2). Based on NosZ protein sequences and their *nos* gene cluster arrangements, the analyzed *nos* gene clusters were classified into three distinct groups: *nosZ* clade II, *nosZ* clade I without *nosPQ* and *nosZ* clade I with *nosPQ*. The *nosPQ* genes were found exclusively in *nosZ* clade I gene clusters, mainly affiliated with the Beta- and Gamma-proteobacteria.

### NosPQ are essential for N_2_O reduction at low pH

To investigate the role of NosPQ in N_2_O reduction at low pH, a *Trinickia* sp. Z7 *nosPQ* deletion strain was generated by a double crossover recombination. Gas kinetics demonstrated that this *Trinickia* Δ*nosPQ*::*Gm* mutant retained the N_2_O reduction activity at pH 6.8, but lost it at pH 4.8 (Fig. 3a).

**Fig. 3.**
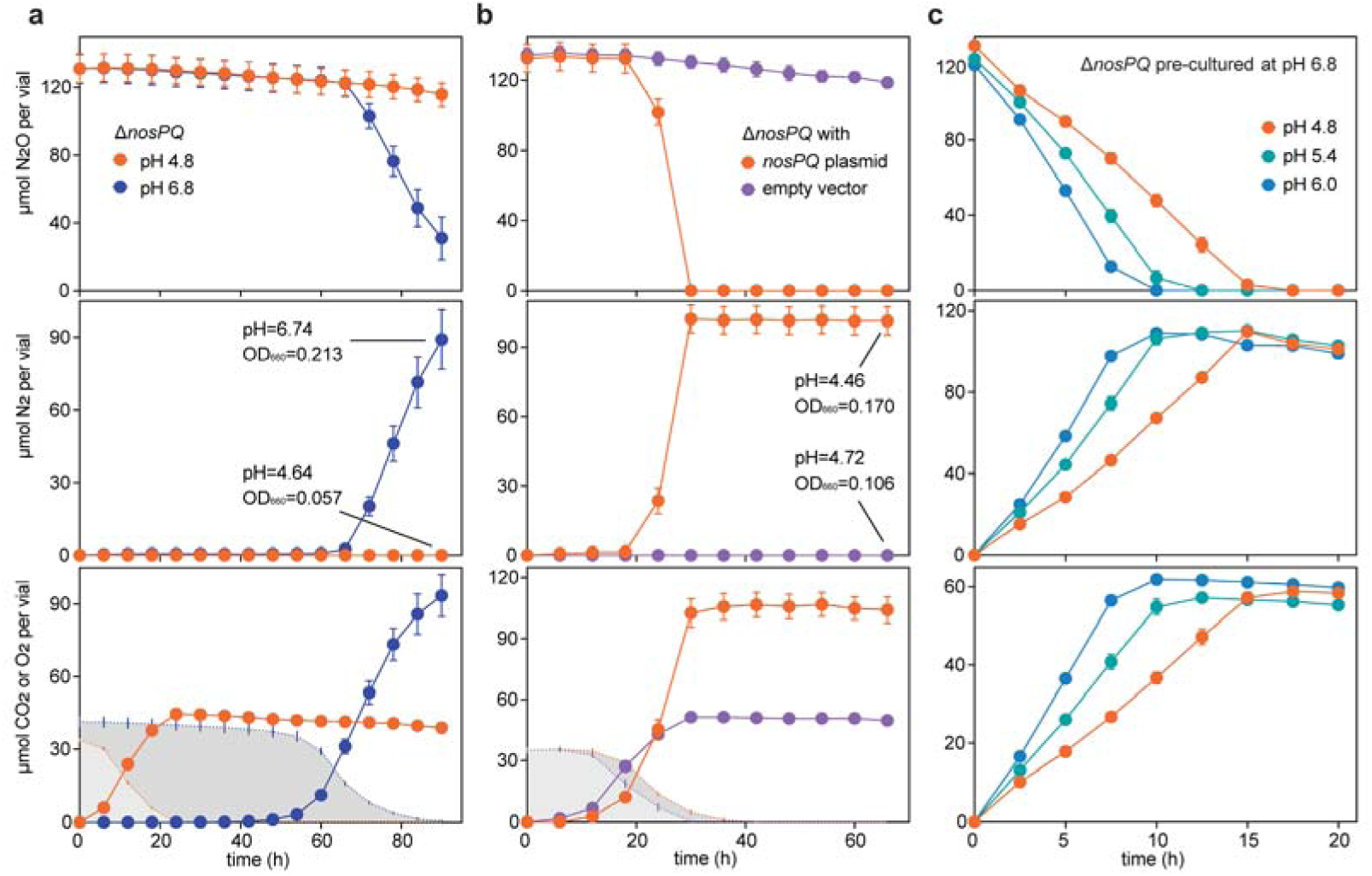
The role of NosPQ in mediating NDO-reducing activity under low-pH conditions, verified by gene manipulation of *Trinickia* sp. Z7. (**a)** N_2_O reduction by *Trinickia* Δ*nosPQ*::*Gm* mutant at pH 4.8 and 6.8. The cultures were incubated in vials containing 30 mL of Sistrom’s medium supplemented with 25 μg/mL gentamycin. 20 mM MES buffer was used to maintain the pH of medium. Prior to the incubation, the vials were made anoxic by replacing the headspace with He, after which 1 mL of O_2_ was added in the headspace to ensure cell growth. (**b**) N_2_O reduction by the *Trinickia* Δ*nosPQ*::*Gm* mutant complemented with the *nosP* and *nosQ* carried on a pBBR1MCS-2 plasmid. *nosP* and *nosQ* genes were cloned into a pBBR1MCS-2 plasmid. The empty vector was used as control. The inocula for the experiments shown in panels a and b were precultured for >15 generations to deplete the cells of any N_2_O reductase that may have been synthesized earlier under non-acidic conditions. The shaded areas show O_2_. (**c**) N_2_O reduction by *Trinickia* Δ*nosPQ*::*Gm* at various pHs. The cultures were first raised in pH 6.8 medium to allow the cells to produce functional N_2_O reductases, then transferred to anoxic media in which pH was set to 4.8, 5.4 or 6.0.

**Fig. 4.**
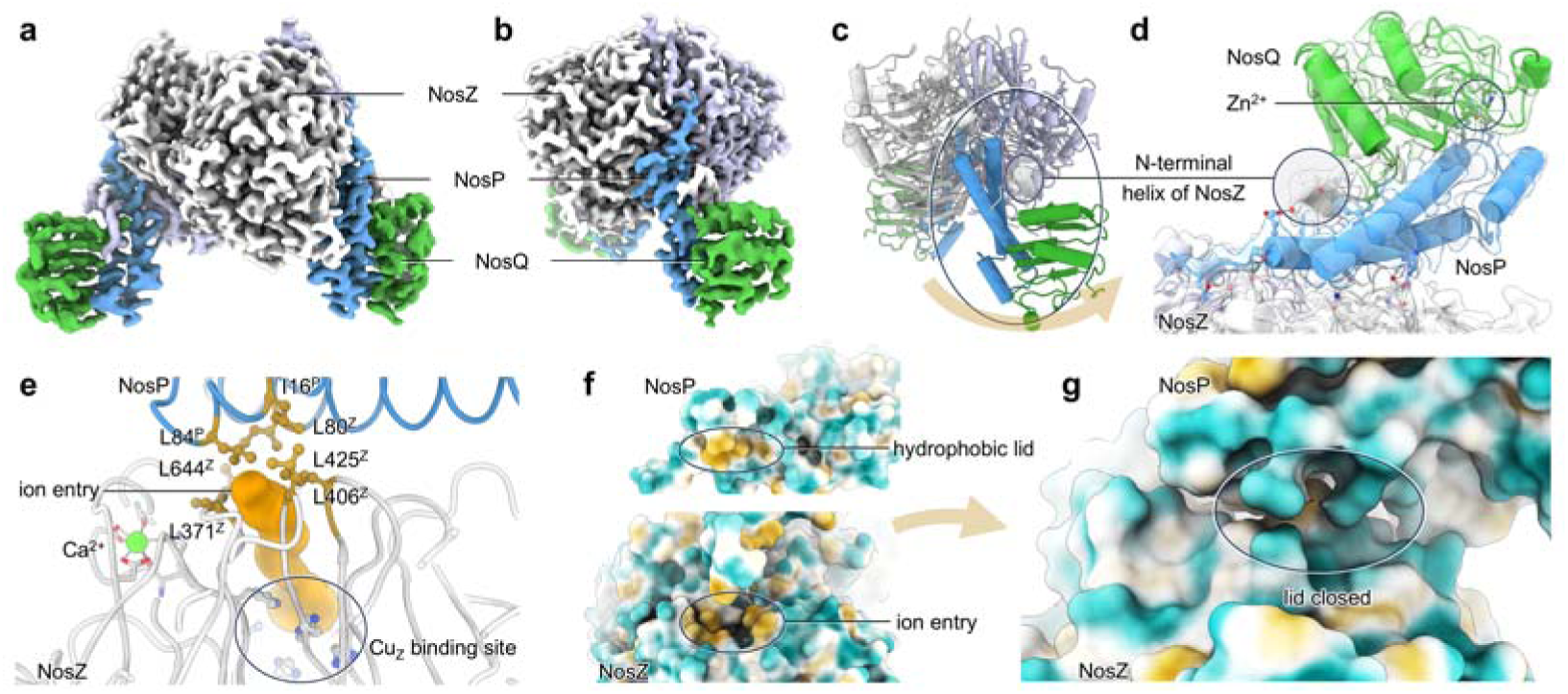
The cryo-EM structure of the NosZPQ complex. Cryo-EM maps of *Trinickia* NosZPQ at 2.4 Å resolution in front (**a**) and side view (**b**). NosZ is shown in grey, NosP in blue and NosQ in green. (**c**) Overall architecture of the NosZ_2_P_2_Q_2_ complex. (**d**) The interactions among NosZ, NosP and NosQ. A zinc ion is bound between NosP and NosQ, stabilizing the NosPQ dimer, and NosP directly binds to the ion entry site of NosZ. The N-terminus of NosZ further interacts with both NosP and NosQ to stabilize the ternary complex. (**e**) The hydrophobic residues at the ion entry site. (**f**) Surface hydrophobicity of NosZ and NosP. (**g**) The ion entry is closed after complex formation.

Complementation experiments were performed using recombinant plasmids carrying *nosPQ* under different promoters (Fig. 3b, Supplementary Fig. 11a). As described above, the mutant strain *Trinickia* Δ*nosPQ*::*Gm*, which had lost the ability to reduce N_2_O at pH 4.8, regained this ability through overexpresses *nosPQ* by transforming constructed plasmid. In contrast, cells transformed with an empty plasmid failed to restoreN_2_O reduction at low pH. Collectively, these results demonstrate that NosPQ is essential for N_2_O reduction under acidic conditions. Next, we investigated whether strains are incapable of reducing N_2_O under acidic conditions could gain this ability by introducing a plasmid expressing *nosPQ*. While such transformation of *Diaphorobacter* sp. RS54 and *Achromobacter* sp. SF6 failed to establish significant N_2_O reduction at low pH, the transformed *Pseudomonas* sp. SH1 strain showed a slight but statistically significant N□O reduction at low pH (Extended Data Fig. 8). Thus, introducing *nosPQ* can apparently facilitate the synthesis of some functional NosZ under acidic conditions in certain N_2_O reducers that naturally lack these genes.

Taken together, these results clearly demonstrate the essential role of NosPQ in N_2_O reduction at low pH. However, a key question remains: does NosPQ function to maintain the activity of matured NosZ at low pH, or is it only involved in the maturation process itself? To address this, the *Trinickia* Δ*nosPQ*::*Gm* mutant was cultured at pH 6.8 under denitrifying conditions, allowing the cells to produce functional NosZ without the NosPQ complex (Supplementary Fig. 11b). The cells were then transferred to acidic media of pHs 4.8, 5.4 and 6.0, to test their N_2_O reducing activity. As shown in Fig. 3c, cells harbouring matured NosZ were able to reduce N_2_O continuously at a low pH of 4.8, although the rate was slightly slower than at pH 5.4 and 6.0, meaning that NosZ was functional at low pH despite the absence of NosPQ. This shows that NosPQ is not required for activity of NosZ at low pH if properly assembled at circumneutral pH. These observations are analogous to earlier studies of *Paracoccus denitrificans* as well as of complex soil bacteria communities^13,15^ which showed that low pH impairs the synthesis of functional NosZ, while NosZ synthesized at circumneutral pH is functional at low pH.

### Molecular basis for NosPQ-dependent protection of NosZ at low pH

Protein structure and complex formation predictions with AlphaFold3^38^ suggested that NosP and NosQ form a heterodimer that can interact with the NosZ complex, which could imply a role in NosZ maturation. We heterologously expressed NosPQ in *E. coli*, with a C-terminal Strep-tag II on NosQ. Coexpression with NosZ resulted in the isolation of a complex fraction containing NosZPQ *via* affinity and size-exclusion chromatography (Supplementary Fig. 12). Cryo-EM single particle analysis yielded a structure at 2.4 Å resolution that (Extended Data Fig. 9) contained two copies of each NosZ, NosP and NosQ, with a total mass of 198 kDa (Fig. 4a, 4b). The *Trinickia* NosZ forms a homodimer, like all known N_2_O reductases^39^, as the core component of the hexameric complex. On each side of the NosZ dimer, a NosPQ heterodimer binds to the surface (Fig. 4c, 4d), where it interacts with NosD in the NosZDFY complex to facilitate the assembly of the copper centers^19^. The NosZ reductase is a periplasmic enzyme, and both the assembly of its metal sites and the catalytic reduction occur in the periplasm^20^. Signal peptide predictions suggest that the *Trinickia* NosZ contains a Tat signal peptide, however, both NosP and NosQ lack signal peptide sequences. Thus, a plausible mechanism is that NosPQ and NosZ first assemble into a complex in the cytoplasm, where NosPQ shield ion access to the copper-binding site of NosZ, thereby protecting it; subsequently, the entire complex is co-translocated into the periplasmic space where the Cu insertion takes place.

The assembly of the Cu_A_ and Cu_Z_ sites require copper delivery from the Cu-chaperon NosL^40^, *via* NosD within the NosDFY complex^19,20^, and then on to NosZ. Previous studies have suggested that acidic pH interferes with the maturation of NosZ in many bacteria^13,14^, likely because the pH in the periplasm is similar to that outside the cell ^41^. As shown in Fig. 1c, *Trinickia* is able to overcome this and effectively reduce N_2_O at pH 4.5, indicating that the molecular interactions were not significantly affected. To find a possible clue to the mechanism behind this, we analyzed the NosZ ion binding sites. Consistent with all known NosZ structures, the Cu_A_ and Cu_Z_ coordination sites in *Trinickia* NosZ are composed of histidine residues that are prone to protonation at low pH, thereby losing the ability to coordinate copper ions. The Cu_Z_-binding site comprises seven histidine residues located within the NosZ dimer, approximately 15 Å from the protein surface. Analysis of ion access to the Cu_Z_ binding site using MOLE^42^ identified a tunnel with a minimum radius of 1.6 Å, sufficient space to deliver protons, possibly along a water chain (Fig. 4e). In our NosZPQ structure, this entry is blocked (Fig. 4g) by hydrophobic interaction (Fig. 4f) between NosZ and NosP, thus effectively preventing protonation of the histidine residues at acidic conditions. In this scenario, the protective complex of NosZPQ would further interact with the copper delivery system NosDFY to facilitate copper sites assembly.

### Widely distributed acid-tolerant N_2_O reducers

To explore the distribution of *nosPQ* genes, we used the NCBI Web BLAST tool to retrieve all protein sequences identified as NosP and NosQ homologues in the nr database. In total, 298 and 378 sequences producing significant alignments with NosP and NosQ, respectively, were successfully identified (Supplementary Table 5). Notably, 21 non-redundant protein sequences displayed significant alignments with both NosP and NosQ, showing structural resemblance to the longer NosP variant identified in *Dyella* sp. L6. Phylogenetic analysis demonstrated that these NosPQ sequences were predominantly associated with the Proteobacteria phylum, particularly within the Beta-proteobacteria, distributed across 42 distinct genera (Supplementary Fig. 13).

To examine the distribution of microorganisms harbouring *nosPQ* genes in soils, we searched for these genes in 535 soil metagenomes from 187 locations worldwide and detected *nosPQ* in 383 metagenomes spanning 163 sites (Fig. 5a, Supplementary Table 10 &11). We investigated the taxonomy of the *nosPQ* gene sequences and their relationship to soil pH (Fig. 5b), and clustered metagenomes based on their *nosPQ* sequences, generating three distinct clusters (Fig 5c, Supplementary Fig. 14 & 15). The mean soil pH values for Cluster 1, Cluster 2, and Cluster 3 were 6.46, 6.09, and 5.16 (Fig. 5d), respectively. Plotting the *nosPQ* gene abundances (as RPM values) of each metagenome against the soil pH revealed substantial overlaps between the clusters, both with respect to pH and *nosPQ* gene abundance. However, Cluster 3 exhibited a significantly (p<0.0001) higher average *nosPQ* gene abundance than the two others. The dominant microbial taxa specific to each cluster were identified via LEfSe analysis (Fig. 5e). In the more strongly acidic soil samples of Cluster 3, dominant genera included *Paraburkholderia*, *Polaromonas*, *Dyella*, *Ralstonia*, *Azoarcus*, *Trinickia* and *Burkholderia*, which were less abundant in the samples of other two clusters.

**Fig. 5.**
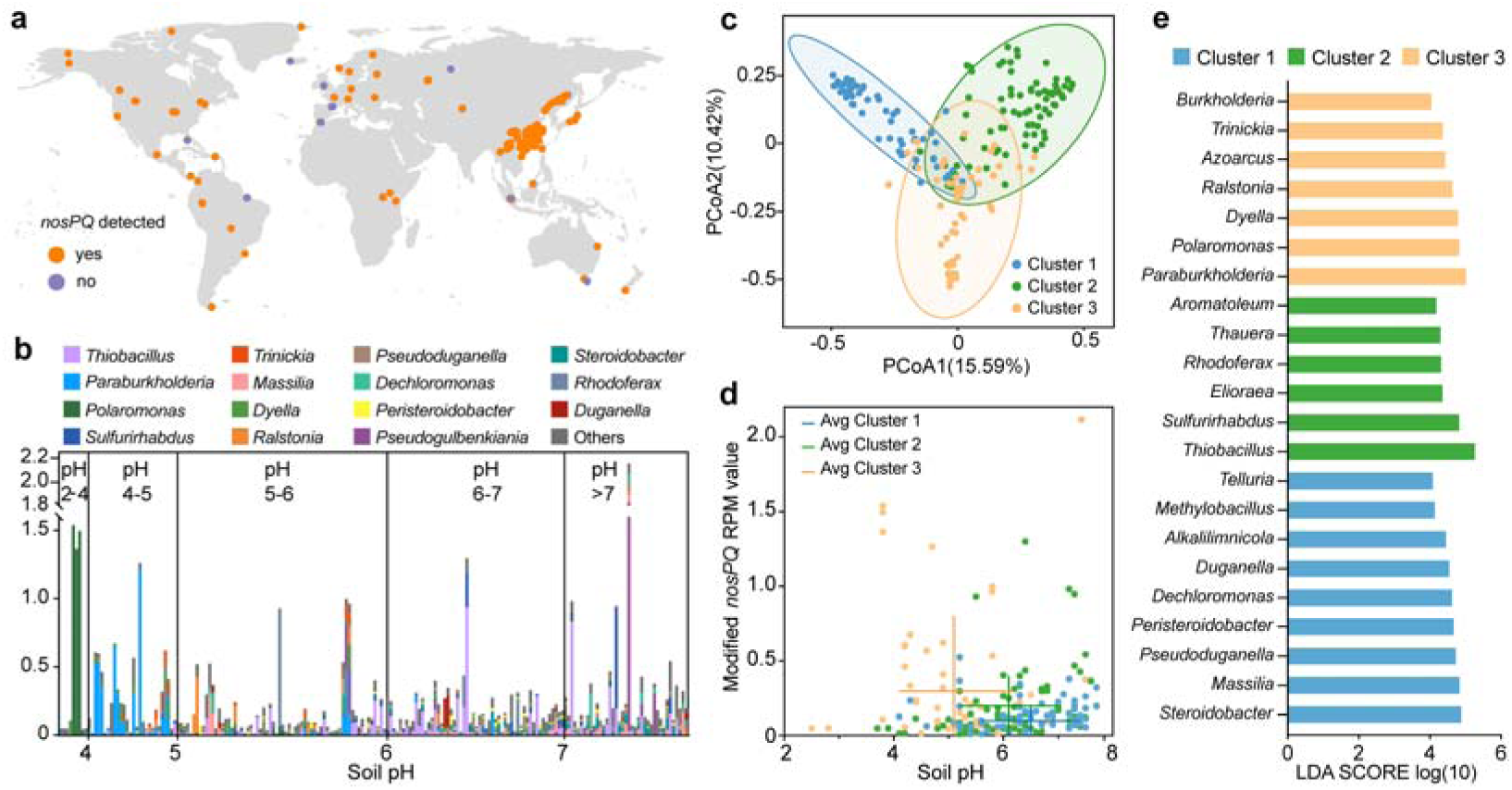
The global soil metagenome analysis of *nosPQ* genes. (**a**) Geographic origin of soil metagenomes analyzed for *nosPQ* genes. Each point represents one location except for some overlapped points. (**b**) The abundance and taxonomy of *nosPQ* genes in the metagenomes arranged according to soil pH. The vertical axis shows the modified RPM values, including the top 15 genera and ‘Others’, with the unclassified reads removed to enhance the visibility of the taxonomy, unclassified reads were removed. 204 samples with RPM values >0.1 and counts of total mapped reads ≥10 were retained. 6 samples were not annotated with classified taxa and were discarded, yielding 198 samples for subsequent analysis. (**c**) Principal coordinate analysis (PCoA) of the *nosPQ* gene-containing metagenomes. Each point represents a sample and the ordinations were calculated based on the Bray–Curtis distances of the *nosPQ* sequences. Samples were divided into three clusters according to the UPGMA analysis results. The PERMANOVA analysis shows p<0.0001. (**d**) *nosPQ* abundance (RPM) within the three clusters of each metagenome, plotted against soil pH. The crosses are average pH- and RPM-values for each cluster, +/- standard deviation. Cluster Ordinary one-way ANOVA, followed by a Tukey’s test for multiple group comparisons, showed that Cluster 3 had significantly lower pH and higher *nosPQ* RPM values than the two other clusters (**p<0.01). (**e**) Bar plot of the Linear Discriminant Analysis (LDA) scores of the significantly different taxa among three cluster as determined by LEfSe analysis. Only taxa with LDA score log (10) >4 are displayed.

Of note, the measured abundances of *nosPQ* genes in the metagenomes from all soils are low, even in most of the metagenomes from acidic soils. Recorded *nosZ* clade I abundances in other metagenome studies range from 5-10 RPM ^17,43,44^, which is 1-2 orders of magnitude higher than the *nosPQ* abundances recorded here. This comparison indicates that most bacteria with *nosZ* clade I lack *nosPQ*, even in acidic soils. The dominance of organisms unable to synthesize functional *nosZ* likely explains why liming acidic soils leads to an almost immediate enhancement of their N_2_O reduction capacity ^36^. One would expect that high soil pH would favor organisms with *nosZ* clade II over *nosZ* clade I, as observed by Frostegård et al ^17^, and that organisms with *nosPQ* would be favored by low soil pH. We investigated this by searching the metagenomes for the abundance of the genes in question. We found that the *nosZ* II/*nosZ* I abundance ratio (R) increased with increasing soil pH (linear regression, p<0.001, Extended Data Fig. 10) corroborating the conclusion by Frostegård et al ^17^, although the regression function only explain 7.9 % of the variation (r^2^=0.0785). However, we found no evidence for low pH favoring organisms with *nosPQ* (linear regression, not significant, Supplementary Fig. 16). The ratio between the abundance of *nosPQ* and *nosZ* clade I was basically unaffected by soil pH (linear regression, not significant, Supplementary Fig. 17). This would suggest that the selective advantage of being able to reduce N_2_O in acid soils is too marginal to allow such organisms to become dominant in acid soils, which could open an opportunity to combat N_2_O emission from acid soils by bioaugmentation of such organisms.

### Concluding remarks

Acidic environments pose significant challenges to microbial N_2_O reduction, and are recognized as hotspots for N_2_O emissions. Despite its importance, the precise mechanisms underlying the impaired N_2_O reduction at acidic pH are not understood, although strong evidence points to a post-transcriptional phenomenon^13,15,17^. Nevertheless, the capacity for N_2_O reduction in acidic soil has been frequently reported^17,25–27,43,45^, and also demonstrated for two acidic soils in this study: bacteria in these soils were evidently able to synthesize functional NosZ at low pH, a capability confirmed for three isolated strains. Further investigations of the molecular mechanisms using *Trinickia* sp. Z7 as a representative, revealed two novel proteins, encoded by genes designated as *nosP* and *nosQ*, that are crucial for the maturation of N_2_O reductase under acidic pH conditions, but not required for its function if assembled at circumneutral pH. Structural analysis of these novel proteins offered detailed insights into their role in the proper assembly of the Cu_Z_ center, a critical step in the maturation of N_2_O reductase under acidic conditions.

We searched for the *nosPQ* gene sequences in the NCBI nr database and in 535 published global soil metagenomes. These genes were identified in taxonomically diverse bacteria, mostly within the Proteobacteria, and showed a ubiquitous distribution across a wide range of soils. Further studies on the physiology and ecology of N_2_O-reducing bacteria in acidic soils are needed to better understand their functional role *in situ*. Such knowledge could facilitate the development of innovative strategies to curb N_2_O emissions from widely distributed acidified farmlands, thereby making a substantial contribution to the mitigation of global climate change.

## Methods

### Soil samples

The paddy soil was collected from a rice field in Ji’an city, Jiangxi Province, China (27°10′N, 114°25′E). This location lies within a typical red-soil hilly area in southern China, which is characterized by a subtropical monsoon climate. Double-cropping of paddy rice is the predominant agricultural practice in this region. Classified as Ultisols according to the USDA Soil Taxonomy^46^, the soil is a sandy loam, consisting of approximately 58 % sand, 31 % silt, and 11 % clay. The forest soil was collected from an adjacent evergreen coniferous forest dominated by pine trees. At each site, soil samples were randomly collected from the topsoil layer (0∼20 cm) across multiple plots in May of 2021. After transporting to the laboratory, the soils were air-dried, ground, then passed through a 2-mm sieve to remove stones and plant debris. The pH of each soil sample was measured in a soil-to-distilled water mixture at a ratio of 1:2.5 (w/v).

### Nitrous oxide reduction in two acidic soils

Thirty grams of processed paddy soil or forest soil was placed in separate 120 mL glass vials, respectively. The soil water content was adjusted to 30% of the dry soil weight using distilled water. All vials were incubated at 25□ under oxic conditions for one day, then crimp-sealed and made anoxic through repeated rounds of evacuations and helium flushing. Subsequently, 5 mL of N_2_O (approximately 404 μmol N) or 3 mg of NO ^-^ (approximately 48 μmol N) was added to each vial. The gas kinetics was monitored in real time using an robotic gas sampling system ^47^. Three replicates were set up for each treatment.

N_2_O reduction was also measured in paddy soil slurries, to minimize heterogeneity and effects of soil aggregation. For this, paddy soil was mixed with distilled water at a ratio of 1:5 (w/v). The mixture was sterilized at 121□ for 20 minutes. The supernatant obtained after centrifugation of the sterilized mixture was supplemented with glucose to a final concentration of 1 g/L and used as the soil extract□glucose (SEG) medium in subsequent experiments. One gram of paddy soil from a previous experiment, in which it had been incubated anoxically for 3 days in the presence of 5 mL of N_2_O, was mixed into 30 mL of SEG medium in a 120 mL serum vial. The headspace of the vials was replaced with helium supplemented with 5 mL of N_2_O, after which gas kinetics were monitored. Vials containing the same SEG medium, but without soil inoculation, served as abiotic controls. All vials were stirred vigorously (500 rpm) using a magnetic stir bar (2.5 cm×0.5 cm).

### Paddy soil incubations

Paddy soil (10 g per vial) was preincubated under oxic conditions at 25□ to activate the soil microorganisms (soil O), after which it was treated with different amendments. Group G received 2 mL of 10 mg/mL glucose to stimulate heterotrophic respiration; Group C received 2 mL of 0.01 g/mL calcium hydroxide suspension as liming treatment; and Group S received 2 mL of sterile distilled water as a control. Soil pH was measured immediately after the additions. All vials were then rendered anoxic by Helium flushing and injected with 5 mL of N□O. Gas kinetics were continuously monitored in three replicates, and parallel vials were destructively sampled in triplicates at 6 h, 18 h, and 42 h for molecular and physiochemical analyses. All collected samples were stored at −80 □ until further analysis. At the end of the 3-day incubation, the pH was measured in all samples. In an additional experiment, the N_2_O reduction of the intact paddy soil samples was compared to that of paddy soil slurries (1:2.5, w/v), prepared as described above.

### Meta-omics sequencing and bioinformatic analysis

To identify the microbes that could possibly be responsible for N_2_O reduction in acidic paddy soil during the incubations, metagenomic and metatranscriptomic sequencing was performed on selected soil samples. Samples for metagenomic sequencing were taken from soil O, which had been preincubated under oxic conditions without further incubation, and from Groups S, G and Group C after 42 h of incubation. Metatranscriptomic sequencing was used to identify active bacteria and functional genes in the S and G treatments incubated for 6, 18 and 42 h (S6/S18/S42, G6/G18/G42); from the limed soil incubated for 18 and 42 h (C18/C42); and from soil O. The procedures and data processing methods for soil metagenomic and metatranscriptomic sequencing are described in detail in the Supplementary Methods chapter 1.1 & 1.2. All statistical analyses were conducted using R script, including calculations of Shannon indices, Bray□Curtis distance metrics, permutational multivariate analysis of variance (PERMANOVA) tests, ordinary one-way analysis of variance (ANOVA).

A reads-based annotation method was employed to analyze the *nosZ* genes and transcripts via comparison against a custom dataset ^48^. Reads that had a matching region of >30 amino acids and an identity of >60% were considered matching. The output of *nosZ* reads was normalized to RPM (reads per million total reads) values to compare their abundances across different samples. To explore variations in *nosZ* reads abundances among various samples, the normalized abundances were summarized at the genus level. Co-occurrence network analysis was conducted to examine the response patterns of these N_2_O reducers to different amendments^49^. Only genera present in >20% of all samples were included in the network analysis, yielding 321 qualified genera. The co-occurrence among these genera was detected by the “psych” package in R (v4.3.2) with the cutoffs of Spearman’s rho>0.6 and FDR P<0.05. Both the visualization of the co-occurrence network and module classification were implemented in Gephi (v0.9.2) (https://gephi.org/).

### Isolation and identification of N_2_O reducers

Samples for isolation of bacteria were taken from the soil incubation experiments described above, including SEG medium inoculated with paddy soil, glucose-amended soil (G42), lime-treated soil (C42), and newly activated pine forest soil. Soil dilutions were spread on agar plates of SEG medium (pH 4.5) and 1/10 TSA medium (pH 6.2). For isolation of non-spore forming bacteria, beef extract-peptone medium (pH 6.2) was also used, while LB medium (pH 6.5 and 4.8) was used for isolation of spore forming bacteria too. The plates were incubated under oxic conditions (30 □) to target as wide a diversity of bacteria as possible^30^. For isolation of sporulating N_2_O reducing bacteria, sample dilutions were heated at 80 □ for 10 minutes prior to plating to inactivate vegetative cells.

The taxonomy of each isolate was determined by 16S rRNA gene sequencing. Presence of the *nosZ* gene was confirmed by PCR amplification using primers targeting *nosZ* clade I and II^50^, followed by sequencing of PCR product. The N_2_O reduction capabilities of *nosZ*-containing isolates at acidic and circumneutral pH were verified in unbuffered media corresponding to their initial isolation conditions. Briefly, 300 μL of bacterial culture was inoculated into 30 mL of the respective medium with 5 mL of N_2_O in the headspace. N_2_O reduction and N_2_ production were monitored under continuous stirring (500 rpm) during anoxic incubation. Verified N_2_O reducers were whole genome sequenced (details described in Supplementary Methods chapter 1.3).

### Verification of N_2_O reduction at low pH

Cultures of *Trinickia* sp. Z7, which was the primary strain studied under various conditions, were raised from frozen stocks and plated on 1/10 TSA (pH 6.2). Single colonies were then transferred to 10 mL tubes containing 5 mL of 1/10 strength TSB medium. After 24 h of incubation in a shaker (150 rpm, 25□), 1 mL of the bacterial culture was inoculated into a 250 mL flask containing 100 mL of acidic 1/10 TSB medium (pH 4.6) containing phosphate buffer (20 mM H_3_PO_4_ and 20 mM Na_3_PO_4_) to mitigate pH increase. The flasks were covered with gauze and newspaper, and shaken continuously (150 rpm) to maintain cell dispersion during growth. After 24 h of incubation, 1 mL of the bacterial culture was transferred to a second flask containing the same acidic medium. Following three successive transfers, the bacterial culture was then used for inoculation.

For monitoring of N_2_O reduction, 300 μL of the prepared acidic bacterial culture was inoculated into 120 mL vials containing 30 mL of buffered 1/10 TSB media to an initial OD_660_ of approximately 0.001. The pH was set to 4.2, 4.6, 5.0, 5.5, 6.0 and 6.5. After purging with helium, all sealed 120 mL vials were supplemented with 3 mL (corresponding to approximately 120 µmol) of N_2_O, then incubated at 25□ with vigorous stirring (500 rpm) to prevent cell aggregation. At the end of the incubation period, the pH of each vial was measured. To continuously monitor the pH changes and N_2_O reduction dynamics during anoxic incubation, the experiments using pH 5.5 and pH 4.6 media were repeated with multiple replicates in an additional test. Gas kinetics were monitored in three replicates, while the remaining vials were destructively sampled at different timepoints to measure pH and OD. Gene expressions of the cultures in the pH 4.6 medium was assessed by transcriptome profiling (Supplementary Methods chapter 1.4) at three time points: 0 h (i.e. the pre-cultures after the third transfer), 24 h and 36 h after N_2_O addition. The Gammaproteobacteria isolate *Dyella* sp. L6 was first pre-cultured in acidic R2A medium (pH 5.3, supplemented with 20 mM MES buffer) following the same procedure as described above. After three serial transfers (16 generations), the cells were inoculated into 120 mL vials containing 30 mL of acidic R2A medium (pH 5.3 and 4.8) with 3 mL of N_2_O and 1 mL of O_2_ in the headspace.

Since 1/10 TSB and R2A media contain amino acids, microbial metabolism of these substrates may result in an increase of pH in the medium through consumption of acids and production of ammonia. To ensure that N□O reduction proceeded under stable acidic conditions, an amino acid-free, non-ammonium-generating medium (modified Sistrom’s medium) was additionally employed to verify N□O reduction under strict low-pH conditions (Supplementary Table 1). After one day of incubation at 28□ (150 rpm) in 1/10 TSB medium, the *Trinickia* sp. Z7 cells were washed three times with acidic Sistrom’s medium (pH 4.2, supplemented with 20 mM MES buffer). 300 μL of the resuspended cells were inoculated into 30 mL of Sistrom’s medium (pH 4.2) in 120 mL glass vials wrapped with aluminum foil. The vials were incubated at 25□ with vigorous stirring (500 rpm) using a magnetic stir bar to ensure complete cell dispersion. After three rounds of 24-hour incubation with serial transfers (1%), the bacterial cells were ready for use as inoculum and were observed via a transmission electron microscope (Talos L120C G2, Thermofisher) after staining with 1% (w/v) phosphotungstic acid.

The N_2_O reduction assay was conducted following the above-mentioned procedure. 20 mM of MES buffer was used to maintain the medium pH if mentioned. Firstly, N_2_O reduction was tested in unbuffered Sistrom’s medium (pH 4.5) to confirm the N_2_O reduction ability of *Trinickia* sp. Z7 at acidic pH. Experiments were then conducted across a wide pH range (buffered pH at 4.2, 4.8, 5.5, 6.2, 6.8, and 7.1). In addition, a parallel set of vials were supplemented with O_2_ in the headspace to monitor N_2_O reduction with the presence of O_2_. 1.5 mL, 3 mL or 18.9 mL (21%) of O_2_ was injected into the headspace of vials containing Sistrom’s medium at varying pH values. In addition to *Trinickia*, several other bacteria harboring *nosP* and *nosQ* genes—obtained from different microbial culture collections—were tested for N_2_O reduction activity under both acidic and circumneutral pH conditions in Sistrom’s medium (Supplementary Methods chapter 1.5).

### Genetic manipulation of *Trinickia* sp. Z7 and function assay

To explore the potential functions of the newly identified genes, a *nosPQ* double-deletion mutant of *Trinickia* sp. Z7, designated as Δ*nosPQ*::*Gm*, was constructed by inserting a gentamicin resistance cassette via double-crossover homologous recombination. The experimental details are described in Supplementary Methods chapter 1.6. To verify the N_2_O reduction by the *Trinickia* Δ*nosPQ*::Gm mutant, its cells were first cultured in Sistrom’s medium supplemented with 25 μg/mL gentamycin (pH 4.2, buffered with 20 mM MES), then serially transferred three times following the methods described above. Subsequently, 300 μL of the pre-cultured cells were inoculated into 30 mL of media (pH 4.8 and 6.8, buffed with 20 mM MES, 25 μg/mL gentamycin) with 3 mL N_2_O and 1 mL O_2_ in the headspace.

A complementation assay was performed in the *Trinickia* Δ*nosPQ*::Gm mutant to test the role of these two novel genes for N_2_O reduction under low-pH condition, (Supplementary Methods chapter 1.7). Specifically, 3 mL of N_2_O and 1 mL of O_2_ were added to the headspace, and the cultures of plasmid transformed cells were incubated under the same conditions as described above to monitor N_2_O reduction. In addition to the *Trinickia* Δ*nosPQ*::Gm mutant, the reconstructed expression plasmids harbouring the *nosP* and *nosQ* genes were electroporated into several other bacterial strains that contain *nosZ* but lack these two genes, including *Diaphorobacter* sp. RS54, *Achromobacter* sp. SF6, and *Pseudomonas* sp. SH1. These plasmid-transformed cells were then tested for N_2_O reduction ability in acidic 1/5 TSB medium (pH 5.3, buffered with 20 mM MES).

In a subsequent experiment, the *Trinickia* Δ*nosPQ*::*Gm* mutant was first grown in 30 mL of pH 6.8 medium to synthesize functional N_2_O reductase at circumneutral pH. After depleting the N_2_O, the bacterial culture was centrifuged and the liquid was removed. Subsequently, the cell pellets were resuspended in 30 mL of acidic media with different pH values (4.8, 5.4 and 6.0). Then, 3 mL of N_2_O was added to each vial, and gas kinetics was monitored immediately.

### Recombinant production and purification of the NosZPQ complex

The *nosPQ* and *nosZ* genes were amplified from the genomic DNA of *Trinickia* sp. Z7 and cloned into expression vectors to generate pET22S-*nosPQ*(Tr) and pET30-*nosZ*(Tr), respectively. The expressed NosQ will carry a C-terminal Strep-tag, while NosZ will have a C-terminal His-tag. After co-transformation of *E. coli* BL21(DE3) with the two plasmids, a single colony was cultivated in 100 mL LB medium containing ampicillin (100 μg/mL) and kanamycin (50 μg/mL) at 37°C and grown overnight. The preculture was transferred and grown in 4 L flasks until OD_600_ reached 0.6-0.8. Then 1 mM isopropyl β-D-thiogalactopyranoside (IPTG) was added to induce gene expression, and the culture was further incubated at 18°C for 18 h. The cells were harvested by centrifugation at 4°C (5,000 rpm, 10 min) and the pellet were resuspended in lysis buffer (100 mM Tris, pH 8.0, 150 mM NaCl, 10% glycerol), and lysed by French press. Cell debris was then removed by centrifugation at 4°C (15,000 rpm, 40 min). The protein was bound to 5 mL of Strep-Tactin 4FF Flow column (SMART, Suzhou, China) equilibrated with binding buffer (100 mM Tris pH 8.0, 150 mM NaCl, 10 % glycerol) using AKTA system (Cytiva), and eluted with 50 mL of elution buffer (100 mM Tris, pH 8.0, 150 mM NaCl, 5 mM desthiobiotin, 10 % glycerol). The purified protein was analyzed by SDS-PAGE. Target complex fractions were concentrated and further purified with size-exclusion chromatography (Superdex S200, 16/600, Cytiva) in SEC buffer (20 mM Tris, pH 8.0, 150 mM NaCl). Finally, the selected fractions containing the complexes were concentrated, aliquoted and stored at −80°C.

### Cryo-EM data collection, processing and model building

The ternary complex of NosZPQ was diluted to 10 mg/mL and mixed with 0.25% CHAPSO. Cryo-EM grids were prepared using a Vitrobot Mark IV (Thermo Fisher Scientific) as follows: 3 µL of the protein sample was applied to glow-discharged Quantifoil Cu R1.2/1.3 300-mesh grids, waited for 10 s and blotted for 5 s with filter paper, and plunge-frozen in liquid ethane cooled by liquid nitrogen. Data collection was performed on a 300 kV Krios G4 cryo-TEM (Thermo Fisher Scientific) equipped with a Falcon 4i direct electron detector. Images were acquired using a 10 eV slit width on the energy filter, with a defocus range of −1.0 to −2.0 µm and a total dose of 50 e□/Å² in electron-event representation (EER) format.

The micrographs were processed in cryoSPARC v4.6 ^51^ (Extended Data Fig. 10). The EER data were imported (40 fractions), motion corrected and contrast transfer function (CTF) estimated. Initial particle picking employed a blob picker, followed by extraction (3×downsampling) and 2D classification. Selected particle subsets underwent *ab-initio* reconstruction to generate reference volumes for heterogeneous refinement. Concurrently, representative 2D classes were used for template-based picking, with subsequent particles processed through two rounds of heterogeneous refinement. The particles of the best 3D classes were combined, and re-extracted without downsampling. An additional round of hetero-refinement was performed to further remove bad particles, and the best class was subjected to non-uniform refinement^52^. At this point, the AlphaFold model could be well fitted into the map. However, the density for NosPQ and the core-region of NosZ were relatively weak likely due to the dynamic nature. Multiple rounds of 3D classification with focused mask (Extended Data Fig. 10) were performed to generate a map in which all the components were well resolved. The final particle sets were subjected to non-uniform refinement with minimize over per-particle scale, optimize per-particle defocus and per-group CTF params switched on. The resulting map for NosZPQ was refined to a resolution of 2.40 Å.

The AlphaFold^53,54^ models of NosZ_2_P_2_Q_2_ were used as starting models for building. The model was fitted into the density map using UCSF ChimeraX^55^, followed by iterative refinement in COOT^56^ and real-space refined with PHENIX^57^. Structure validation was performed using MolProbity^58^. Data collection and refinement statistics are summarized in Supplementary Table 4. Figures were generated using PyMOL (Schrödinger LLC) or UCSF ChimeraX^55^.

### Diversity profiling of potential acid-tolerant N_2_O reducers

To explore the taxonomic diversity of *nosPQ* genes, the protein sequences of NosP and NosQ from *Trinickia* sp. Z7 were searched against the NCBI non-redundant protein sequences (nr) database (update date: 25-11-2024) through Web BLAST with an expected threshold set at 0.00001, respectively. The matched sequences, along with their respective accession numbers and taxonomic classifications, were downloaded for further analysis.

All the downloaded non-redundant NosP and NosQ protein sequences, along with the sequences derived from the isolates in this study (*Trinickia* sp. Z7, *Paraburkholderia* sp. L5, *Dyella* sp. L6), were made into a custom “NosPQ “ protein dataset, consisting of 660 protein sequences (Supplementary Table 5). A total of 600 soil metagenomes data were collected, with their corresponding pH values and geological information recorded (Supplementary Table 6). Fastp (v0.22.0) ^59^ was used for quality control to get clean reads. Samples with cleans reads >10 million were retained for subsequent analysis, yielding 535 samples for subsequent analyses. Metagenomic reads were aligned against the custom NosPQ dataset using Diamond (v2.1.11) ^60^ with an e-value cutoff of 0.00001. Mapped reads, which had a sequence identity of >60% and bit score >50, were considered matching and then extracted by Seqtk (v1.4) (https://github.com/lh3/seqtk). The resulting reads were normalized to RPM (Reads Per Million mapped reads) values and annotated by Kaiju (v1.10.1) ^61^ for taxonomic classification. In the following analyses, to ensure the robustness of results, only those samples in which the mapped *NosPQ* genes had RPM values >0.1 and total read counts ≥10, were retained. Additionally, hits annotated as unclassified *NosPQ* genes were removed from all samples. These filtering steps yielded 198 samples for downstream analysis. For the *nosZ* gene annotation, clean reads were aligned to a custom data set^44^, classifying them into NosZ clade I and NosZ clade II^43^. The computations were run on Siyuan-1 cluster supported by HPC Center at Shanghai Jiao Tong University.

## Data availability

All the sequencing data and genomes have been deposited in the NCBI Sequence Read Archive (SRA) database under the bioproject accession number PRJNA980409 (release date 2025-12-31). The atomic coordinates and cryo-EM map have been deposited in the Protein Data Bank (PDB) and the Electron Microscopy Data Bank (EMDB) with accession numbers 9X7X and EMD-66648.

## Acknowledgements

This work was supported was supported by the National Key Research and Development Program of China (2024YFB4105700), the National Natural Science Foundation of China (42577128, 31971526 and 32570137), the National Key Research and Development Program of China (2017YFD0200102), and the Research Council of Norway (projects No. 325770 awarded to Å.F. and No. 344289 awarded to LRB). The authors thank Suzhen Wang for the soil collection; Ningyi Zhou and Chaofan Yin at Shanghai Jiao Tong University for providing plasmids; Louise Sennett and James Shapleigh for providing sequences for the curated database used to extract NosZ clade I/II abundances from the metagenomes; Bo Chen, Qingwei Liu and Hanqing Zhao at Shanghai Jiao Tong University for data processing; Gang Fu at the structural biology core facility of Shenzhen Medical Academy of Research and Translation for his support during cryo-EM data collection; the Core Facility and Service Center of School of Life Sciences and Biotechnology at Shanghai Jiao Tong University for collecting gas kinetics data.

## Author contributions

X.W, X.Z, L.R.B., Å.F. and L.Z. designed the experiments. X.W, L.M., S.Y. and X.P. conducted the bacteria isolation and phenotyping experiments. X.W., H.L. and C.Y. performed the genetic manipulation for mutant construction and complementation. X.W., Y.Z., B.X., C.L., M.L. J.W. and M.Z. processed Omics data. J.Z. and L.Z. performed the protein complex purification and structural data collection. L.Z. processed the cryo-EM data and built the structural model. X.W., X.Z, L.Z., L.R.B. and Å. F. wrote the manuscript.

## Corresponding authors

Correspondence to Xiaojun Zhang, Lin Zhang, Åsa Frostegård or Lars R Bakken.

## Additional information Competing interest declaration

All authors declare no competing interests.

## Extended Data

**Extended Data Fig. 1.**
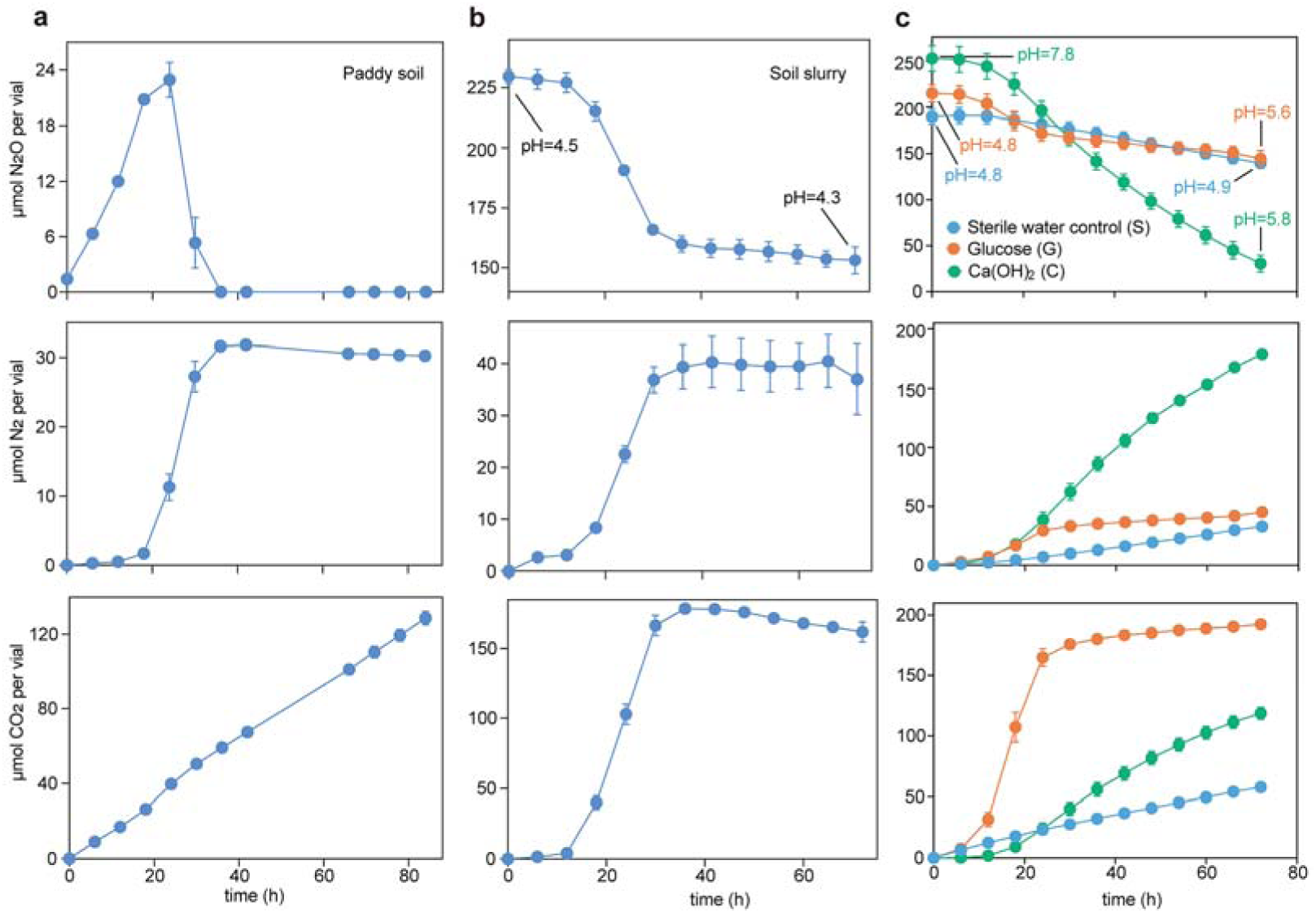
Comparison of N_2_O reduction in intact soil vs soil slurry of the acidic paddy soil. **(a)** 30 g (wet weight; ww) of intact paddy soil supplemented with 48 μmol NO_3_^-^. **(b)** 30 mL of soil slurry prepared as follows: 1 g of N_2_O-enriched paddy from the incubation shown in Figure 1b (main paper) was added to 30 mL of soil extract-glucose (SEG) medium (pH 4.5). The slurry was stirred using a magnetic bar (500 rpm) during the entire incubation. **(c)** 10 g of intact paddy soil supplemented with glucose (G): 2 mL of a 10 mg/mL glucose solution; Ca(OH)_2_ (C): 2 mL of 135 mM Ca(OH)_2_ solution; or sterile water (S): 2 mL of sterile distilled water. Gas kinetics were monitored during anoxic incubation of paddy soil (intact soil or soil slurry) in gas tight vials containing 5 mL of N_2_O in the headspace. pH was measured at the beginning and end of the incubation period.

**Extended Data Fig. 2.**
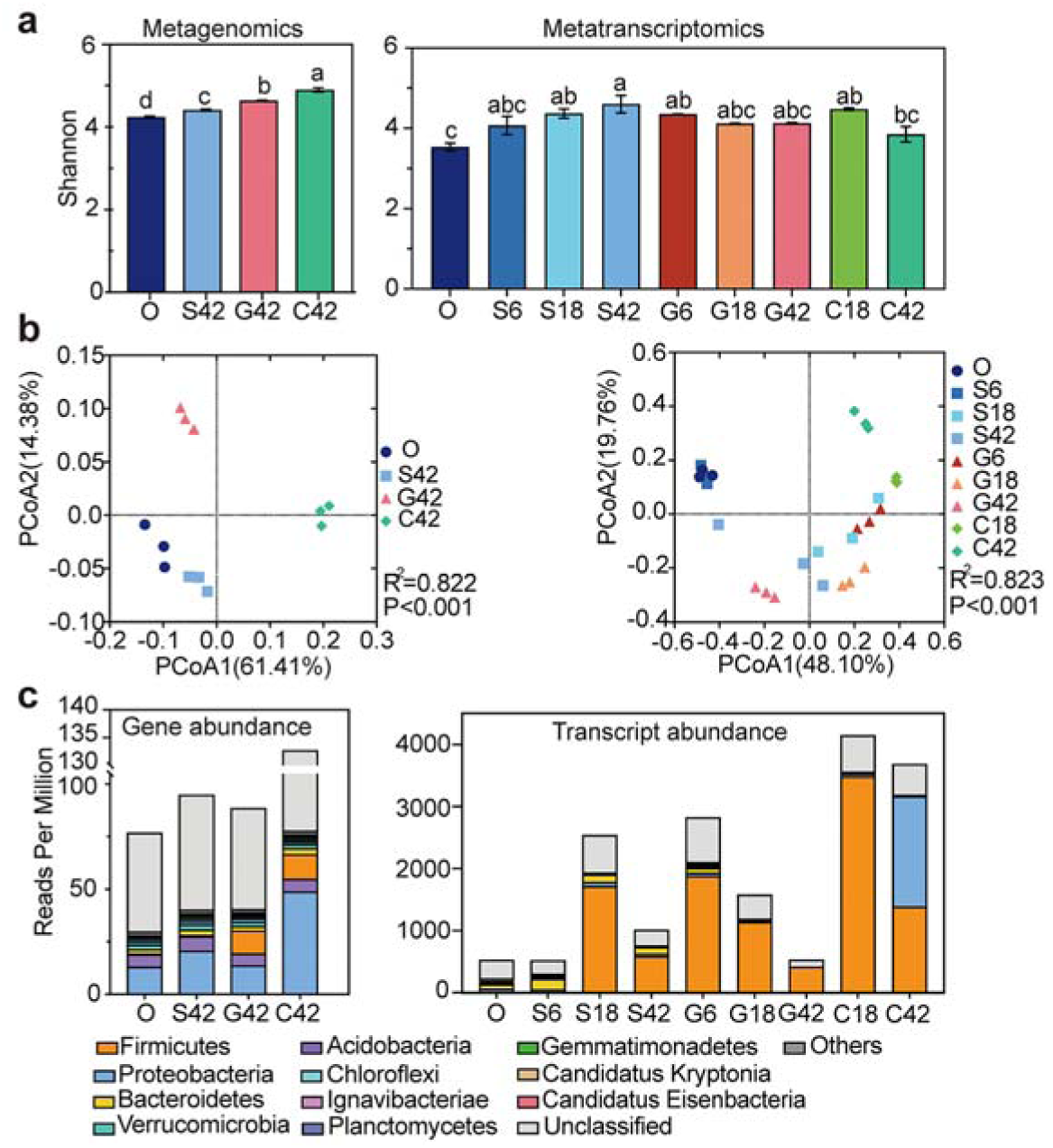
Shifts of the *nosZ*-containing microbial community in the paddy soil during N_2_O reduction in response to glucose addition and liming. Soil samples (10 g, ww) were exposed to one of three pretreatments and then incubated in sealed, anoxic vials containing helium and approximately ∼204 μmol N_2_O in headspace. O: Oxic soil sampled only at time 0, serving as the original soil control. S: 2 mL of sterile distilled water added as control treatment. G: Addition of 2 mL of a 10 mg/mL glucose solution. C: Addition of 2 mL of a 135 mM Ca(OH)_2_ solution. Soil samples were collected from treatments S, G and C at 42 h after N_2_O addition for metagenomics analysis and at 6, 18, 42 hours after N_2_O addition for metatransctiptomics analysis. The four-group figures (left) show metagenomics results and the nine-group figures (right) show metatranscriptomics results. (**a**) Alpha diversities measured by Shannon index with ordinary one-way analysis of variance (ANOVA) followed by a Fisher’s Least Significant Difference (LSD) test. (**b**) Principal-coordinate analysis (PCoA) along with corresponding PERMANOVA comparisons (9,999 permutations) for each sample. The ordinations were calculated based on the BraylJCurtis distances. (**c**) Relative abundances of the top 10 *nosZ*-containing phyla.

**Extended Data Fig. 3.**
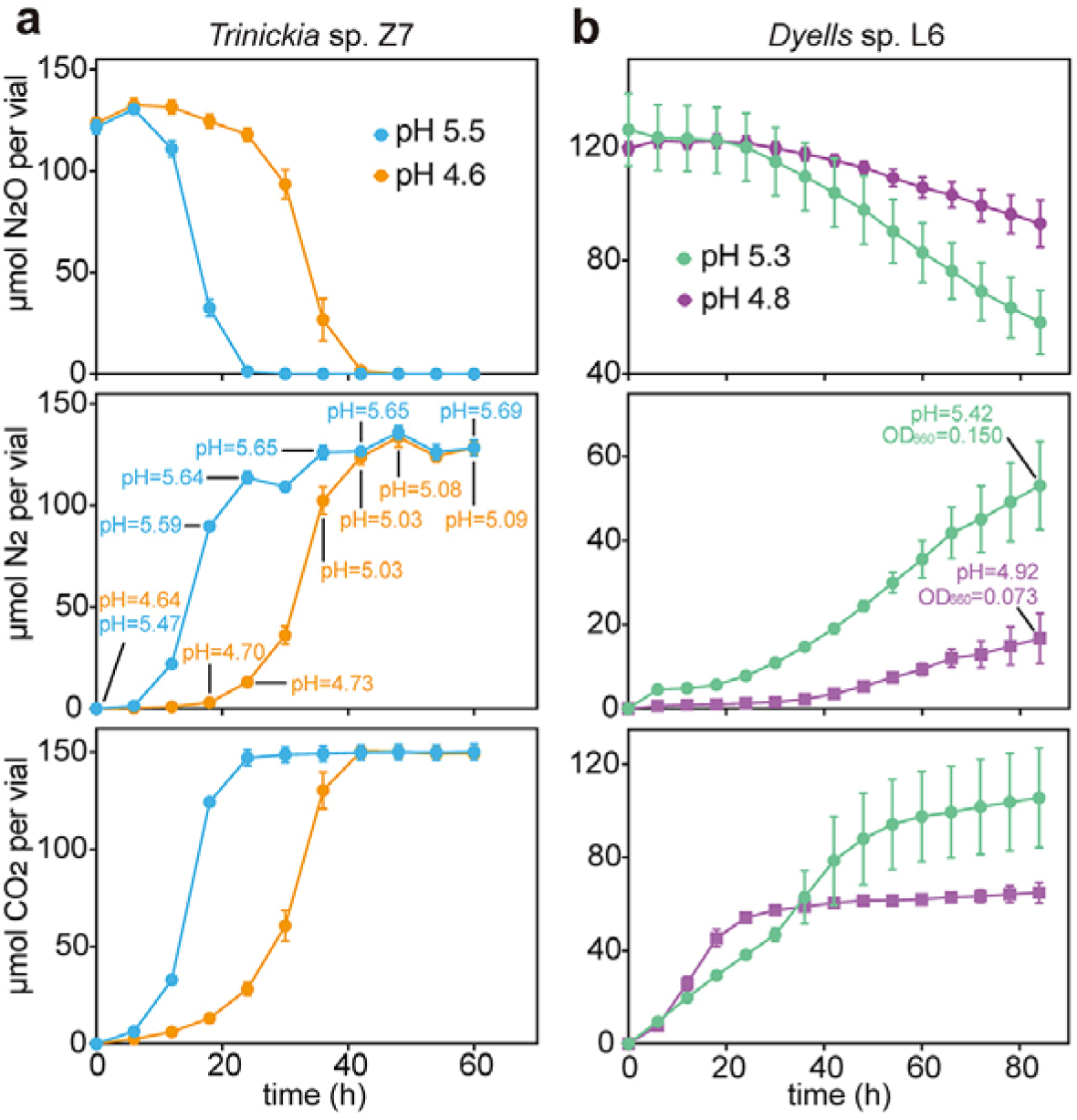
N_2_O reduction by *Trinickia* sp. Z7 and *Dyella* sp. L6 at acidic pH. The cells were pre-cultured in acidic medium under oxic conditions with vigorous stirring and with regular transfers to new medium to eliminate the possibility that the inoculum carried previously synthesized N_2_O reductase that could have matured under non-acidic conditions. From these pre-cultures, 300 μL (OD_660_=0.3) were inoculated into anoxic vials containing 30 mL of medium with 3 mL of NlJO in a He headspace. (**a**) N_2_O reduction by *Trinickia* sp. Z7 in acidic 1/10 TSB medium at pH 4.6 and pH 5.5. The medium was supplied with 20 mM H_3_PO_4_/20 mM Na_3_PO_4_ to mitigate pH increase induced by the metabolism of *Trinickia* when growing in TSB. pH values were measured at different timepoints using destructive sampling of replicates. (**b**) N_2_O reduction by *Dyella* sp. L6 in acidic R2A medium at pH 4.8 and 5.3. 20 mM MES buffer was used to maintain the medium pH.

**Extended Data Fig. 4.**
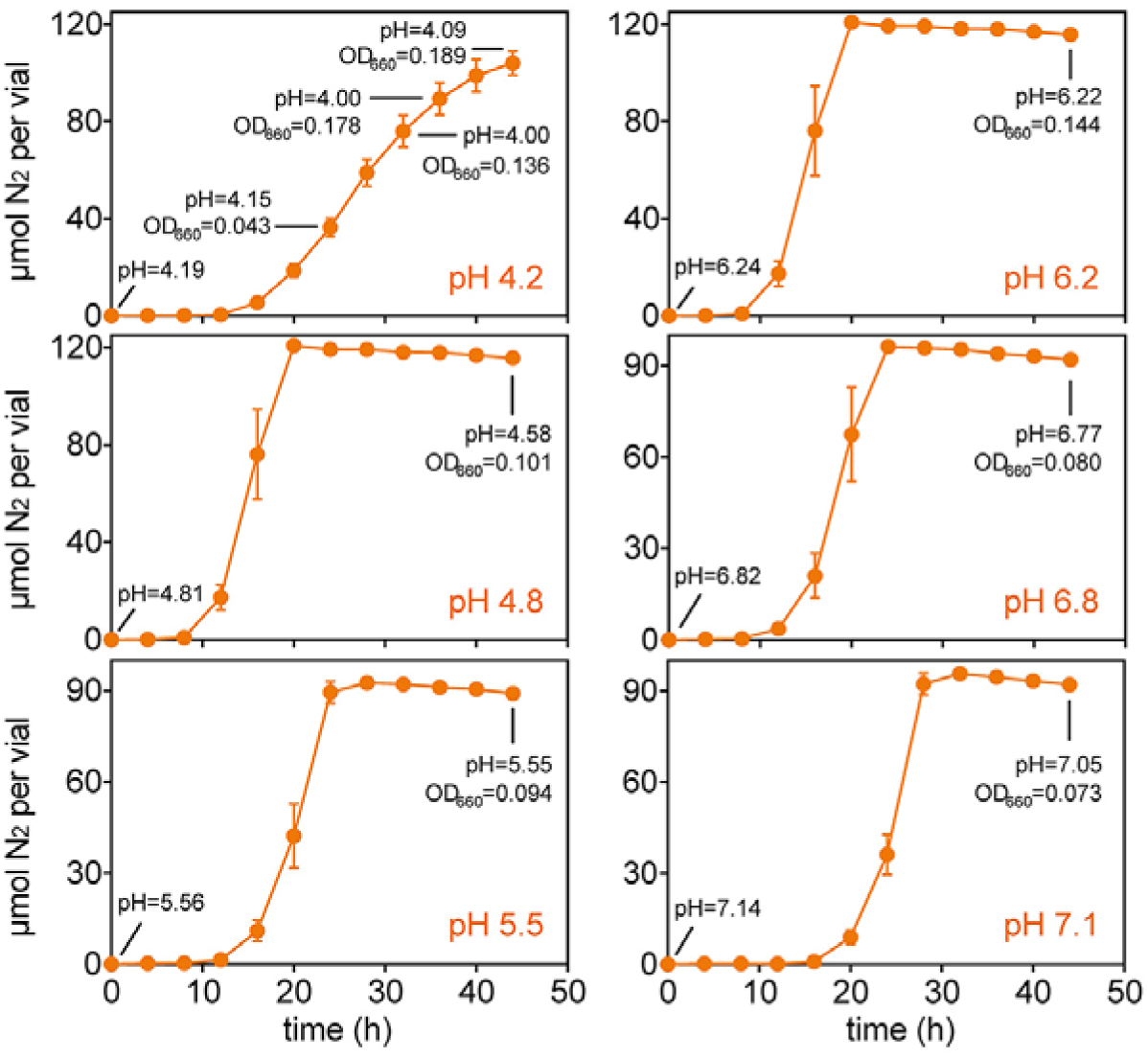
N_2_O reduction by *Trinickia* sp. Z7 in 20 mM MES buffered Sistrom’s media across a wide pH range. The cells were first grown in acidic (pH 4.2) Sistrom’s medium under oxic conditions with vigorous stirring and with regular transfers to new medium (>23 generations) to eliminate the possibility that the inoculum carried previously synthesized N_2_O reductase that could have matured under non-acidic conditions. Then, 300 μL was taken from these precultures and inoculated into 30 mL of Sistrom’s medium and incubated under continuous stirring (500 rpm). The medium had been set to different pHs before sterilization (indicated in each panel). Measured pH and OD at various time points are shown for each graph.. Each vial was supplemented with 3 mL of N_2_O in the headspace. Each treatment was measured in three replicate vials (n=3).

**Extended Data Fig. 5.**
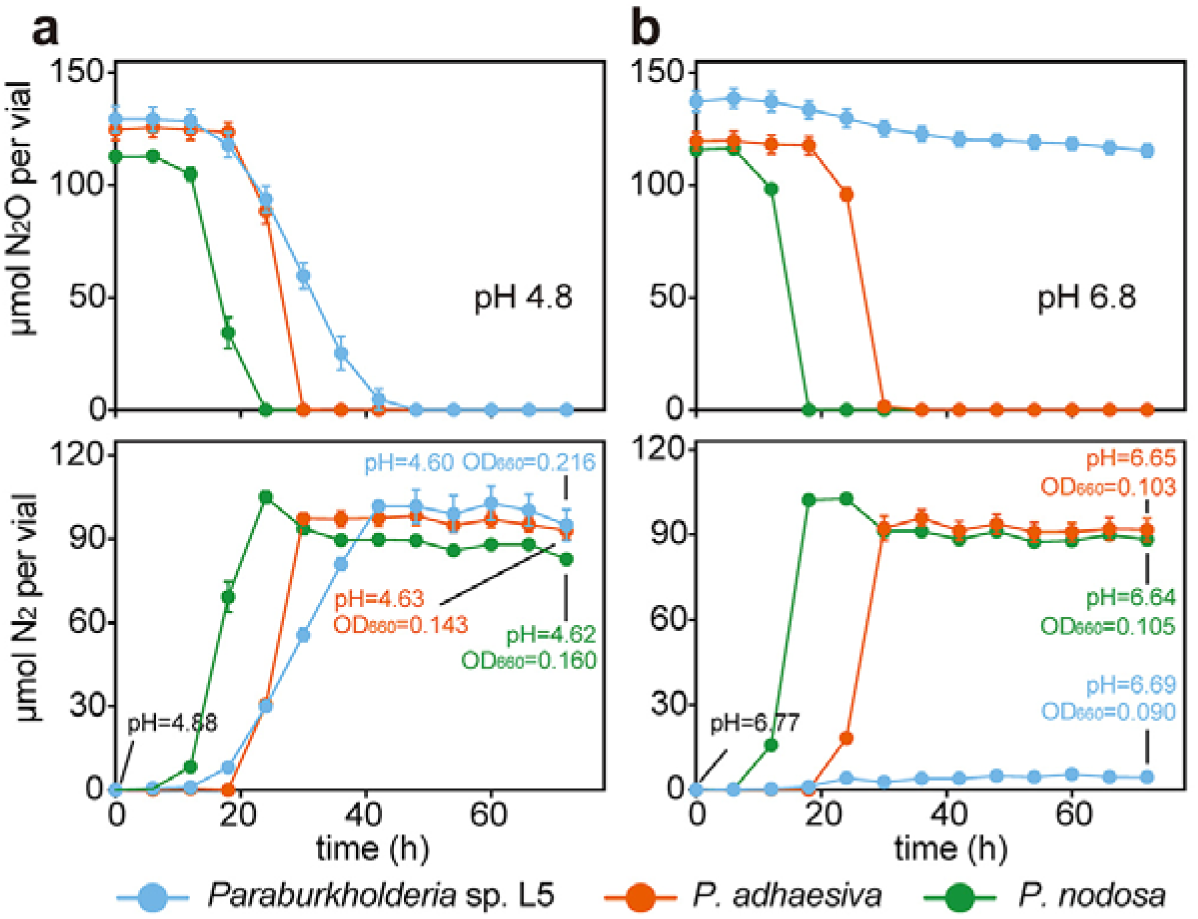
N_2_O reduction by three *Paraburkholderia* strains when incubated at acidic and circumneutral pH. Gas kinetics were monitored for the three *nosPQ* carrying strains *Paraburkholderia* sp. L5 (isolated in this study) and *P. adhaesiva and P. nodosa* (obtained from culture collections; Supplementary Table 2) when incubated anoxically in the presence of N_2_O at (**a**) pH 4.8 and (**b**) pH 6.8. The cells were first grown in acidic (pH 4.2) Sistrom’s medium under oxic conditions with vigorous stirring and with regular transfers to new medium to eliminate the possibility that the inoculum carried previously synthesized N_2_O reductase that could have matured under non-acidic conditions. Then, 300 μL was taken from these precultures and inoculated into 30 mL of Sistrom’s medium and incubated under continuous stirring (500 rpm). The medium had been set to different pHs before sterilization. Each vial was supplemented with 3 mL of NlJO and 1 mL of O_2_ in the headspace. Each treatment was measured in three replicate vials (n=3). *Paraburkholderia* sp. L5 grow poorly at pH=6.8 and reduced only a fraction of the provided N_2_O during the incubation.

**Extended Data Fig. 6.**
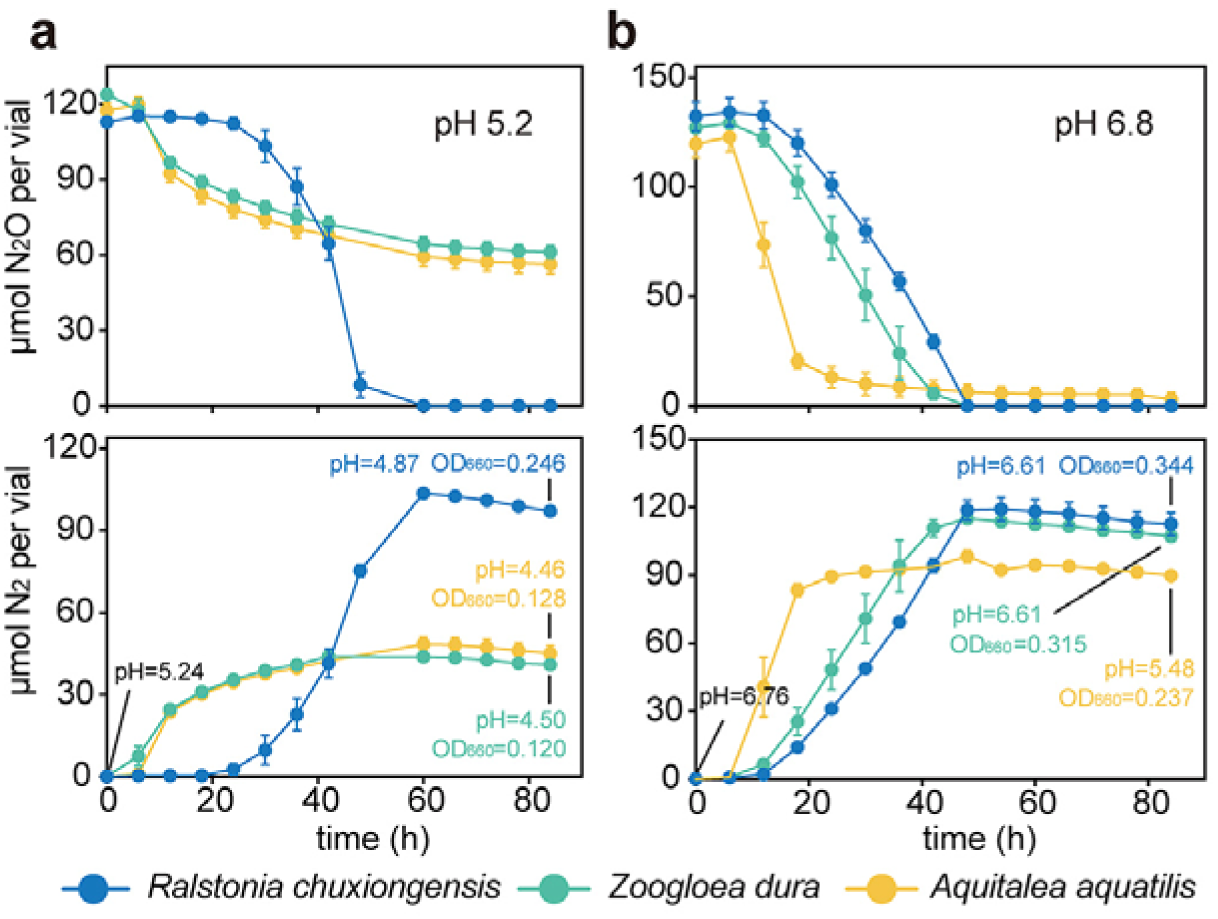
N_2_O reduction at acidic and circumneutral pH by three *nosPQ* carrying strains obtained from culture collections. The panels show N_2_O reduction and N_2_ production by the three *nosPQ* carrying strains *Ralstonia chuxiongensis, Zoogloea dura* and *Aquitaleea aquatilis* when incubated anoxically in the presence of N_2_O at (**a**) pH 5.2 and (**b**) pH 6.8. The experiment was designed to stringently test their ability to synthesize functional NosZ at low pH (as described for *Trinickia*). All the strains were first grown in acidic (pH 5.2) Sistrom’s medium under oxic conditions with vigorous stirring and with regular transfers to new medium to eliminate the possibility that the inoculum carried previously synthesized N_2_O reductase that could have matured under non-acidic conditions. Then, 300 μL was taken from these precultures and inoculated into 30 mL of Sistrom’s medium and incubated under continuous stirring (500 rpm). The medium had been set to different pHs before sterilization. Each vial was supplemented with 3 mL of NlJO and 1 mL of O_2_ in the headspace. Each treatment was measured in three replicate vials (n=3). The culture collections from which the strains were obtained are listed in Supplementary Table 2. The experiment was designed to stringently test their ability to synthesize functional NosZ at low pH (as adopted for *Trinickia*).

**Extended Data Fig. 7.**
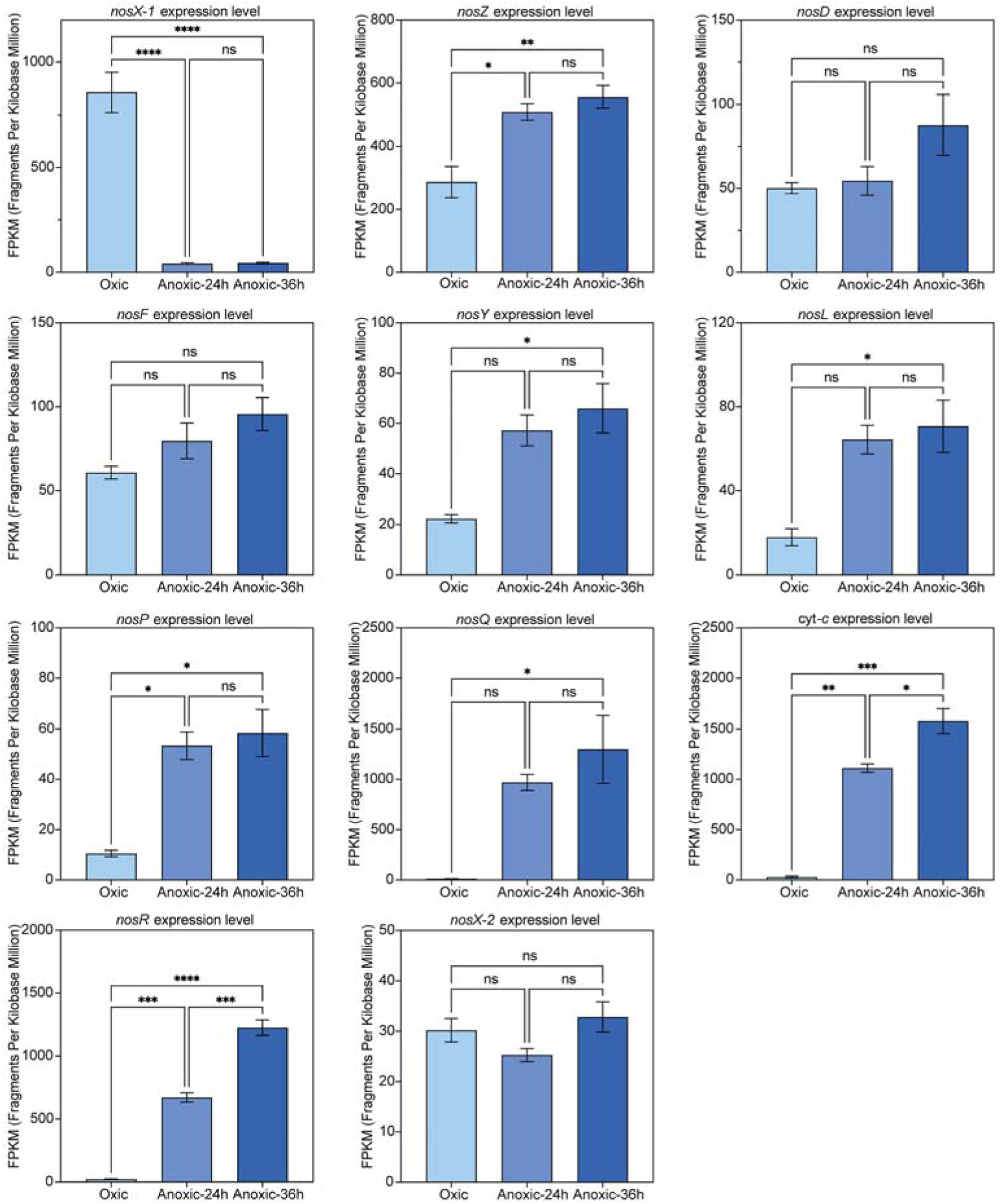
Transcription of genes within the *nos* cluster in *Trinickia* sp. Z7 during aerobic respiration and following N_2_O reduction at acidic pH. Samples were taken from *Trinickia* sp. Z7 in the pH 4.6 culture (Extended Data Fig. 3a) at three timepoints; the beginning of the incubation (Oxic), 24 h and 36 h of N_2_O reduction. *nosX*-1 is located in the *nos* cluster, while *nosX*-2 is somewhere else on the genome.

**Extended Data Fig. 8.**
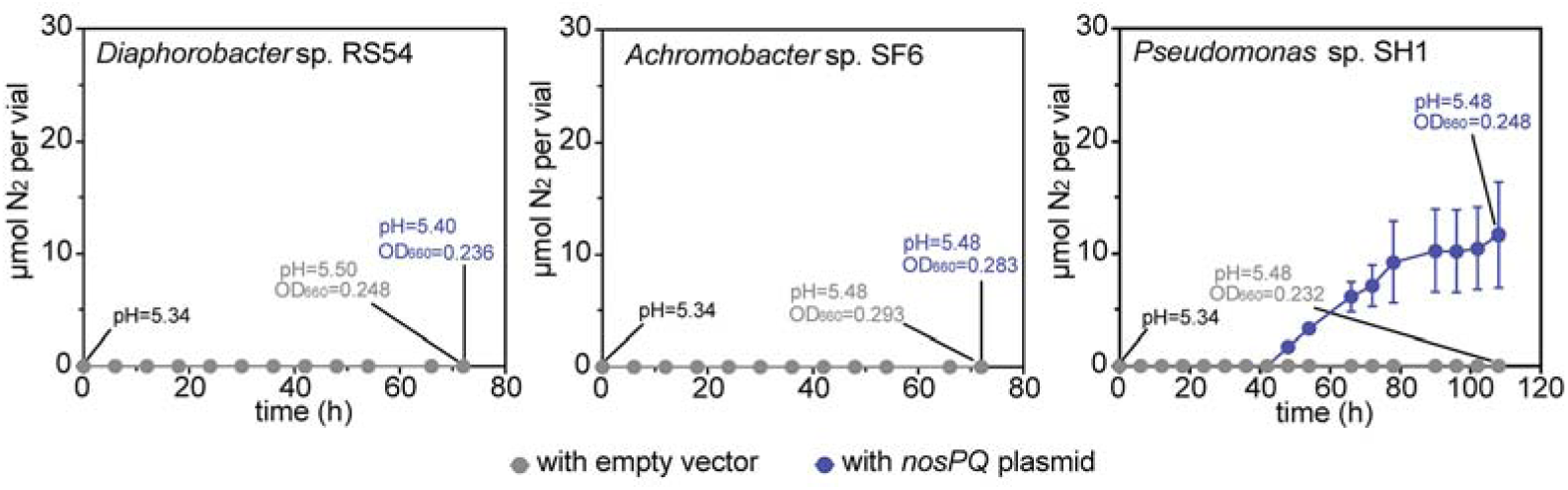
N_2_ production from N_2_O reduction by three strains transformed with a *nosPQ* carrying plasmid. The strain *Diaphorobacter* sp. RS54, *Achromobacter* sp. SF6, *Pseudomonas* sp. SH1 had been proven to reduce N_2_O in circumneutral pH medium, but not at low pH. They were transformed with a *nosPQ*-integrated plasmid and, for comparison, with a blank plasmid of the same type. 3 mL of N_2_O awas added into each vial, and N_2_O reduction was tested in 30 mL of 20 mM MES buffered 1/5 TSB medium (original pH=5.3). Three replicates were used (n=3); bars indicate SEM.

**Extended Data Fig. 9.**
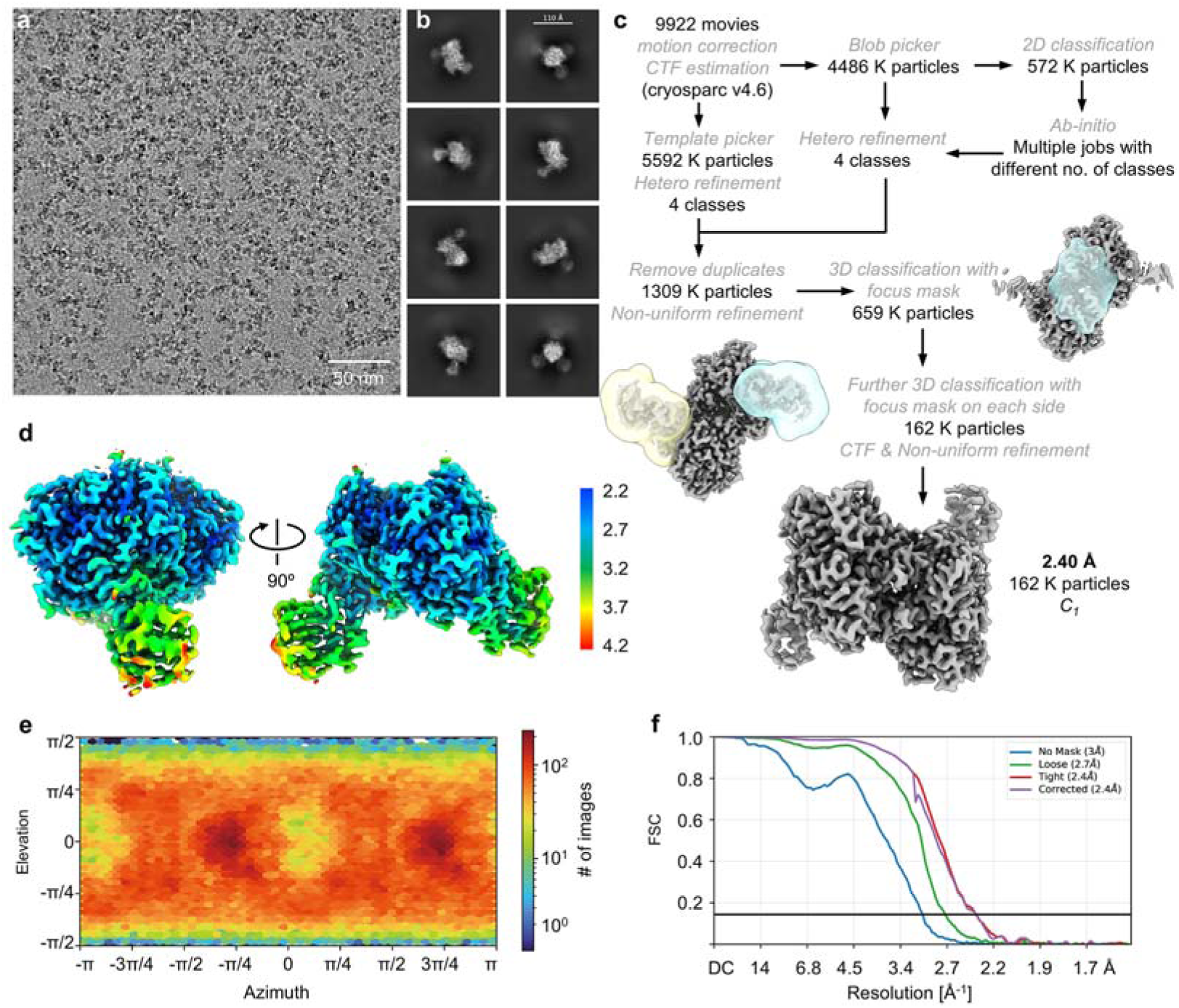
Data statistic and workflow for the cryo-EM data processing of the NosZPQ complex. (**a**) Representative micrograph of 9922 movies. (**b**) 2D class averages with different views. (**c**) Workflow for the data processing, leading to a refined map at 2.4 Å resolution using 162 K particles. (**d**) Local resolution maps for different orientations of the NosZPQ complex. (**e**) Direction distribution of particles used for 3D reconstruction. (**f**) Fourier shell correlation (FSC) curves.

**Extended Data Fig. 10.**
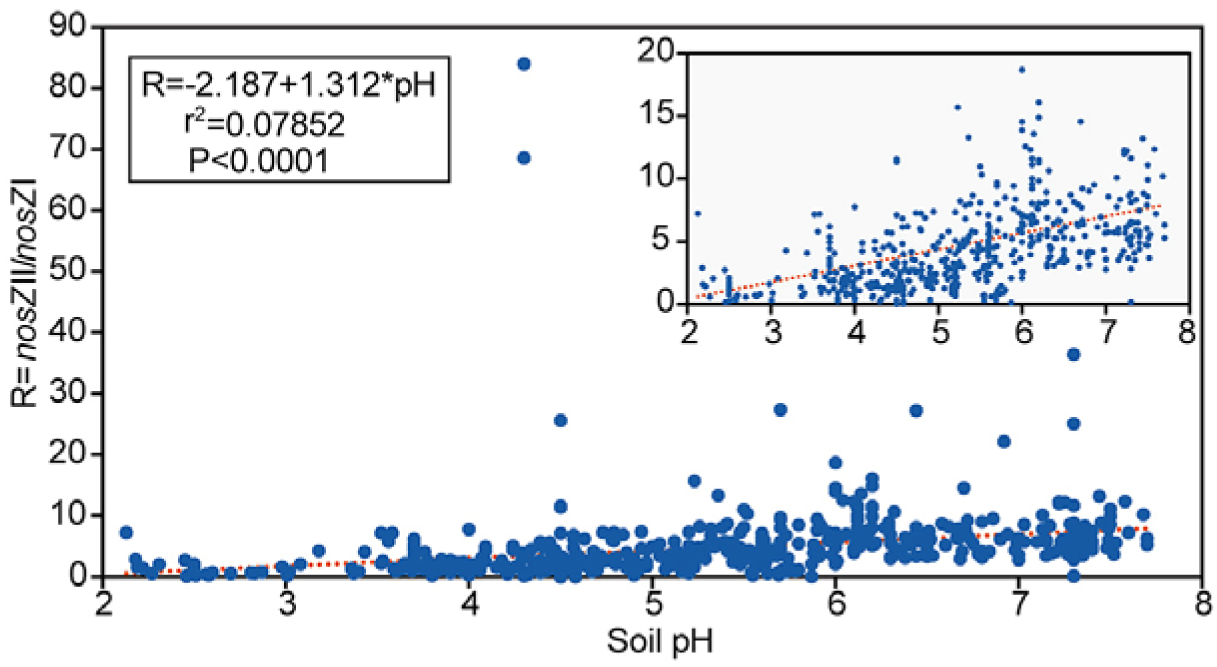
The relative abundance of genomes with *nosZ* clade II increase with soil pH. The panel shows the ratio between the abundance of *nosZ* clade II- and *nosZ* clade I-reads in the soil metagenomes (R), plotted against the pH of the soil from which the metagenomes had been extracted. The insert shows the same data scaled differently, to illustrate the trend for R<20 more clearly. The box shows the result of the linear regression analysis, and red line illustrated regression function.

## References

1 Cannon, A. J. Twelve months at 1.5 °C signals earlier than expected breach of Paris Agreement threshold. Nat Clim Change 15 (2025). 10.1038/s41558-025-02247-8

2 Cavicchioli, R. et al. Scientists’ warning to humanity: microorganisms and climate change. Nat Rev Microbiol 17, 569–586 (2019). 10.1038/s41579-019-0222-5

3 Stein, L. Y. & Lidstrom, M. E. Greenhouse gas mitigation requires caution. Science 384, 1068–1069 (2024). 10.1126/science.adi0503

4 Robertson, G. P. in Encyclopedia of Agriculture and Food Systems (ed Neal K. Van Alfen) 185–196 (Academic Press, 2014).

5 Ravishankara, A. R., Daniel, J. S. & Portmann, R. W. Nitrous Oxide (N_2_O): The Dominant Ozone-Depleting Substance Emitted in the 21st Century. Science 326, 123–125 (2009). 10.1126/science.1176985

6 Tian, H. Q. et al. A comprehensive quantification of global nitrous oxide sources and sinks. Nature 586, 248–256 (2020). 10.1038/s41586-020-2780-0

7 Bouwman, A. F., Boumans, L. J. M. & Batjes, N. H. Emissions of N_2_O and NO from fertilized fields: Summary of available measurement data. Global Biogeochem Cy 16, 6–1–6–13 (2002). 10.1029/2001gb001811

8 Shcherbak, I., Millar, N. & Robertson, G. P. Global metaanalysis of the nonlinear response of soil nitrous oxide (N_2_O) emissions to fertilizer nitrogen. Proc Natl Acad Sci U S A 111, 9199–9204 (2014). 10.1073/pnas.1322434111

9 Qiu, Y. P. et al. Intermediate soil acidification induces highest nitrous oxide emissions. Nat Commun 15, 2695 (2024). 10.1038/s41467-024-46931-3

10 Qian, H. Y. et al. Legacy effects cause systematic underestimation of N_2_O emission factors. Nat Commun 16, 2775 (2025). 10.1038/s41467-025-58090-0

11 Zumft, W. G. & Kroneck, P. M. Respiratory transformation of nitrous oxide (N_2_O) to dinitrogen by Bacteria and Archaea. Adv Microb Physiol 52, 107–227 (2007). 10.1016/S0065-2911(06)52003-X

12 Wijler, J. & Delwiche, C. C. Investigations on the Denitrifying Process in Soil. Plant Soil 5, 155–169 (1954). 10.1007/Bf01343848

13 Bergaust, L., Mao, Y. J., Bakken, L. R. & Frostegård, Å. Denitrification Response Patterns during the Transition to Anoxic Respiration and Posttranscriptional Effects of Suboptimal pH on Nitrogen Oxide Reductase in Paracoccus denitrificans. Appl Environ Microb 76, 6387–6396 (2010). 10.1128/Aem.00608-10

14 Bueno, E. et al. Anoxic growth of *Ensifer meliloti* 1021 by N_2_O-reduction, a potential mitigation strategy. Front Microbiol 6, 537 (2015). 10.3389/fmicb.2015.00537

15 Liu, B., Frostegård, Å. & Bakken, L. R. Impaired Reduction of N_2_O to N_2_ in Acid Soils Is Due to a Posttranscriptional Interference with the Expression of nosZ. mBio 5, 10.1128/mbio.01383–01314 (2014). 10.1128/mbio.01383-14

16 Liu, B. B., Morkved, P. T., Frostegård, Å. & Bakken, L. R. Denitrification gene pools, transcription and kinetics of NO, N2O and N2 production as affected by soil pH. FEMS Microbiol Ecol 72, 407–417 (2010). 10.1111/j.1574-6941.2010.00856.x

17 Frostegård, Å., Vick, S. H. W., Lim, N. Y. N., Bakken, L. R. & Shapleigh, J. P. Linking meta-omics to the kinetics of denitrification intermediates reveals pH-dependent causes of N_2_O emissions and nitrite accumulation in soil. The ISME Journal 16, 26–37 (2022). 10.1038/s41396-021-01045-2

18 Fujita, K. & Dooley, D. M. Insights into the Mechanism of N_2_O Reduction by Reductively Activated N_2_O Reductase from Kinetics and Spectroscopic Studies of pH Effects. Inorg Chem 46, 613–615 (2007). 10.1021/ic061843f

19 Müller, C. et al. Molecular interplay of an assembly machinery for nitrous oxide reductase. Nature 608, 626–631 (2022). 10.1038/s41586-022-05015-2

20 Zhang, L., Wüst, A., Prasser, B., Müller, C. & Einsle, O. Functional assembly of nitrous oxide reductase provides insights into copper site maturation. Proc Natl Acad Sci USA 116, 12822–12827 (2019). 10.1073/pnas.1903819116

21 Chapuis-Lardy, L., Wrage, N., Metay, A., Chotte, J. L. & Bernoux, M. Soils, a sink for N_2_O? A review. Global Change Biol 13, 1–17 (2007). 10.1111/j.1365-2486.2006.01280.x

22 Siljanen, H. M. P. et al. Atmospheric impact of nitrous oxide uptake by boreal forest soils can be comparable to that of methane uptake. Plant Soil 454, 121–138 (2020). 10.1007/s11104-020-04638-6

23 Palmer, K., Biasi, C. & Horn, M. A. Contrasting denitrifier communities relate to contrasting N_2_O emission patterns from acidic peat soils in arctic tundra. The ISME Journal 6, 1058–1077 (2012). 10.1038/ismej.2011.172

24 Buessecker, S. et al. Coupled abiotic-biotic cycling of nitrous oxide in tropical peatlands. Nat Ecol Evol 6, 1881–1890 (2022). 10.1038/s41559-022-01892-y

25 Zhong, J. M. et al. Strong N_2_O uptake capacity of paddy soil under different water conditions. Agr Water Manage 278, 108146 (2023). 10.1016/j.agwat.2023.108146

26 Zheng, X. Z. et al. Low pH inhibits soil nosZ without affecting N_2_O uptake. J Soil Sediment 23, 422–430 (2023). 10.1007/s11368-022-03324-7

27 Wang, X. H. et al. Using adaptive and aggressive N_2_O-reducing bacteria to augment digestate fertilizer for mitigating N_2_O emissions from agricultural soils. Sci Total Environ 903, 166284 (2023). 10.1016/j.scitotenv.2023.166284

28 He, G. et al. Sustained bacterial N_2_O reduction at acidic pH. Nat Commun 15, 4092 (2024). 10.1038/s41467-024-48236-x

29 Awala, S. I. et al. Nitrous oxide respiration in acidophilic methanotrophs. Nat Commun 15, 4226 (2024). 10.1038/s41467-024-48161-z

30 Lycus, P. et al. Phenotypic and genotypic richness of denitrifiers revealed by a novel isolation strategy. The ISME Journal 11, 2219–2232 (2017). 10.1038/ismej.2017.82

31 He, G. et al. A novel bacterial protein family that catalyses nitrous oxide reduction. Nature (2025). 10.1038/s41586-025-09401-4

32 Butterbach-Bahl, K., Baggs, E. M., Dannenmann, M. et al. Nitrous oxide emissions from soils: how well do we understand the processes and their controls? Philos T R Soc B 368, 20130122 (2013). 10.1098/rstb.2013.0122

33 Russenes, A. L., Korsaeth, A., Bakken, L. R. & Dörsch, P. Spatial variation in soil pH controls off-season N_2_O emission in an agricultural soil. Soil Biol Biochem 99, 36–46 (2016). 10.1016/j.soilbio.2016.04.019

34 Wang, Y. J. et al. Soil pH as the chief modifier for regional nitrous oxide emissions: New evidence and implications for global estimates and mitigation. Global Change Biol 24, E617–E626 (2018). 10.1111/gcb.13966

35 Hiis, E. G., Vick, S. H. W., Molstad, L. et al.. Unlocking bacterial potential to reduce farmland N_2_O emissions. Nature 630, 421–428 (2024). 10.1038/s41586-024-07464-3

36 Qu, Z., Wang, J. G., Almoy, T. & Bakken, L. R. Excessive use of nitrogen in Chinese agriculture results in high N_2_O/(N_2_O+N_2_) product ratio of denitrification, primarily due to acidification of the soils. Global Change Biol 20, 1685–1698 (2014). 10.1111/gcb.12461

37 Sanford, R. A., Wagner, D. D., Wu, Q. Z. et al. Unexpected nondenitrifier nitrous oxide reductase gene diversity and abundance in soils. Proc Natl Acad Sci USA 109, 19709–19714 (2012). 10.1073/pnas.1211238109

38 Abramson, J. et al. Accurate structure prediction of biomolecular interactions with AlphaFold 3. Nature 630, 493–500 (2024). 10.1038/s41586-024-07487-w

39 Pauleta, S. R., Dell’Acqua, S. & Moura, I. Nitrous oxide reductase. Coordin Chem Rev 257, 332–349 (2013). 10.1016/j.ccr.2012.05.026

40 Prasser, B., Schöner, L., Zhang, L. & Einsle, O. The Copper Chaperone NosL Forms a Heterometal Site for Cu Delivery to Nitrous Oxide Reductase. Angew Chem Int Edit 60, 18810–18814 (2021). 10.1002/anie.202106348

41 Wilks, J. C. & Slonczewski, J. L. pH of the cytoplasm and periplasm of *Escherichia coli*: rapid measurement by green fluorescent protein fluorimetry. J Bacteriol 189, 5601–5607 (2007). 10.1128/JB.00615-07

42 Berka, K. et al. MOLEonline 2.0: interactive web-based analysis of biomacromolecular channels. Nucleic Acids Res 40, W222–W227 (2012). 10.1093/nar/gks363

43 Sennett, L. B. et al. Determining how oxygen legacy affects trajectories of soil denitrifier community dynamics and N_2_O emissions. Nat Commun 15, 7298 (2024). 10.1038/s41467-024-51688-w

44 Nadeau, S. A. et al. Metagenomic analysis reveals distinct patterns of denitrification gene abundance across soil moisture, nitrate gradients. Environ Microbiol 21, 1255–1266 (2019). 10.1111/1462-2920.14587

45 Qin, H. L. et al. A few key nirK- and nosZ-denitrifier taxa play a dominant role in moisture-enhanced N_2_O emissions in acidic paddy soil. Geoderma 385, 114917 (2021). 10.1016/j.geoderma.2020.114917

## References

46 Staff, S. S. Soil taxonomy: A basic system of soil classification for making and interpreting soil surveys. 2nd edition edn, U.S. Department of Agriculture Handbook 436 (Natural Resources Conservation Service, 1999).

47 Molstad, L., Dorsch, P. & Bakken, L. R. Robotized incubation system for monitoring gases (O2, NO, N2O, N2) in denitrifying cultures. J Microbiol Meth 71, 202–211 (2007). 10.1016/j.mimet.2007.08.011

48 Li, W., Li, H., Liu, Y. D., Zheng, P. & Shapleigh, J. P. Salinity-Aided Selection of Progressive Onset Denitrifiers as a Means of Providing Nitrite for Anammox. Environ Sci Technol 52, 10665–10672 (2018). 10.1021/acs.est.8b02314

49 Wang, C. et al. Bacterial genome size and gene functional diversity negatively correlate with taxonomic diversity along a pH gradient. Nat Commun 14, 7437 (2023). 10.1038/s41467-023-43297-w

50 Chee-Sanford, J. C., Connor, L., Krichels, A., Yang, W. H. & Sanford, R. A. Hierarchical detection of diverse Clade II (atypical) *nosZ* genes using new primer sets for classical- and multiplex PCR array applications. J Microbiol Meth 172, 105908 (2020). 10.1016/j.mimet.2020.105908

51 Punjani, A., Rubinstein, J. L., Fleet, D. J. & Brubaker, M. A. cryoSPARC: algorithms for rapid unsupervised cryo-EM structure determination. Nat. Methods 14, 290–296 (2017). 10.1038/Nmeth.4169

52 Punjani, A., Zhang, H. W. & Fleet, D. J. Non-uniform refinement: adaptive regularization improves single-particle cryo-EM reconstruction. Nat. Methods 17, 1214–1221 (2020). 10.1038/s41592-020-00990-8

53 Jumper, J. et al. Highly accurate protein structure prediction with AlphaFold. Nature 596, 583–589 (2021). 10.1038/s41586-021-03819-2

54 Varadi, M. et al. AlphaFold Protein Structure Database: massively expanding the structural coverage of protein-sequence space with high-accuracy models. Nucleic Acids Res 50, D439–D444 (2022). 10.1093/nar/gkab1061

55 Goddard, T. D. et al. UCSF ChimeraX: Meeting modern challenges in visualization and analysis. Protein Sci 27, 14–25 (2018). 10.1002/pro.3235

56 Emsley, P. & Cowtan, K. Coot: model-building tools for molecular graphics. Acta Crystallogr D Biol Crystallogr 60, 2126–2132 (2004).

57 Liebschner, D. et al. Macromolecular structure determination using X-rays, neutrons and electrons: recent developments in Phenix. Acta Crystallogr D Struct Biol 75, 861–877 (2019).

58 Chen, V. B. et al. MolProbity: all-atom structure validation for macromolecular crystallography. Acta Crystallogr D Biol Crystallogr 66, 12–21 (2010). 10.1107/S0907444909042073

59 Chen, S. F. Ultrafast one-pass FASTQ data preprocessing, quality control, and deduplication using fastp. iMeta 2, e107 (2023). 10.1002/imt2.107

60 Buchfink, B., Reuter, K. & Drost, H. G. Sensitive protein alignments at tree-of-life scale using DIAMOND. Nat Methods 18, 366–368 (2021). 10.1038/s41592-021-01101-x

61 Menzel, P., Ng, K. L. & Krogh, A. Fast and sensitive taxonomic classification for metagenomics with Kaiju. Nat Commun 7, 11257 (2016). 10.1038/ncomms11257

