## Supplementary for "Two Small Proteins Enable N_2_O Reduction in Acidic Environments"

### List of Supplementary Tables

*Supplementary Tables are provided as a separate Excel file*

**Supplementary Table 1**. Composition of modified Sistrom's medium.

**Supplementary Table 2**. Isolates used for N_2_O reduction at low pH in this study.

**Supplementary Table 3**. Primers used for genome manipulation.

**Supplementary Table 4**. Cryo-EM data collection, refinement, and validation statistics.

**Supplementary Table 5**. NosP and NosQ protein sequences obtained from NCBI.

**Supplementary Table 6**. Global soil metagenomes and annotation results.

**Supplementary Table 7**. *nosZ* gene abundance at genus level in each sample and module classification.

**Supplementary Table 8**. Metagenome binning results.

**Supplementary Table 9**. The preparation of cells in buffered acidic (pH 4.2) Sistrom's medium.

**Supplementary Table 10**. Taxonomy and abundance of *nosPQ* genes in each soil metagenome.

**Supplementary Table 11**. Clade abundance and taxonomy.

### List of Supplementary Figures

**Supplementary Figure 1**. N_2_O reduction capacities of the acidic (pH 4.4) pine forest soil as observed in anaerobic incubations.

**Supplementary Figure 2**. Dissolved organic carbon content in soil samples collected at different timepoints (0 h, 6 h, 18 h, 42 h) across treatment groups.

**Supplementary Figure 3**. N_2_O reduction in paddy soil with different treatments.

**Supplementary Figure 4**. Shifts in microbial community composition at the DNA and mRNA levels after different amendments.

**Supplementary Figure 5**. Abundance and taxonomic classification of *nosZ* reads at genus level.

**Supplementary Figure 6**. Co-occurrence network analysis of *nosZ*-containing bacteria identified in metagenome results.

**Supplementary Figure 7**. N_2_ production and taxonomy of identified N_2_O reducers.

**Supplementary Figure 8**. Gas kinetics and pH changes of *Trinickia* sp. Z7 cultured in different acidic media.

**Supplementary Figure 9**. Transmission electron micrographs (TEM) of *Trinickia* sp. Z7 cells.

**Supplementary Figure 10**. N_2_O reduction of *Trinickia* sp. Z7 cultured in different MES buffered Sistrom's media with the presence of O_2_.

**Supplementary Figure 11**. N_2_O reduction of *Trinickia* Δ*nosPQ*::*Gm* mutant.

**Supplementary Figure 12**. Production of the NosZPQ complex.

**Supplementary Figure 13**. Taxonomy and counts of NosPQ sequences recruited from publicly available database at class (left panel) and genus level (right panel), respectively.

**Supplementary Figure 14**. Sample clustering by UPGMA based on the microbial composition of *nosPQ* genes.

**Supplementary Figure 15**. The *nosPQ* containing microbial composition and pH of each sample in three clusters.

**Supplementary Figure 16**. The abundance of *nosPQ* gene abundance as affected by soil pH.

**Supplementary Figure 17**. The ratio between the abundance of *nosPQ*- and *nosZ* clade I genes as affected by soil pH.

### 1. Supplementary methods

#### 1.1 Metagenome sequencing and binning

The procedures used for DNA extraction, metagenomic sequencing, and data processing have been comprehensively detailed in a previous study ^1^. On average, 201,981,223 reads were generated per sample. Kaiju (v1.9.2) ^2^ was used to get the taxonomic classification of clean reads against the NCBI-nr database (v2022-03-10) with five allowed mismatches and E-value of 0.00001. Gene abundances were quantified and normalized using RPM (Reads Per Million) values.

As for metagenomic binning, the clean reads were assembled into contigs using MEGAHIT (v1.2.9, k-min 27, k-max 127, k-step 10) ^3^ with the option of min-contig-len 500. Genome binning of the assembled contigs was conducted by MetaBAT2 (v2.15) ^4^, followed by refinement of the bins with RefineM (v1.4.3) ^5^. CheckM (v1.1.3) ^6^ was used to evaluate the quality of the bins and dRep (v3.0.0) ^7^ was used to remove the redundant bins. Taxonomic assignments of these non-redundant bins were performed using the GTDB-Tk (v1.7.0) ^8^. Genome annotation was carried out by RAST (v2.0) (https://rast.nmpdr.org/), and the predicted gene models were annotated using KofamKOALA (v2022-05-08) ^9^ against KEGG database (release 103.0) to identify the KEGG Orthologs (KOs).

#### 1.2 Metatranscriptome sequencing

Total RNA was extracted from soil samples using the RNA PowerSoil® Total RNA Isolation Kit (Mobio Laboratories Inc., Carlsbad, CA, USA). Isolated total RNA was qualified by 1.5% agarose gel electrophoresis, and quantified using an ultraviolet spectrophotometer. To enrich for mRNA, rRNA sequence-specific probes were hybridized with the total RNA, and the resulting rRNA-probe complexes were subsequently removed using magnetic beads. At last, the total mRNA was purified with ethanol.

For the construction of metatranscriptome libraries, the detailed methodology has been previously described ^10^. Sequencing was conducted using the Illumina NovaSeq platform (Illumina, San Diego, CA, USA) along with the PE150 strategy, generating an average of 12,904,127,040 base pairs per sample. The sequencing workflow and subsequent bioinformatic processing were consistent with those employed for metagenome sequencing. Gene expression levels were normalized and represented using RPM values.

#### 1.3 Genome sequencing

The isolates *Trinickia* sp. Z7, *Paraburkholderia* sp. L5, *Dyella* sp. L6 were cultured at 30°C with agitation in natural pH 1/10 TSB medium, pH 4.8 1/10 TSB medium, natural pH R2A medium, respectively. Genomic DNA was extracted from the cell pellets with an E.Z.N.A.® Bacterial DNA Kit (Omega Bio-tek) according to the manufacturer’s instructions. The sequencing libraries were prepared following the guide of Hieff NGS® MaxUp II DNA Library Prep Kit for Illumina®. Genomes were sequenced using the Illumina HiSeq platform with the PE150 strategy.

Raw paired-end reads were trimmed and quality controlled by Trimmomatic (v0.36) ^11^. Genome assembly was performed with SPAdes (v3.5.0) ^12^, and the quality of the genome was assessed using CheckM (v1.0.18) ^6^. Gene models were identified by using GeneMarkS (v4.28) ^13^, and all predicted gene models were annotated by KofamKOALA (v2023-04-01) ^9^ against KEGG release 106.0 to identify the KOs. Reference genomes were downloaded from NCBI database and annotated using the same pipeline. The phylogenetic maximum likelihood tree was constructed by IQ-TREE (v2.2.0.3) ^14^ and finally visualized on iTOL ^15^. Average Nucleotide Identity (ANI) and Average Amino acid Identity (AAI) values were calculated by FastANI (v1.3) ^16^ and EzAAI (v1.2.3) ^17^, respectively.

#### 1.4 Transcriptome sequencing

Total RNA was isolated and purified using the RNA 6000 nano kit (Agilent). The quality and integrity of the RNA were determined using a NanoDrop spectrophotometer (Thermo Scientific) and a Bioanalyzer 2100 system (Agilent). Zymo-Seq RiboFree Total RNA Library Kit was used to remove rRNA from total RNA. Following this, cDNA fragments of the desired size (approximately 380 bp) were selected, and library fragments were purified using the AMPure XP system (Beckman Coulter, Beverly, CA, USA). Sequencing was performed on the Illumina Novaseq 6000 platform, generating 16,703,996~20,320,894 reads per sample.

The raw sequencing data were filtered using fastp (v0.22.0) ^18^, which included the removal of reads containing adapter, ploy-N, and low quality reads. The quality-filtered reads were then aligned to the *Trinickia* sp. Z7 genome using Bowtie2 (v2.5.1) ^19^, resulting in a total of 16,590,473–20,081,692 mapped reads per sample (mapping proportion 98.82–99.51%). Gene-level read counts were determined with HTSeq (v0.9.1) ^20^, and gene expression abundances were normalized and represented as FPKM (Fragments Per Kilo bases per Million fragments mapped) values.

#### 1.5 N_2_O reduction at low pH of *nosPQ* genes containing bacteria

Firstly, three acid-tolerant *Paraburkholderia* strains (Supplementary Table 2) were tested due to their close phylogenetic relationship with *Trinickia*. *Paraburkholderia* sp. L5, was isolated from acidic paddy soil (pH 4.8) in this study. The *Paraburkholderia adhaesiva* strain was acquired from the Guangdong Microbial Culture Collection Center (GDMCC) and was originally isolated from forest acidic lateritic red earth soils with a pH of 4.0~5.0 in South China ^21^. The *Paraburkholderia nodosa* strain was obtained from the China Center of Industrial Culture Collection (CICC), which was isolated from nodules on *Mimosa scabrella* in Brazil ^22^. Prior to N_2_O reduction experimentation, cells were first activated in R2A medium (pH 6.76) and then transferred three times in Sistrom's medium (pH 4.2, 20 mM MES buffer) following the same protocol as described for *Trinickia* sp. Z7 preculture. For the experimental setup, 300 μL of cells were inoculated into 30 mL of media (pH 4.8 and 6.8, buffered with 20 mM MES) with 3 mL N_2_O and 1 mL O_2_ added in the headspace.

Subsequently, we selected three non-acidophilic isolates which harbor the novel genes of *nosP* and *nosQ*, and tested their N_2_O reduction capabilities under both acidic and circumneutral pH conditions. The *Ralstonia chuxiongensis* and *Zoogleoa dura* strains were obtained from the GDMCC. *Ralstonia chuxiongensis* was originally isolated from tobacco-planting soil in South China ^23^, while *Zoogleoa dura* was derived from forest soil in Republic of Korea ^24^. The *Aquitalea aquatilis* strain was sourced from the China Center for Type Culture Collection (CCTCC), which was isolated from a waterfall in Republic of Korea ^25^. Cells were first precultured in acidic Sistrom's medium (pH 5.2, buffered with 20 mM MES) and then verified their N_2_O reduction abilities at pH 5.2 and 6.8, following the experimental protocol described previously.

#### 1.6 Gene deletion

Three fragments were PCR amplified to construct the conjugative plasmid: a region of 802 bp upstream of *nosPQ* was amplified using primers NOS-5F and NOS-5R (Supplementary Table 3), a gentamycin resistance cassette of 890 bp was amplified from plasmid pJQ200SK (Sangon Biotech Co., Ltd., Shanghai, China) using primers NOS-GmF and NOS-GmR, and a region of 786 bp downstream region of *nosPQ* was amplified using primers NOS-3F and NOS-3R. After each amplification, the three fragments were joined using fusion PCR by primers NOS-5F and NOS-3R. The PCR product with right size was further purified and integrated into the pCVD442 plasmid (Sangon Biotech Co., Ltd., Shanghai, China). The resulted plasmid pCVD442-Δ*nosPQ*::*Gm* was then electroporated into *Escherichia coli* DH5α λpir. Cells were transformed and then plated on LB plates containing 50 μg/mL ampicillin and 30 μg/mL gentamycin. A resistant colony was verified using PCR and sequencing with primers 442F3 and 442R3. After that, this colony was cultured in LB broth with 50 μg/mL ampicillin and 30 μg/mL gentamycin for plasmid isolation. The extracted plasmid was introduced into a diaminopimelic acid (DAP) auxotroph of *Escherichia coli* β2155 by electroporation. Cells were transferred and plated on LB plates containing 100 μg/mL ampicillin and 0.5 mM DAP, and resistant colonies were cultured and used as donor strains.

As for the conjugation, cultures of donor (*Escherichia coli* β2155) and recipient (*Trinickia* sp. Z7) cells were grown overnight to reaching an OD_660_ value of >0.8. Bacterial conjugation with 0.15 mL of each of donor and recipient cells was carried out on a 0.22 μm filter, which was incubated for one day on the surface of a 1/5 TSA plate containing 0.5 mM DAP. Then, this filter was cultured in 30 mL of 1/5 TSB medium with 50 μg/mL gentamycin for one day. Transconjugants were selected on 1/5 TSA plates with gentamycin. Single colonies were then cultured and plated onto the same 1/5 TSA plates but supplied with 10% sucrose to select double-crossover mutants. Resistant colonies were validated by both PCR and DNA sequencing with primers NOS-outF and NOS-outR, which covered the whole sequences from upstream to downstream. Primers NOS-inF and NOS-inR were used to confirm the absence of *nosPQ* genes. Colonies with right sequences and without *nosPQ* genes, which qualified as target *Trinickia* Δ*nosPQ*::*Gm* mutant, were selected for subsequent experiments.

#### 1.7 Complementation assay

A fragment containing *nosPQ* genes was PCR amplified from *Trinickia* sp. Z7 genome and integrated into the pBBR1MCS-2 plasmid by homologous recombination. Two different promotors were used, including the plasmid's built-in promotor pLacI and the promotor of the *Trinickia* sp. Z7 *nos* gene cluster pnosZ. HIS tag was also added to the C-terminal of NosQ to verify their successful expressions, resulting a total of 4 different reconstructed plasmids. The sequences of these expression vectors were verified using PCR and sequencing with primers BBR-SEQ-F and BBR-SEQ-R. The details about these primers and plasmids are provided in Supplementary Table 3. The plasmids were verified by sequencing and then electroporated into *Trinickia* Δ*nosPQ*::*Gm* mutant cells, while the original pBBR1MCS-2 plasmid was used as blank control. Cells were transformed and then plated on 1/5 TSA plates containing 50 μg/mL kanamycin, and resistant colonies were verified the successful electroporation by PCR. After obtaining the right transformed colonies, 300 μL of bacteria were added to 30 mL of acidic (pH 4.8, with 20 mM MES buffer) Sistrom's medium containing 25 μg/mL gentamycin and 25 μg/mL kanamycin to test N_2_O reduction.

### 2. Supplementary results

#### 2.1 Carbon source and liming promoted N_2_O reduction

Various amendments were applied to characterize the N_2_O reducing bacteria in acidic paddy soil, including application of organic carbon and pH management. During the three-day incubation period, 33, 45, and 179 μmol N_2_ were produced in sterile water-amended soil (S), glucose-amended soil (G)，and limed soil (C), respectively (Extended Data Fig. 1c). After a 12-hour anaerobic incubation, the paddy soil began to reduce N_2_O in S group soils, and the N_2_O reduction rate remained stable (~0.7 μmol N_2_O/hour) without much increase throughout the entire incubation. The pH value in this group remained nearly constant, fluctuating only slightly from 4.8 to 4.9. In the glucose-amended soil, the N_2_O reduction rate was initially enhanced but declined sharply from 2.8 to 0.2 μmol N_2_O/hour after 24 hours of incubation, probably due the depletion of glucose with CO_2_ production rate decreasing. Additionally, glucose amendment resulted in a significant increase in soil pH, rising to 5.6 by the end of the incubation, a phenomenon consistent with previous findings ^26^. On the contrary, the pH value of the limed soil decreased from an initial pH value of 7.8 to 5.8 over the course of the incubation. Although N_2_O reduction persisted continuously in the limed soil, the reduction rate gradually declined from 4.5 to 2.3 μmol N_2_O/hour after 30 hours, likely due to the gradual reacidification of the soil.

The results further indicated that the dissolved organic carbon (DOC) content remained relatively stable throughout the incubation period in the sterile water-amended soils (Supplementary Fig. 2). In contrast, liming led to a slight increase in DOC content, rising from 0.071 to 0.169 mg/g soil after 18 hours of incubation, a trend consistent with observations in other limed acidic soils ^27,28^. In the glucose-amended soil, the DOC content reached to 0.528 mg/g soil after 6 hours but gradually declined as the incubation progressed.

To further investigate whether elevated available carbon could enhance N_2_O reduction, a tenfold increase in glucose addition was employed. The results demonstrated that N_2_O reduction commenced after 12 hours of anoxic incubation (Supplementary Fig. 3). The addition of 200 mg of glucose facilitated a more substantial N_2_O reduction, resulting in the production of approximately 100 μmol N_2_ by the end of the incubation period. In the soil slurry, N_2_O was completely reduced, exhibiting a pattern similar to that observed in the limed soil. Interestingly, the pH of this soil mixture increased to 6.5 at the end.

#### 2.2 Shifts of whole microbial community

Compared to the initial soil (O), the incubation with N_2_O did not significantly alter the Shannon indices at either the DNA or mRNA level in the control group soils (S) (Supplementary Fig. 4a). The principal-coordinate analysis also shows that the O group samples clustered closely with the S group samples (Supplementary Fig. 4b). At the three sampling time points, the differences in microbial transcripts between different S group samples were marginal. However, the addition of glucose (G) and liming (C) substantially enhanced alpha diversity and induced significant shifts in the microbial community structure (P<0.001), as evidenced by both meta-omics analyses. Furthermore, the transcriptional composition of the samples exhibited significant variations across the different time points as shown on the PCoA plot.

In the metagenomic analysis, Proteobacteria, Actinobacteria, Acidobacteria and Chloroflexi were the most abundant taxa. Only a small increase of Proteobacteria in S42 soil was detected compared to original soil. Both glucose addition and liming greatly increased the relative abundance of Firmicutes (Supplementary Fig. 4c), accounting for 4.1% and 2.8% of the total reads in the G42 and C42 samples, respectively. At genus level, these enriched Firmicutes reads were mainly affiliated with *Clostridium* and *Bacillus*, while a specific Proteobacteria genus, *Noviherbaspirillum*, was notably enriched in the C42 samples (Supplementary Fig. 4d).

In the metatranscriptome sequencing results, transcripts from Firmicute accounted for the majority of total reads, especially after the addition of glucose and liming. These Firmicutes bacteria exhibited exceptionally high transcriptional activity, which reached to 80.2% in the G18 samples. In the acidic soils (S group), the transcriptional profiles at the 6th hour were comparable to those of the original soil, but the transcripts of Firmicutes dominated after 18 hours of anaerobic incubation. *Clostridium* and *Bacillus* were the most abundant genera in the transcripts of all samples. Other Bacilli taxa, such as *Neobacillus*, *Pseudoneobacillus*, which were recently classified as distinct genera from *Bacillus* species ^29,30^, also displayed high abundance of transcripts. Furthermore, liming significantly enhanced the transcriptional activity of *Noviherbaspirillum*, which accounted for 3.9% of the total transcripts after 42 hours of incubation.

#### 2.3 *nosZ* gene and transcripts analysis

To further reveal the predominant and active N_2_O reducing bacteria in the soil microbial community, the *nosZ* reads were extracted from both the metagenome and metatranscriptome and subsequently analyzed. The Shannon indices showed that after a 42-hour incubation the *nosZ* gene diversities increased in all soils compared to original soil (Extended Data Fig. 2a). Additionally, the *nosZ*-containing microbial community underwent significant shifts, with clear separation among the different treatment groups (Extended Data Fig. 2b). In general, the *nosZ* gene reads were predominantly derived from Proteobacteria and Acidobacteria, Bacteroidetes, Verrucomicrobia, Chloroflexi and Planctomycetes across all samples (Extended Data Fig. 2c). Both glucose addition and liming substantially increased the abundance of Firmicutes *nosZ* genes, which were primarily affiliated with *Bacillus* and *Neobacillus* (Supplementary Fig. 5a). In the acidic soil S42 and limed soil C42, *Massilia*, *Rhodanobacter*, *Dyella*, and *Paraburkholderia* were found to be the main sources of increased *nosZ* gene reads (Supplementary Fig. 5b). Furthermore, liming enhanced the abundance of *nosZ* genes originating from *Novi**herbaspirillum*, *Herbaspirillum*, *Pseudogulbenkiania*.

While metagenomic analysis revealed a diverse array of *nosZ* genes across all soil samples, metatranscriptomic results showed that neither different sampling times nor amendments significantly increased the diversity of *nosZ* transcripts during the N_2_O reduction process (Extended Data Fig. 2a). The S6 soils exhibited transcriptional profiles closely resembling those of the original soil (O) on the PCoA plot, with both displaying similar *nosZ* transcript compositions (Extended Data Fig. 2b & 2c). In the other samples, the majority of *nosZ* transcripts originated from Firmicutes, which contributed to a relatively homogeneous *nosZ* transcriptional profile. *Bacillus* and *Neobacillus* dominated the *nosZ* transcripts (Supplementary Fig. 5a). Among Proteobacteria, *Noviherbaspirillum* produced the most abundant *nosZ* transcripts, especially after 42 hours of incubation with liming (Supplementary Fig. 5b).

To investigate the abundance shifts of *nosZ*-containing bacteria in differently cultured soils，the co-occurrence network analysis was used. Firstly, the abundance of *nosZ* genes was summarized at the genus level. A total of 321 genera met the cutoff criterion of showing occurrence in >20% samples. After correlation calculation in R and module classification by Gephi, 8 modules were divided (Supplementary Table 7, Supplementary Fig. 6a), among which module 2, module 6 and module 7 had the highest abundance of *nosZ* genes (Supplementary Fig. 6b). Specifically, module 6 was mainly composed of *Bacillus*, *Neobacillus* and other Firmicutes, which were enriched with glucose and liming amendments. Module 7 was predominantly consisted of *Noviherbaspirillum*, only having great adaption to the elevated pH conditions induced by liming. Module 2 included a group of bacteria with increased *nosZ* gene abundances in both C42 soil and S42 soil (Supplementary Fig. 5). While liming promoted their growth, these bacteria were also found to be enriched in the sterile water control soils, which kept acidic throughout the whole incubation. It implies these increased *nosZ* gene-containing bacteria in module 2 might be acid-tolerant N_2_O reducers, such as *Rhodanobacter*, *Dyella*, *Paraburkholderia*, *Polaromonas*, *Desulfosporosinus*, *Rhodoferax*, *Ralstonia*, *Trinickia*. Among these genera, *Rhodanobacter* ^31,32^, *Polaromonas* ^32^, *Desulfosporosinus* ^33^ isolates have already been reported to reduce N_2_O at acidic pH. Additionally, *Methylocella* and *Methylacidiphilum nosZ* genes were also detected in these soils from this study, but they were divided into module 8 and module 5, respectively.

Through metagenome binning, commonly only dominant taxa with relatively small genome sizes could be assembled to meet high-quality thresholds ^34^. In this study, an average of >30 billion base pairs were sequenced for each sample, but only a total of 31 bins with completeness >75% and contamination <10% were obtained (Supplementary Table 8). Among these, six bins were found to contain *nosZ* genes according to KEGG Orthology database annotation, including two Acidobacteriota, two Proteobacteria, one Firmicutes and one Gemmatimonadota. The assembled *Trinickia* and *Noviherbaspirillum* genomes were identified as the potential N_2_O reducers enriched in the S42 soil and C42 soil, respectively. These two Proteobacteria genomes contained a copy of *nosZ* clade I and a full set of denitrification genes. The Gemmatimonadota bacterium has been considered as an important N_2_O sink in agricultural soils and reduced N_2_O at pH 5 ^35^. The results also suggested that some Acidobacteriota were *nosZ* containing bacteria, which might also be acid-resistant N_2_O reducers.

#### 2.4 Isolation and verification of N_2_O reducers

In the isolation efforts Ⅰ, Bacilli were identified as the target bacteria for isolating acid-tolerant N_2_O reducers according to the analysis of meta-omics results. By using heated soil dilutions, a total of 64, 187, 24, and 364 isolates were obtained on the circumneutral 1/10 TSA medium, circumneutral LB medium, acidic LB medium and acidic soil extract–glucose agar medium, respectively. 90 out of all 639 isolates were randomly selected and sequenced their 16S rRNA genes. Results showed that all sequenced isolates were identified as Bacilli genera. Through *nosZ* clade II PCR, 146 out of all 639 isolates were amplified the target band. The right PCR products derived from 43 randomly selected isolates were sequenced, and NCBI BLAST results revealed that 40 of these 43 sequences were annotated as *nosZ* genes affiliated with *Baciilus*, *Neobacillus*. Through the N_2_O reduction experiment in pure bacterial culture, these *nosZ* gene containing *Baciilus*, *Neobacillus* isolates were found to be incapable of reducing N₂O under acidic conditions, despite their ability to reduce N₂O at circumneutral pH. These findings highlight a potential disconnect between *nosZ* gene expression and enzymatic functionality under acidic conditions.

In the isolation efforts Ⅱ, 14, 43, 47 isolates were randomly picked from the cultured plates of soil extract–glucose agar medium, beef extract-peptone medium and 1/10 TSA medium, respectively. The N_2_O reduction experiment results of pure cultures showed that only 9 out of all 104 isolates could reduce N_2_O at either circumneutral or acidic pH (Supplementary Fig. 7a). These N_2_O reducers were identified as 7 *Trinickia* isolates, 1 *Paraburkholderia* isolate and 1 *Dyella* isolate (Supplementary Fig. 7b), all of which were isolated from the acidic soil extract-glucose medium.

When cultured in circumneutral 1/10 TSB medium (pH 6.3), *Trinickia* and *Dyella* isolates could reduce N_2_O, whereas the *Paraburkholderia* sp. L5 strain grew poorly and was unable to reduce N_2_O. In acidic 1/10 TSB medium with an initial pH of 4.8, *Trinickia* and *Paraburkholderia* isolates successfully reduced N_2_O, accompanied by a gradual neutralization of the medium pH. One of the isolates, named *Trinickia* sp. Z7, had the best N_2_O reduction performance in acidic 1/10 TSB medium (Supplementary Fig. 7a), although it also increased the medium pH to circumneutral (from 4.9 to 6.5) (Supplementary Fig. 8a). In contrast, when cultured in acidic SEG medium, *Trinickia* sp. Z7 effectively reduced N_2_O without increasing the medium pH. Instead, the pH value decreased from 4.5 to 4.3 (Supplementary Fig. 8b), a phenomenon consistent with the observations (Extended Data Fig. 1b) when 1 g of paddy soil was inoculated into the same medium.

#### 2.5 Isolates reducing N_2_O in acidic media

To verify microbial N₂O reduction under acidic conditions, the inoculum must be free of pre-formed N₂O reductases from non-acidic environments. Accurate measurement of whole-cell activity also requires cells to remain dispersed, since aggregation can create local pH microenvironments that mask the true phenotype. To avoid such unexpected pH increase, the *Trinickia* sp. Z7 inoculum was cultured under aerobic conditions and transferred for three rounds in strictly controlled acidic medium, in which 20 mM H_3_PO_4_/20 mM Na_3_PO_4_ was used to retard the pH increase. After each round of incubation for one day in 100 mL of 1/10 TSB medium, the pH value increased from 4.6 to 5.0, reaching an OD_660_ value of ~0.22. Through such three transfer cultures, the cells were deemed suitable for subsequent experiments. As a result, the pre-cultured cells exhibited N₂O reduction activity across a wide pH range, from 4.6 to 7.2, with optimal performance observed between pH 5.0 and 6.0. During continuous pH monitoring of N₂O reduction (Extended Data Fig. 3a), the medium maintained sustained acidity, with only a minor increase from pH 4.6 to 5.0, demonstrating acid-tolerant N_2_O reduction of isolate Z7. In the case of the *Dyella* sp. L6 strain, it failed to grow in acidic 1/10 TSB medium (pH 4.8) (Supplementary Fig. 7a), but exhibited growth in acidic R2A medium (pH 5.3). The inoculum increased the medium pH from 5.3 to approximately 5.5 after each one-day incubation. After three successive pre-culture transfers, the cells obtained in this way also showed the ability to reduce N_2_O at acidic pH (Extended Data Fig. 3b), only causing a marginal increase in the medium pH.

In Sistrom's medium, cell preparation of *Trinickia* sp. Z7 followed the same protocol as in the 1/10 TSB medium experiments. However, the medium pH decreased after each one-day incubation, maintaining a consistently acidic environment (Supplementary Table 9). Transmission electron micrographs (TEM) revealed that *Trinickia* cells produced intracellular storage compounds and formed visible particles during aerobic growth (Supplementary Fig. 9). These globular structures were consistently visible both inside and outside cells, and could be stained with Sudan black B; they are likely probably poly-β-hydroxybutyrate (PHB), as previously observed in *Trinickia dabaoshanensis ^36^* and *Trinickia soli* ^37^.

Inoculation of these acidic cells into non-buffered acidic Sistrom's medium induced progressive acidification during bacterial growth (Fig. 1c). Successful N_2_O reduction still occurred under such strongly acidic conditions after 8 hours of inoculation. We further evaluated the N₂O reduction of *Trinickia* sp. Z7 in MES-buffered Sistrom's medium across a wider pH range. The results showed that N₂O reduction was observed across all tested pH values ranging from 4.2 to 7.1 (Extended Data Fig. 4), showing a wide pH tolerance of N_2_O reductase.

To investigate the effects of O_2_ on N_2_O reduction, experiments were conducted with varying concentrations of O₂. The results indicated that the medium became acidified with cell growth. Cellular growth coupled with oxygen consumption leads to a further decrease in the pH of the medium. Despite the pH decrease, the cultures successfully reduced N₂O even in the presence of O₂ in the headspace at both acidic and circumneutral pH (Supplementary Fig. 10).

#### 2.6 Genome analysis of acid-tolerant N_2_O reducers

The genome size of *Trinickia* sp. Z7 is 5,752,624 bp with a 63% GC ratio, while the *Paraburkholderia* sp. L5 genome length is 7,175,350 bp with a 65% GC ratio. Quality assessment results show that both genomes have a 99.95% completeness. The ANI and AAI values of these two genomes are 79.62% and 75.72% respectively, confirming that they represent distinct species ^16^.

By KEGG Orthology database annotation, it is found that *Trinickia* sp. Z7 contains 2,197 unique KOs with the full-fledged denitrification genes *(narG, nirK, norB, and nosZ)*, which is in consistent with metagenome binning results. *Paraburkholderia* sp. L5 has 2,443 unique KOs, harboring 1,993 KOs in common with Z7. It is an incomplete denitrifier lacking nitrite reductase (*nir)* genes, but nitrogen fixation genes (*nifD*, *nifH*, and *nifK*) are present in the genome. Besides, both genomes also harbor assimilatory nitrate reduction genes *(nasA, nasD, and nasE)* and some genes for pH tolerance, such as decarboxylation and deamination of amino acids involved in the proton-consuming reactions (*gdhA*, *adiA*), basic compounds producing (*ureA, ureB, ureC*), cation and anion efflux pumps (*kdpA, kdpB, kdpC, kdpD, kdpE, kdpF*) ^38^. Interestingly, both genomes contain two copies of *phbC*, three copies of *phbB*, which enable these bacteria to produce polyhydroxyalkanoates (PHAs) when supplied with sugars, fatty acids, or amino acids ^39^.

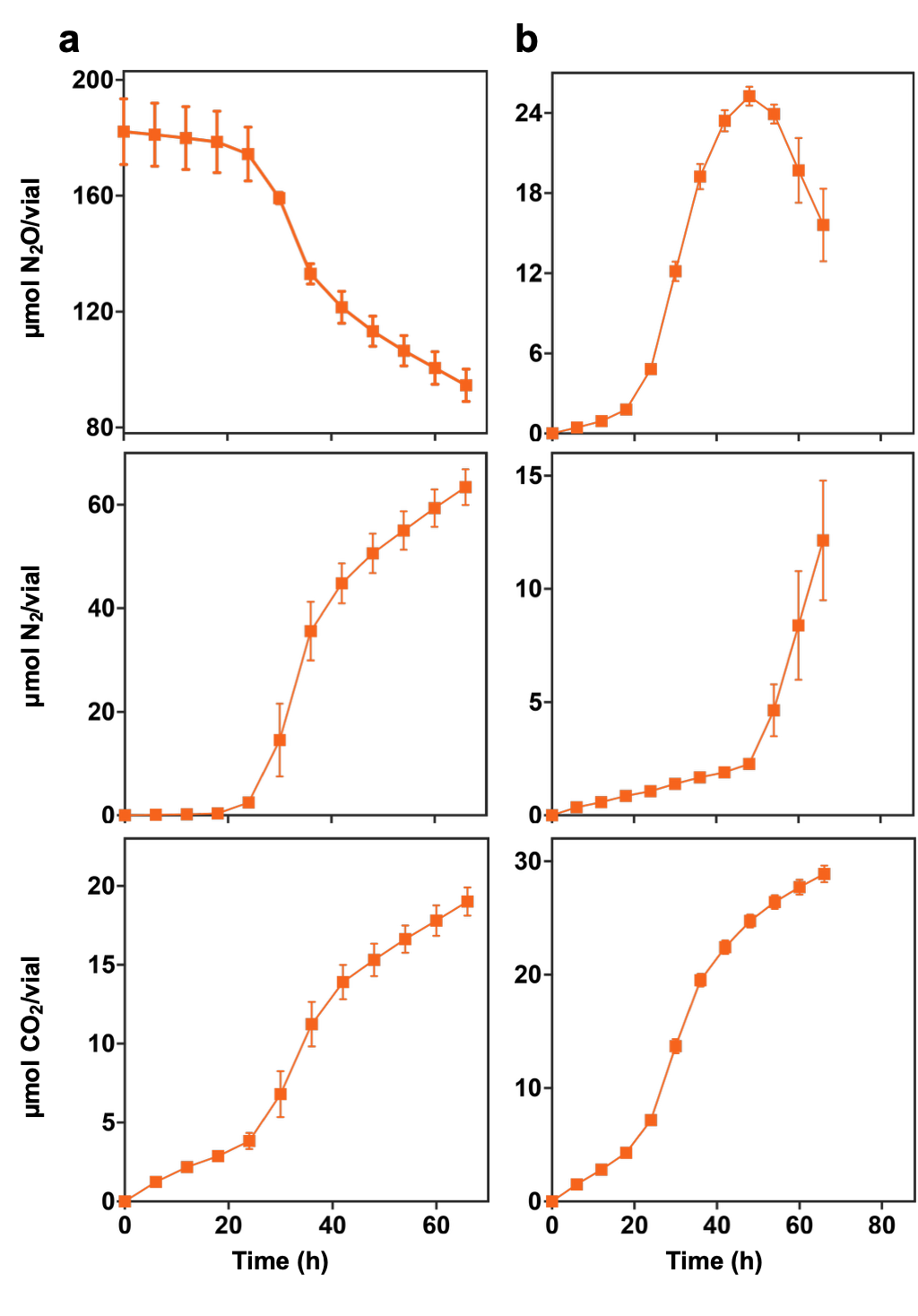

### Supplementary Figures 1-17

**Supplementary Fig. 1 |** **N_2_O reduction capacities of the acidic (pH 4.4) pine forest soil as observed in anaerobic incubations.** Gas kinetics was monitored in 30 g of intact soil. (**a**) Addition with 180 μmol N_2_O. (**b**) Addition with 48 μmol NO_3_^-^.

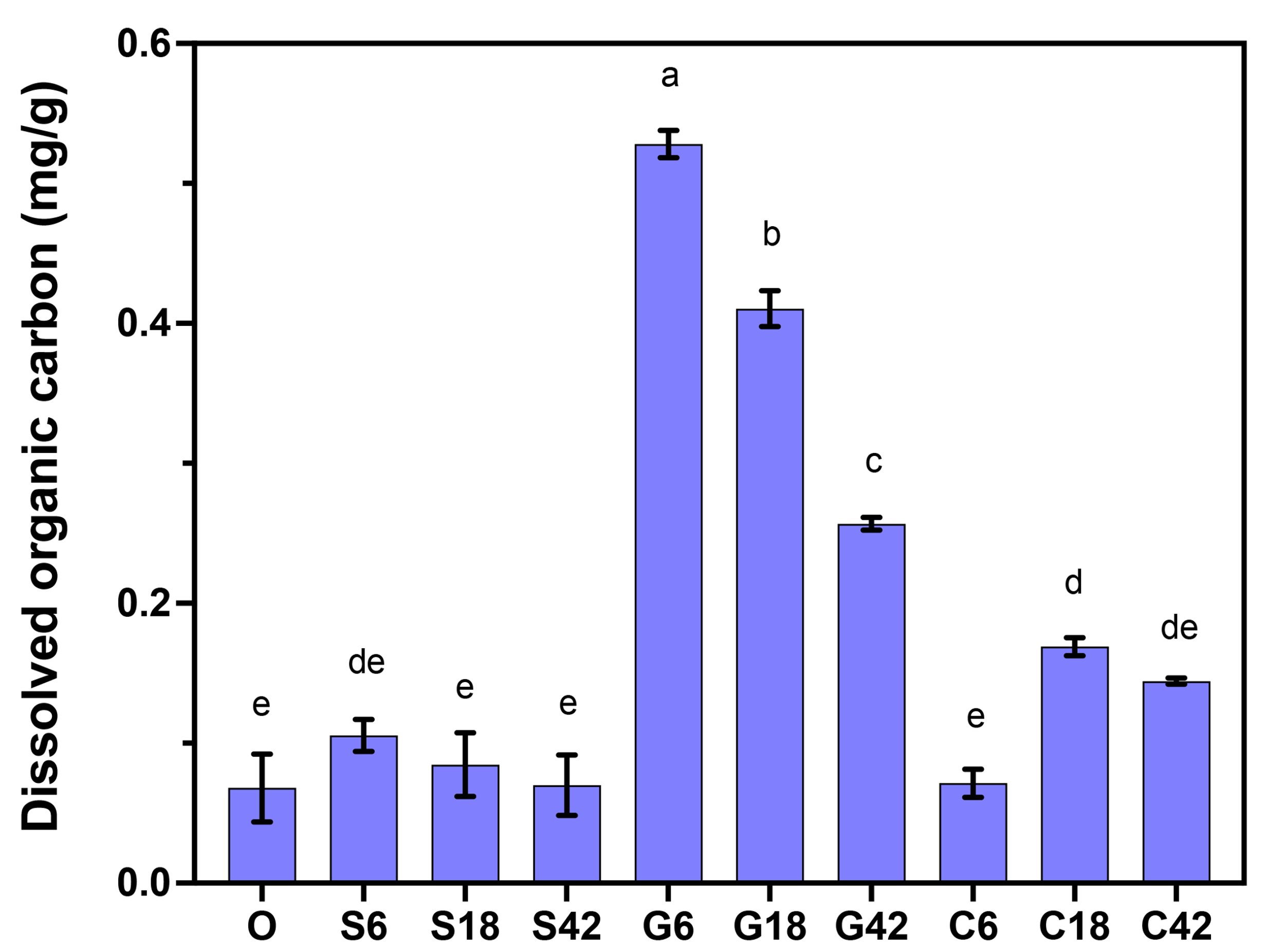
**Supplementary Fig. 2 | Dissolved organic carbon content in soil samples collected at different timepoints (0 h, 6 h, 18 h, 42 h) across treatment groups.** The data are expressed as mean values above the columns with ordinary one-way analysis of variance (ANOVA) followed by a Fisher’s Least Significant Difference (LSD) test. O: Oxic soil sampled only at time 0, serving as the original soil control. S: 2 mL of sterile distilled water added as control treatment. G: Addition of 2 mL of a 10 mg/mL glucose solution. C: Addition of 2 mL of a 135 mM Ca(OH)_2_ solution

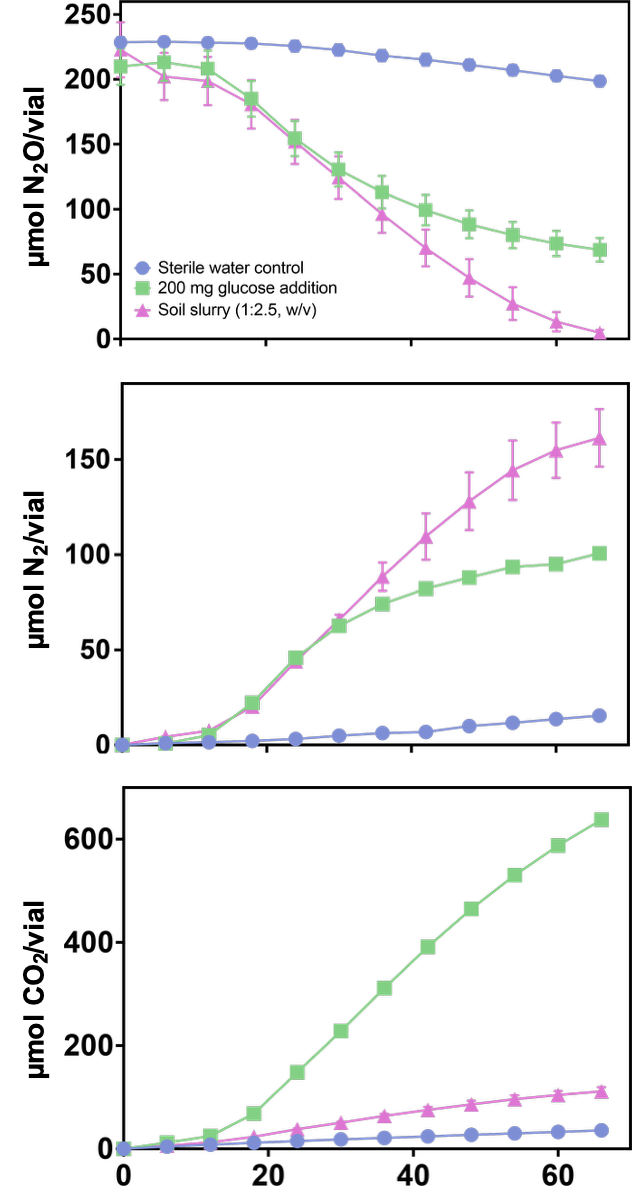

**Supplementary Fig. 3 | N_2_O reduction in paddy soil with different treatments.** 10 g of processed paddy soil was activated in each vial. 2 mL of 100 mg/mL glucose solution was used to test the increased carbon source effect. 25 mL of sterile water was added into the vials to make the soil slurry. A control treatment was prepared by adding 2 mL of sterile distilled water to the soil.

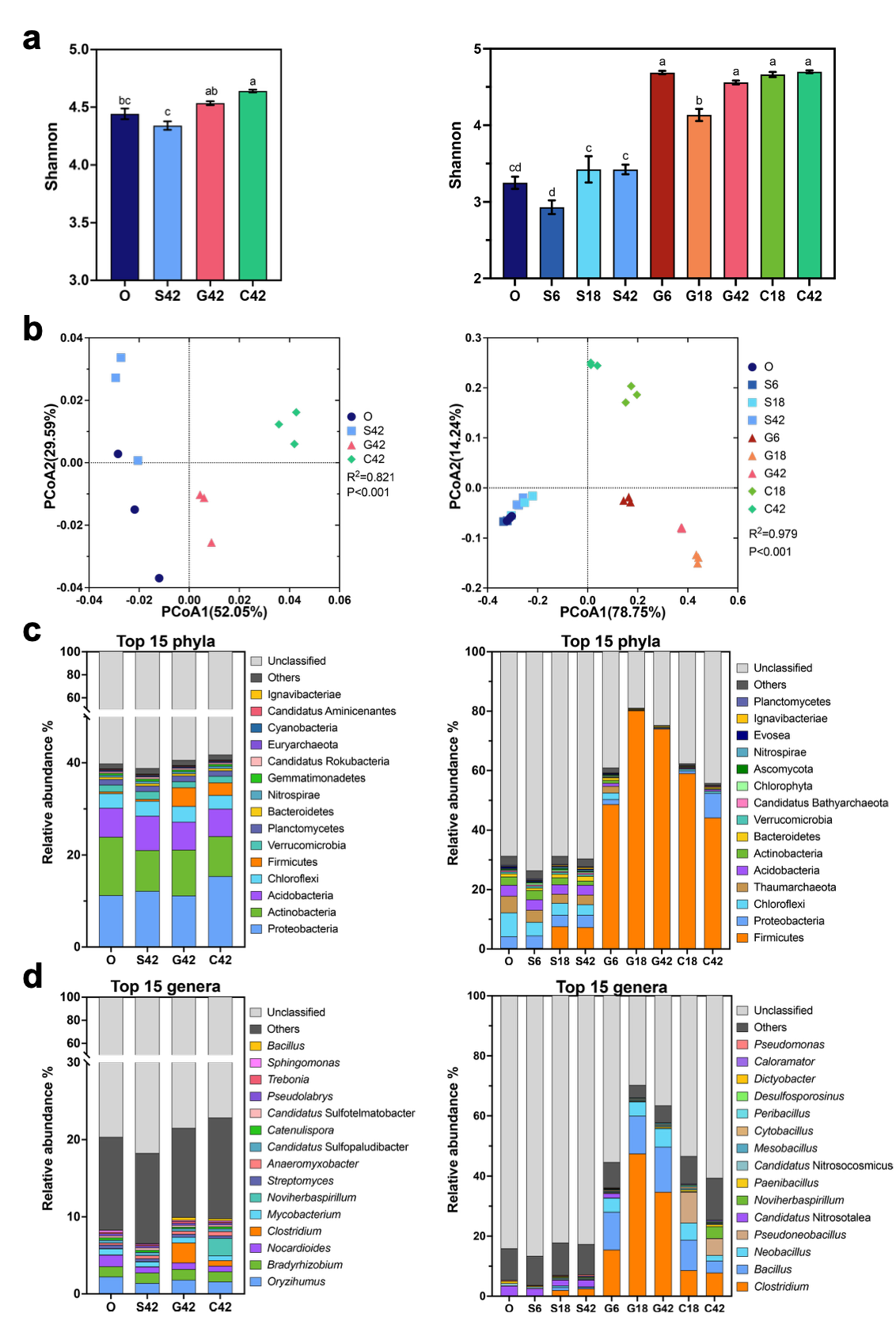

**Supplementary Fig. 4 |** **Shifts in microbial community composition at the DNA and mRNA levels after different amendments.** Soil samples (10 g, ww) were exposed to one of three pretreatments and then incubated in sealed, anoxic vials containing He plus ~204 μmol N_2_O in headspace. O: Oxic soil sampled only at time 0, serving as the original soil control. S: 2 mL of sterile distilled water added as control treatment. G: Addition of 2 mL of a 10 mg/mL glucose solution. C: Addition of 2 mL of a 135 mM Ca(OH)_2_ solution. Soil samples were collected from treatments S, G and C at 42 h after N_2_O addition for metagenomics analysis and at 6, 18, 42 hours after N_2_O addition for metatransctiptomics analysis. The four-group figures (left) show metagenomics results and the nine-group figures (right) show metatranscriptomics results. (**a**) Alpha diversities measured by Shannon index with ordinary one-way analysis of variance (ANOVA) followed by a Fisher's Least Significant Difference (LSD) test. (**b**) Principal-coordinate analysis (PCoA) along with corresponding PERMANOVA comparisons (9,999 permutations) for each sample. The ordinations were calculated based on the Bray‒Curtis distances. (**c**), (**d**) The relative abundances of top 15 phyla and genera obtained from each sequencing results.

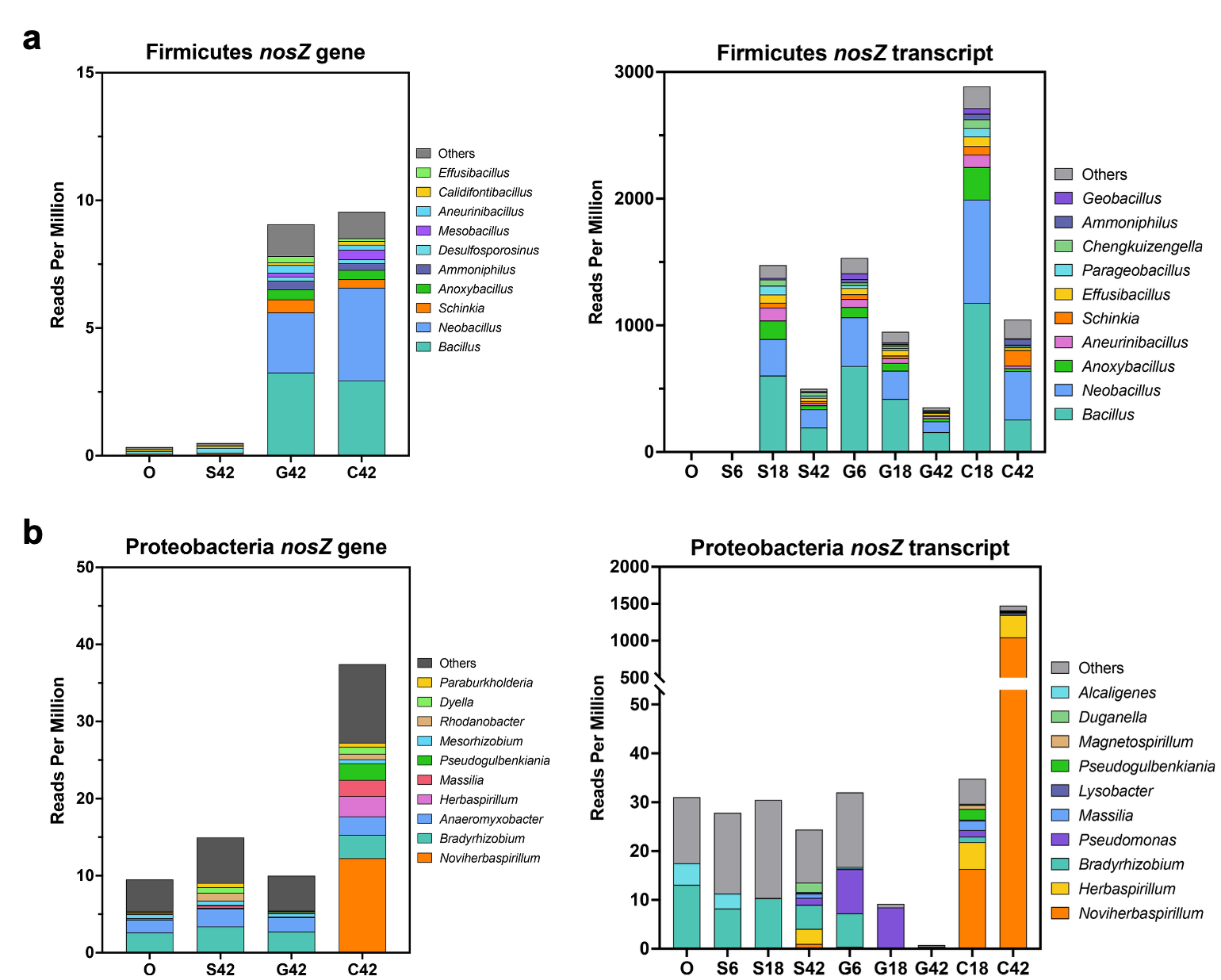

**Supplementary Fig. 5 | Abundance and taxonomic classification of *nosZ* reads at genus level.** (**a**) Firmicutes *nosZ* genes (left panel) and transcripts (right panel). (**b**) Proteobacteria *nosZ* genes (left panel) and transcripts (right panel). The details of treatments are same as described in Supplementary Figure 4.

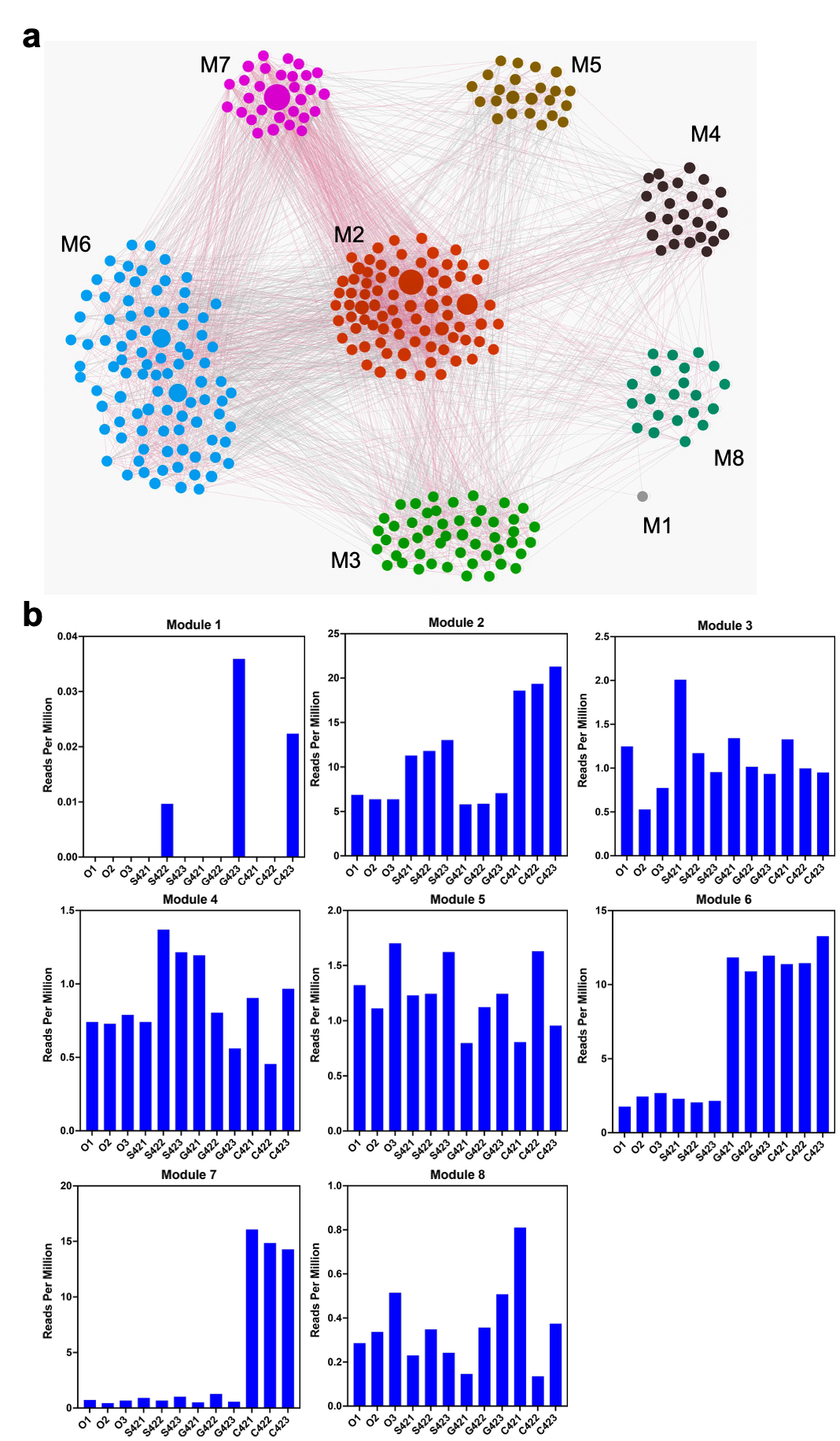

**Supplementary Fig. 6 | Co-occurrence network analysis of *nosZ*-containing bacteria identified in metagenome results.** (**a**) Co-occurrence network of each module. The abundance of *nosZ* reads in each sample were calculated and summarized into genus level. A total of 321 genera that occurred in >20% of all 12 samples were preserved. Co-occurrence was detected using pairwise Spearman’s correlations with cutoffs of Spearman’s rho >0.6 and FDR P<0.05. This co-occurrence network contained 321 nodes and 3,427 edges with a modularity value of 0.742. The module class was divided in Gephi and different modules were presented by different colors. Each node represents a single genus and the size reflected its average abundance in all samples. The red edges between nodes mean positive correlations while gray edges were negative correlations. (**b**) Abundances of different modules in all samples. The details of each module are shown in Supplementary Table 7.

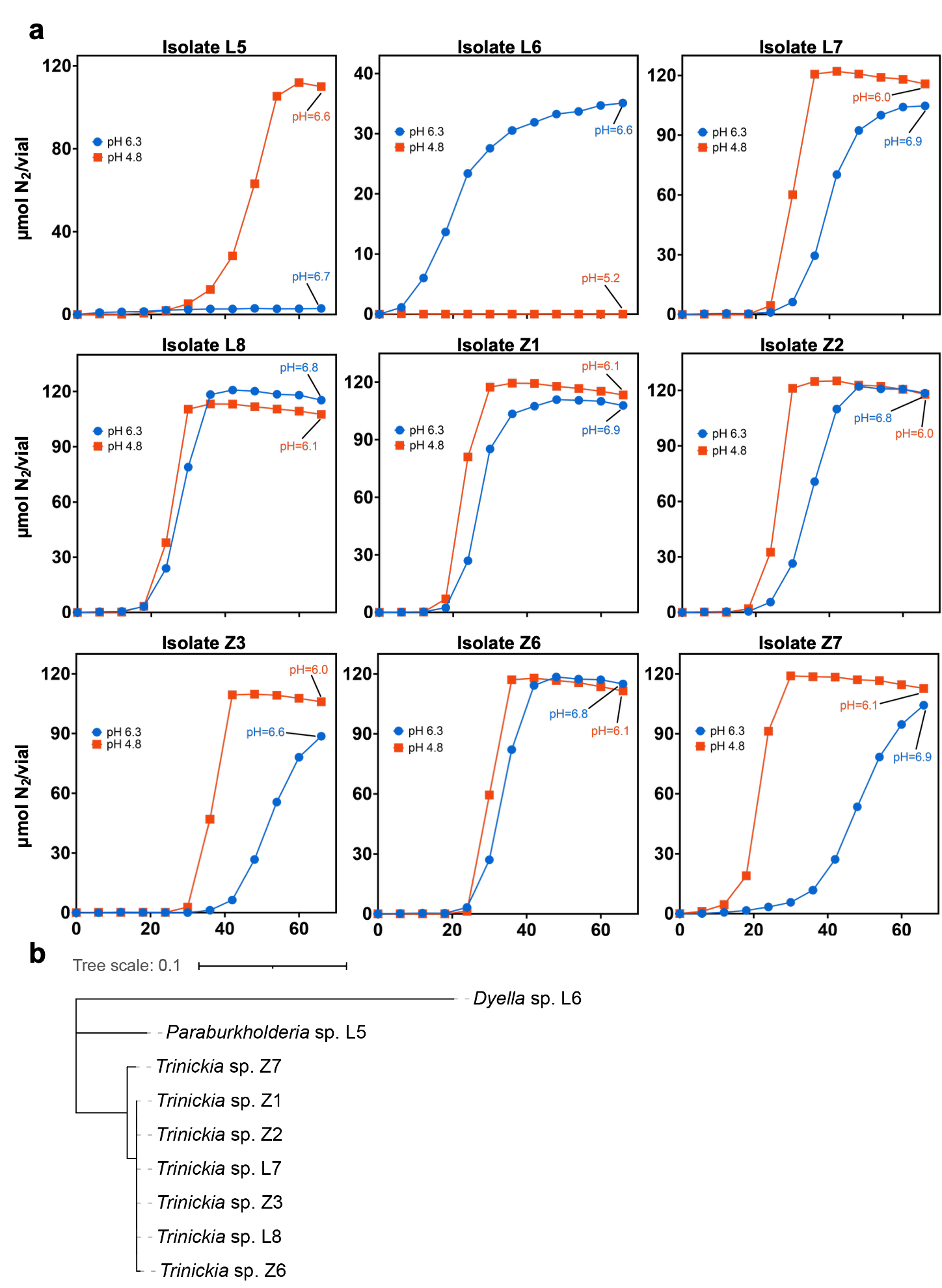
**Supplementary Fig. 7 | N_2_ production and taxonomy of identified N_2_O reducers.** (**a**) N_2_ production of different isolates in acidic (pH 4.8) and circumneutral (pH 6.3) 1/10 TSB media. The inocula were first raised in natural pH 1/10 TSB medium. 300 μL of seeds were inoculated into 30 mL of media with 3 mL of N_2_O in the headspace. No buffer was used to maintain the medium pH. (**b**) Taxonomy and phylogenetic tree of 9 N_2_O reducers based on their 16S rRNA gene sequences.

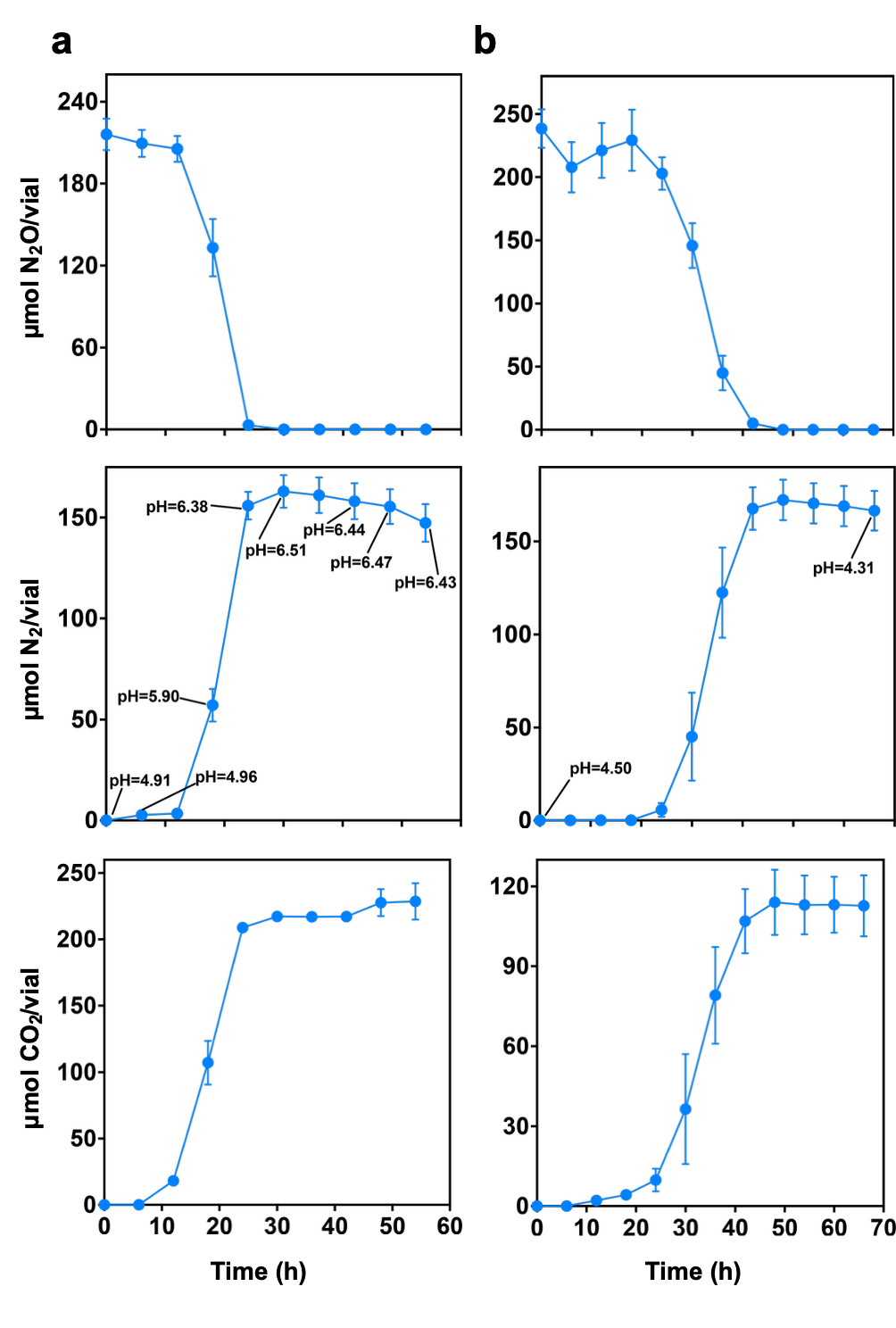
**Supplementary Fig. 8 | Gas kinetics and pH changes of *Trinickia* sp. Z7 cultured in different acidic media.** (**a**) In 1/10 TSB medium with an initial pH 4.9. (**b**) In soil extract-glucose medium with initial pH 4.5. 300 μL of seeds were inoculated into 30 mL of media with 5 mL of N_2_O in the headspace. No buffer was used to maintain the medium pH.

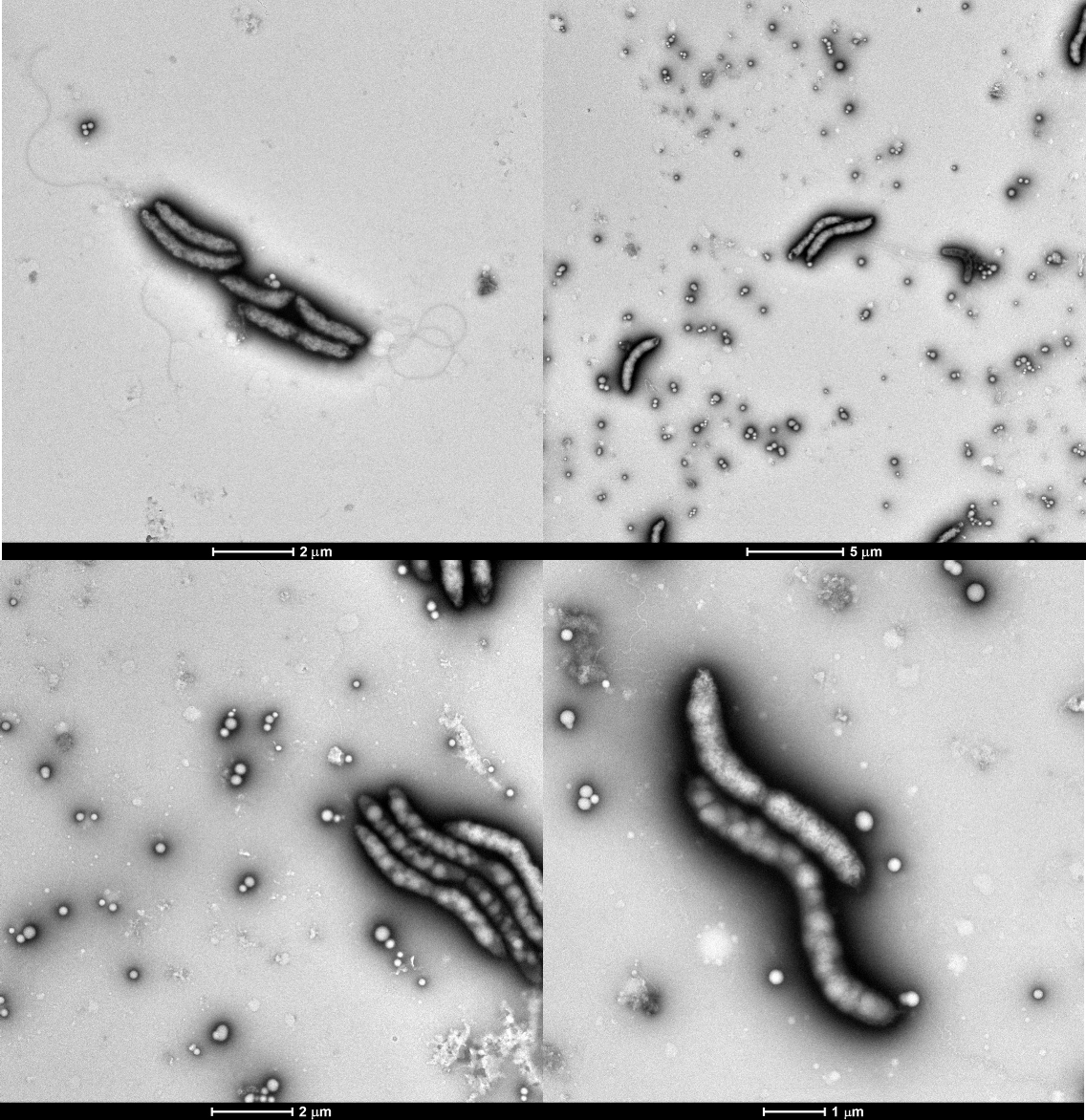

**Supplementary Fig. 9 | Transmission electron micrographs (TEM) of *Trinickia* sp. Z7 cells.** *Trinickia* sp. Z7 was cultured aerobically in pH 4.2 Sistrom's medium with glucose as the sole carbon source. Medium was removed by centrifugation, obtaining cells and secretions. The small visible particles are most likely to be PHB granules.

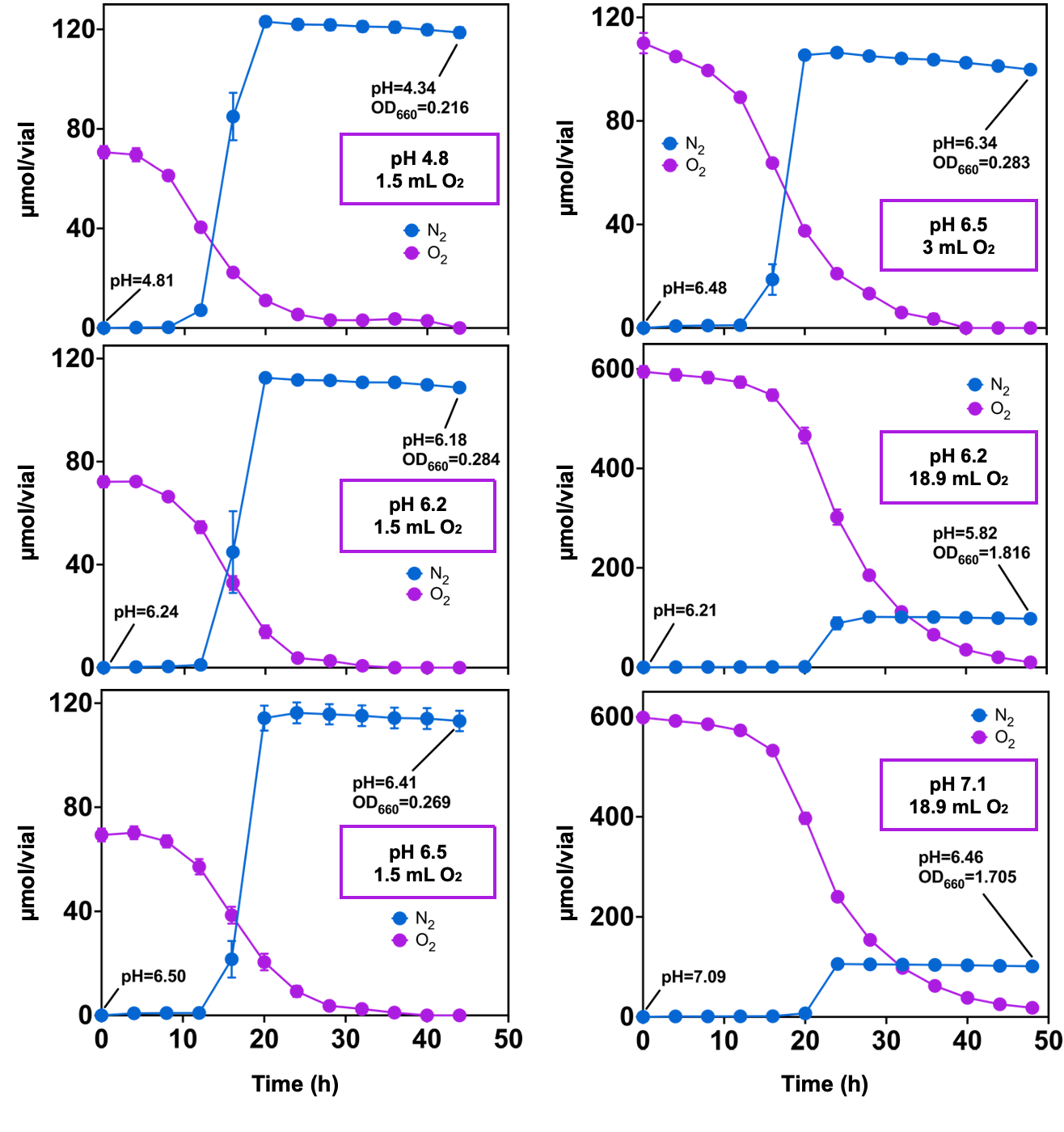

**Supplementary Fig. 10 | N_2_O reduction of *Trinickia* sp. Z7 cultured in different MES buffered Sistrom's media with the presence of O_2_.** Cells were precultured for 3 days to become fully acidic before inoculation, and N_2_O reduction was monitored in 30 mL of 20 mM MES buffered Sistrom's medium containing 300 μL seeds.

**Supplementary Fig. 11 | N_2_O reduction of *Trinickia*** Δ***nosPQ*::*Gm* mutant**. (**a**) N_2_O reduction in *Trinickia* Δ*nosPQ*::*Gm* cells transformed with different reconstructed plasmids. *nosP* and *nosQ* genes were integrated into the pBBR1MCS-2 plasmid.
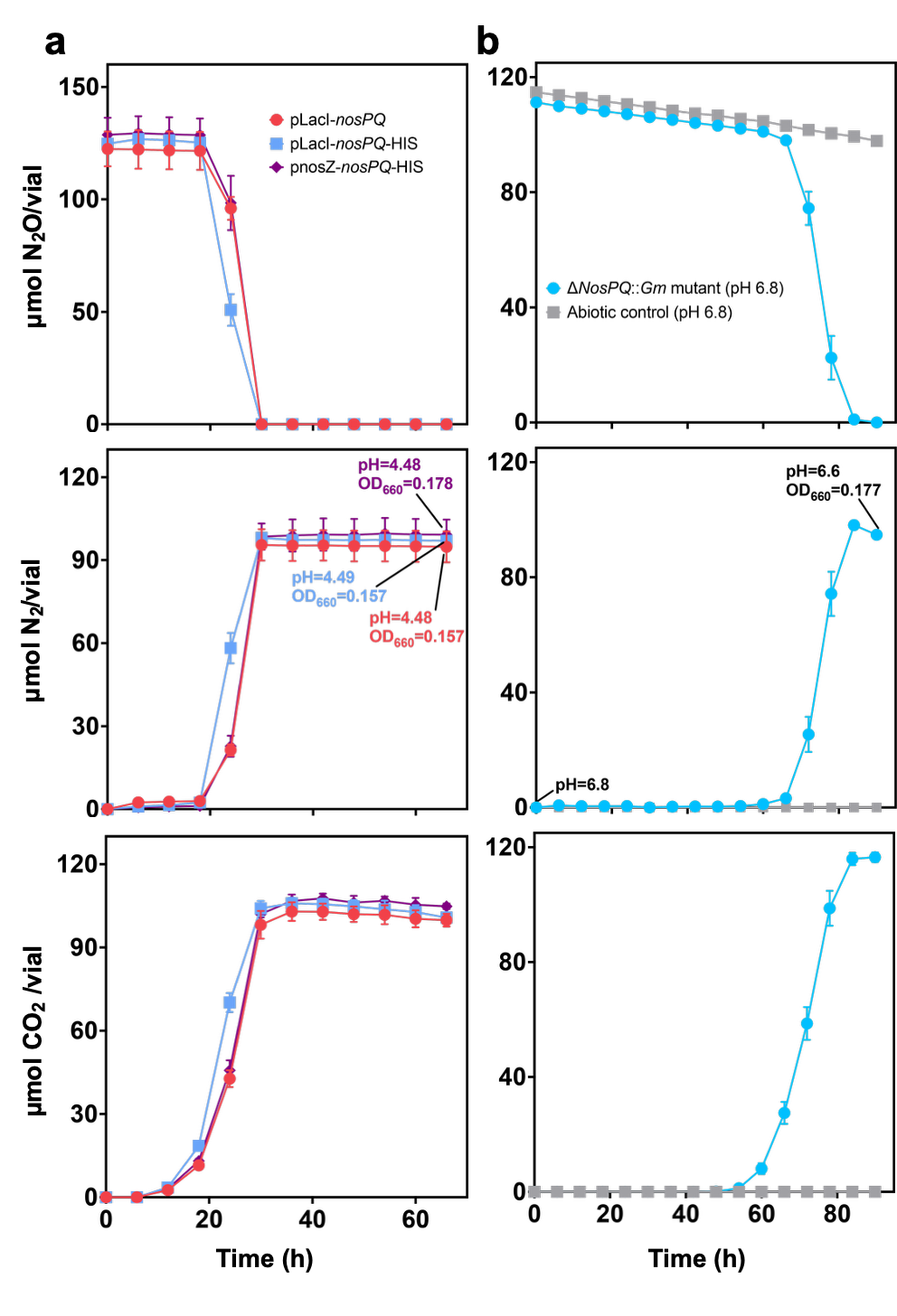
Promoters of pLacI (plasmid's built-in promotor) and pnosZ (derived from the *Trinickia* sp. Z7 *nos* gene cluster) were used for gene expression, respectively. HIS tag was linked to the C-terminal of NosQ. (**b**) N_2_O reduction of mutant cells at pH 6.8. 300 μL of prepared inoculum was inoculated into 30 mL of Sistrom's medium with continuous stirring at 25℃. Twelve replicates were prepared and monitored gas kinetics. The control vials contained the same medium but without bacterial inoculum. At the end of incubation, the cells in each vial were centrifuged and collected to obtain the N_2_O reductases synthesized at circumneutral pH without NosPQ complex.

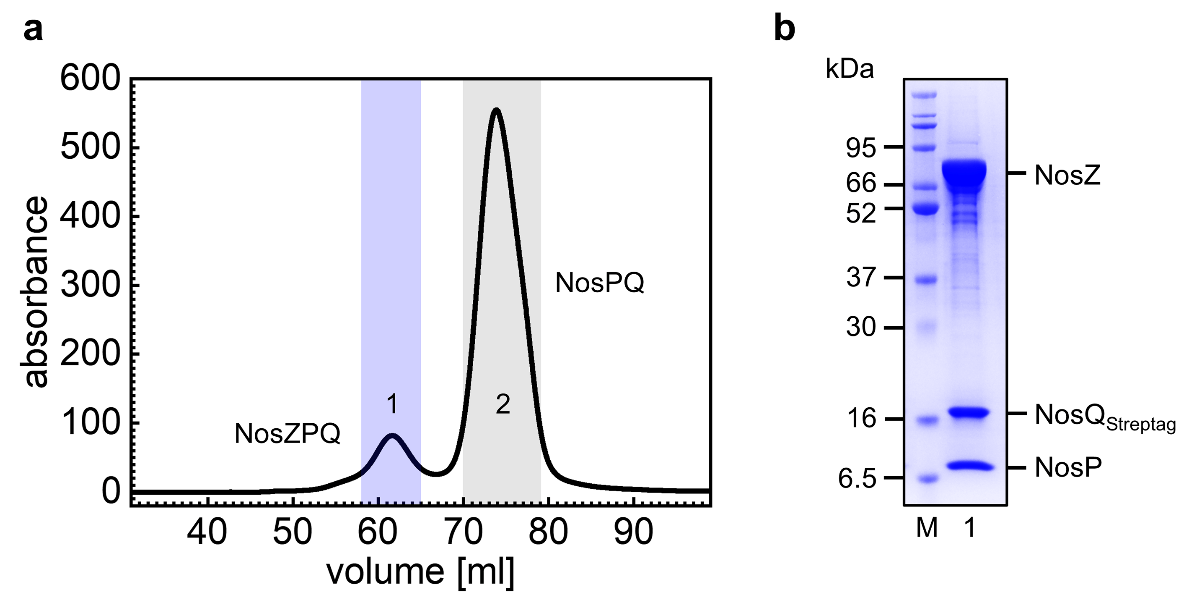
**Supplementary Fig. 12 |Production of the NosZPQ complex.** (**a**) Size-exclusion chromatography (SEC) profile of the NosZPQ complex purification. The fractions highlighted in blue contained the NosZPQ complex, and peak 2 only contained NosPQ heterodimer. (**b**) SDS-PAGE analysis of the complex. The calculated molecular masses (kDa) of the subunits are: 73.1 kDa (NosZ_Histag_), 15.9 kDa (NosQ_Streptag_) and 9.9 kDa (NosP).

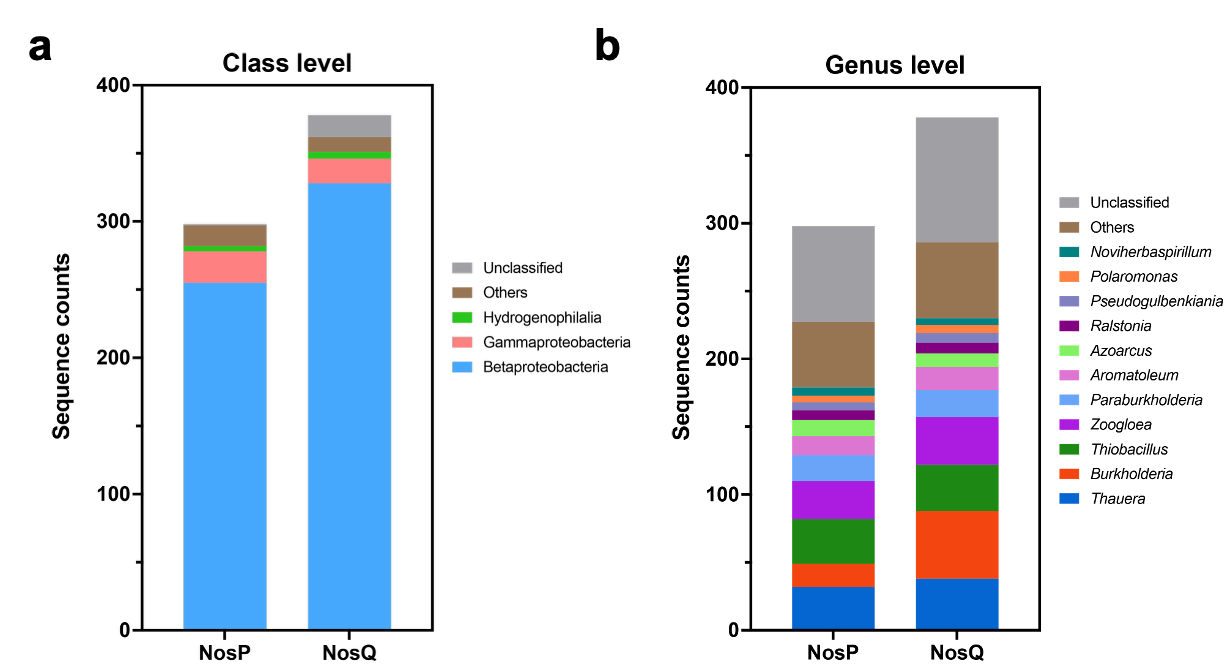

**Supplementary Fig. 13 | Taxonomy and counts of NosP and NosQ sequences recruited from publicly available database at class (left panel) and genus level (right panel), respectively.** The NosP and NosQ sequences from *Trinickia* sp. Z7 were used as query sequences for NCBI Protein BLAST search. The expected threshold was set at 0.00001. Only non-redundant protein sequences from NCBI were used.

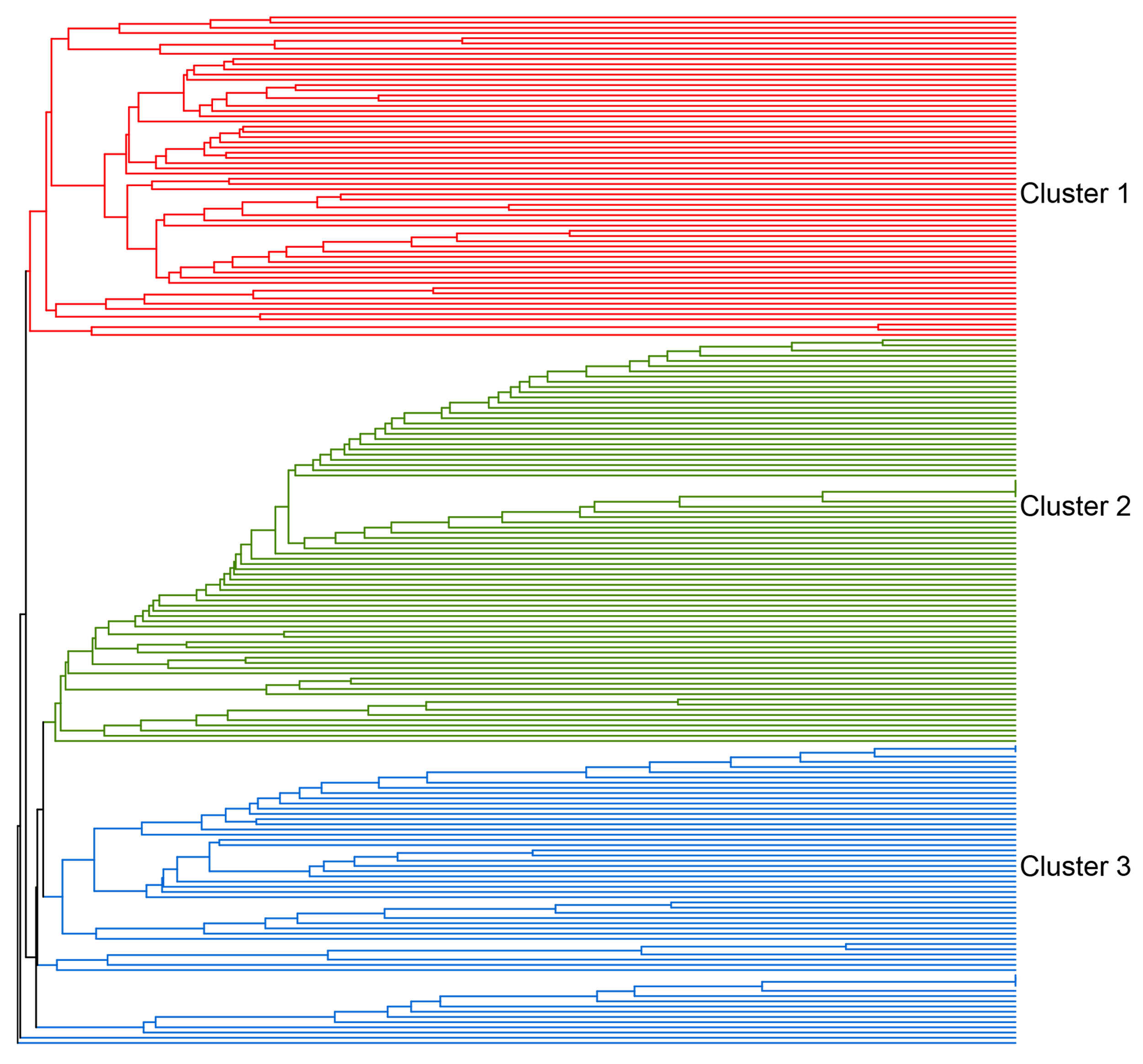

**Supplementary Fig. 14 | Sample clustering by UPGMA based on the microbial composition of *nosPQ* genes.** The calculation and visualization of clustering was done in R by the “vegan” and “phangorn” packages. The proportion of reads identified as unclassified in each sample was removed before the calculation. Cluster 1, Cluster 2 and Cluster 3 contain 62, 78, 58 samples, respectively. Each branch represents a soil metagenome.

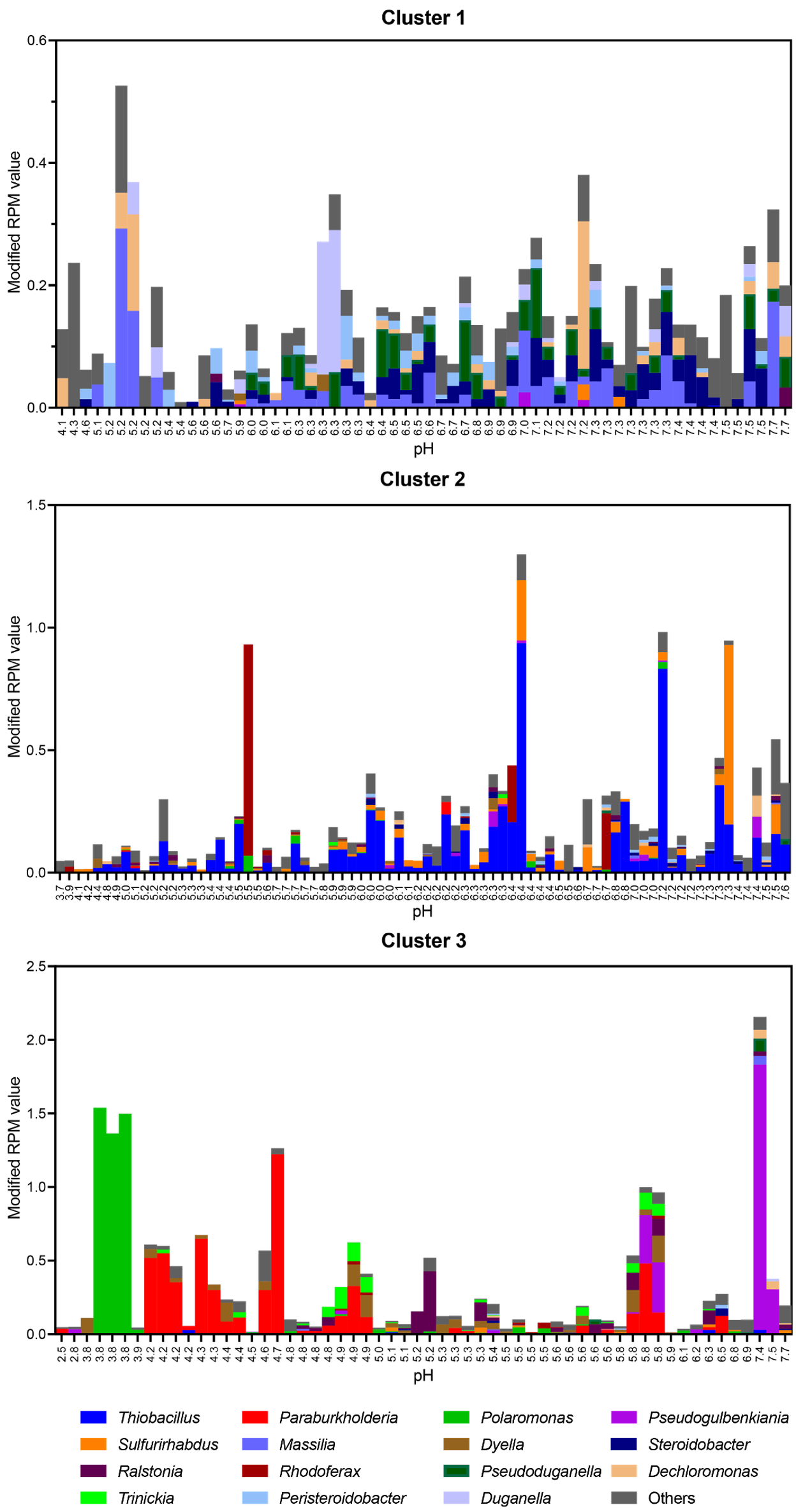

**Supplementary Fig. 15 | The *nosPQ* containing microbial composition and pH of each sample in three clusters.** All samples meet the threshold of RPM value >0.1 and count of total mapped reads ≥10. However, to make the results more clearly, the high proportion of unclassified reads were removed.

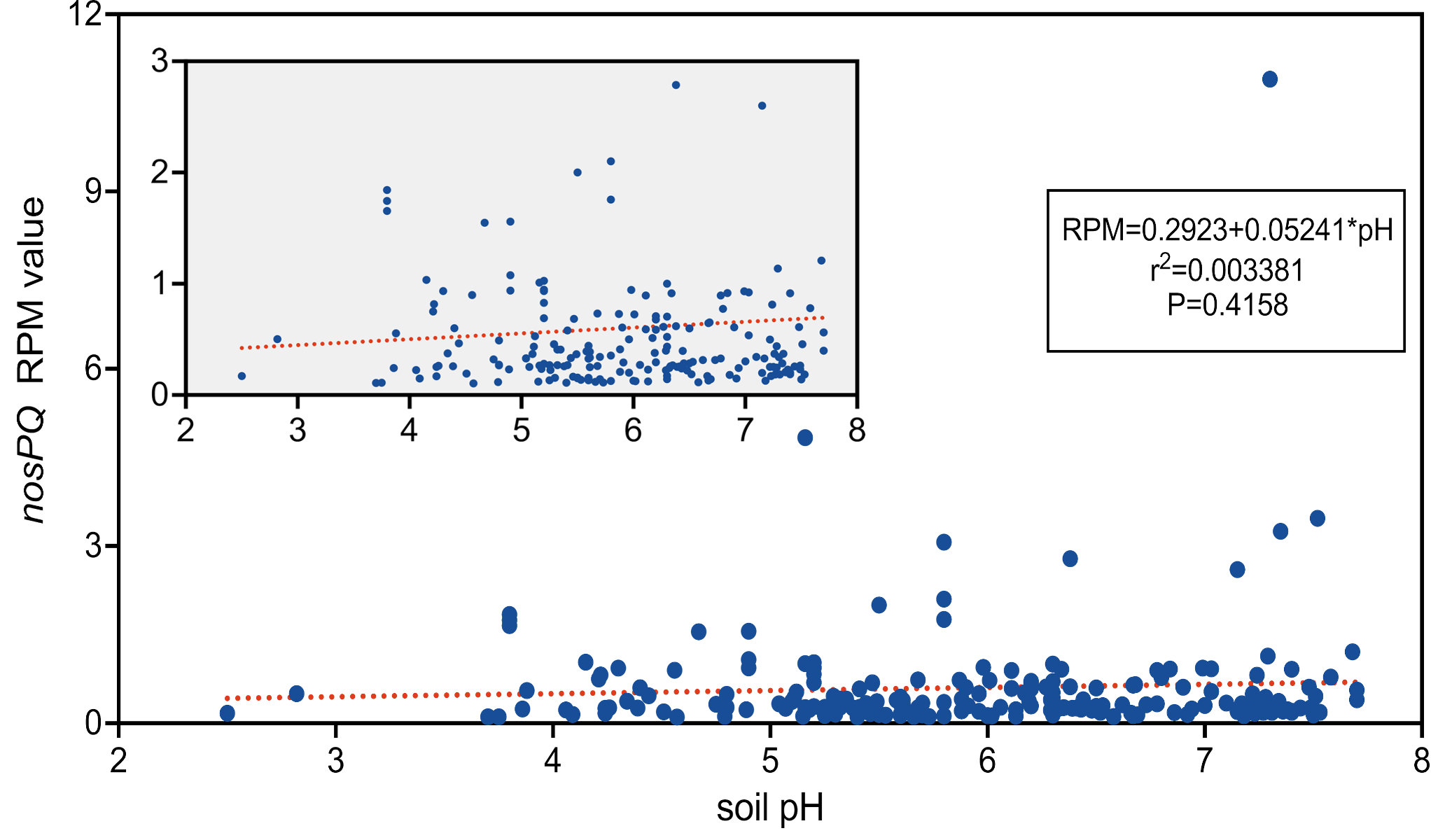

**Supplementary Fig. 16 | The abundance of *nosPQ* gene abundance as affected by soil pH.** The panel shows the abundance of *nosPQ* genes (RPM), plotted against the pH of the soil from which the metagenomes had been extracted. The insert shows the same data scaled differently, to illustrate the trend for metagenomes with RPM values <3 more clearly. The box shows the result of the linear regression analysis, and red line illustrated regression function.

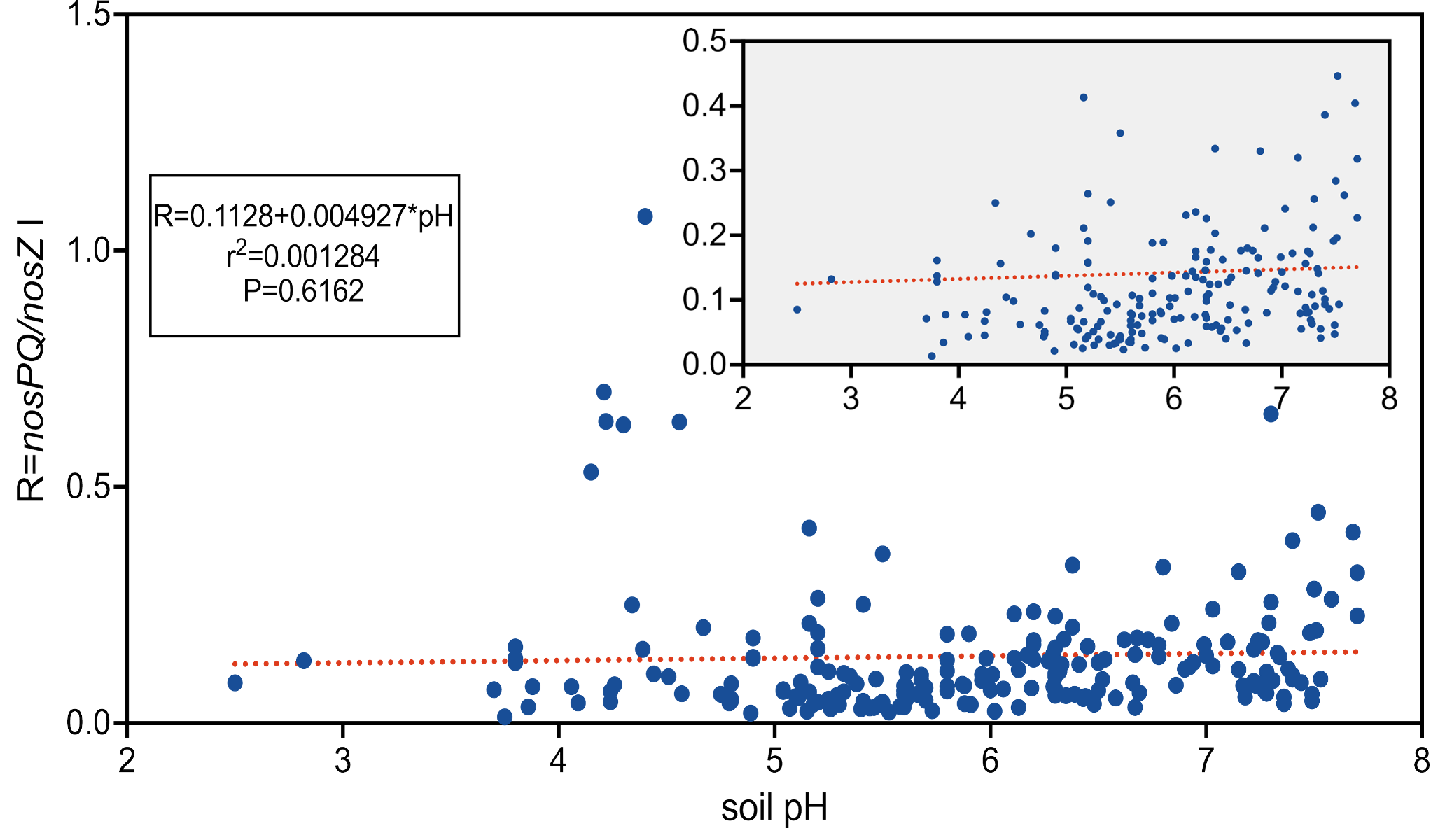

**Supplementary Fig. 17 | The ratio between the abundance of *nosPQ*- and *nosZ* clade I genes as affected by soil pH.** The panel shows the ratio between the abundance of *nosPQ*- and *nosZ* clade I-reads in the soil metagenomes (R), plotted against the pH of the soil from which the metagenomes had been extracted. The insert shows the same data scaled differently, to illustrate the trend for R=0-0.5 more clearly. The box shows the result of the linear regression analysis, and red line illustrated regression function.

### References

1	Wang, X. *et al.* Using adaptive and aggressive N2O-reducing bacteria to augment digestate fertilizer for mitigating N2O emissions from agricultural soils. *Sci Total Environ* **903** (2023). <https://doi.org/10.1016/j.scitotenv.2023.166284>

2	Menzel, P., Ng, K. L. & Krogh, A. Fast and sensitive taxonomic classification for metagenomics with Kaiju. *Nature Communications* **7**, 11257 (2016). <https://doi.org/10.1038/ncomms11257>

3	Li, D. H., Liu, C. M., Luo, R. B. et al. . MEGAHIT: an ultra-fast single-node solution for large and complex metagenomics assembly via succinct de Bruijn graph. *Bioinformatics* **31**, 1674–1676 (2015). <https://doi.org/10.1093/bioinformatics/btv033>

4	Kang, D. W. D., Li, F., Kirton, E. et al. MetaBAT 2: an adaptive binning algorithm for robust and efficient genome reconstruction from metagenome assemblies. *Peerj* **7**, e7359 (2019). <https://doi.org/10.7717/peerj.7359>

5	Parks, D. H., Rinke, C., Chuvochina, M. et al. Recovery of nearly 8,000 metagenome-assembled genomes substantially expands the tree of life. *Nature Microbiology* **2**, 1533–1542 (2017). <https://doi.org/10.1038/s41564-017-0012-7>

6	Parks, D. H., Imelfort, M., Skennerton, C. T. et al. . CheckM: assessing the quality of microbial genomes recovered from isolates, single cells, and metagenomes. *Genome Res* **25**, 1043–1055 (2015). <https://doi.org/10.1101/gr.186072.114>

7	Olm, M. R., Brown, C. T., Brooks, B. et al. . dRep: a tool for fast and accurate genomic comparisons that enables improved genome recovery from metagenomes through de-replication. *The ISME Journal* **11**, 2864–2868 (2017). <https://doi.org/10.1038/ismej.2017.126>

8	Chaumeil, P. A., Mussig, A. J., Hugenholtz, P. GTDB-Tk: a toolkit to classify genomes with the Genome Taxonomy Database. *Bioinformatics* **36**, 1925–1927 (2020). <https://doi.org/10.1093/bioinformatics/btz848>

9	Aramaki, T., Blanc-Mathieu, R., Endo, H. et al. KofamKOALA: KEGG Ortholog assignment based on profile HMM and adaptive score threshold. *Bioinformatics* **36**, 2251–2252 (2020). <https://doi.org/10.1093/bioinformatics/btz859>

10	An, F. Y. *et al.* Metatranscriptome-based investigation of flavor-producing core microbiota in different fermentation stages of dajiang, a traditional fermented soybean paste of Northeast China. *Food Chem* **343** (2021). <https://doi.org/10.1016/j.foodchem.2020.128509>

11	Bolger, A. M., Lohse, M. & Usadel, B. Trimmomatic: a flexible trimmer for Illumina sequence data. *Bioinformatics* **30**, 2114–2120 (2014). <https://doi.org/10.1093/bioinformatics/btu170>

12	Bankevich, A. *et al.* SPAdes: A New Genome Assembly Algorithm and Its Applications to Single-Cell Sequencing. *J Comput Biol* **19**, 455–477 (2012). <https://doi.org/10.1089/cmb.2012.0021>

13	Besemer, J., Lomsadze, A. & Borodovsky, M. GeneMarkS: a self-training method for prediction of gene starts in microbial genomes. Implications for finding sequence motifs in regulatory regions. *Nucleic Acids Res* **29**, 2607–2618 (2001). <https://doi.org/10.1093/nar/29.12.2607>

14	Minh, B. Q. *et al.* IQ-TREE 2: New Models and Efficient Methods for Phylogenetic Inference in the Genomic Era. *Mol Biol Evol* **37**, 1530–1534 (2020). <https://doi.org/10.1093/molbev/msaa015>

15	Letunic, I. & Bork, P. Interactive Tree of Life (iTOL) v6: recent updates to the phylogenetic tree display and annotation tool. *Nucleic Acids Res* (2024). <https://doi.org/10.1093/nar/gkae268>

16	Jain, C., Rodriguez-R, L. M., Phillippy, A. M., Konstantinidis, K. T. & Aluru, S. High throughput ANI analysis of 90K prokaryotic genomes reveals clear species boundaries. *Nature Communications* **9**, 5114 (2018). <https://doi.org/10.1038/s41467-018-07641-9>

17	Kim, D., Park, S. & Chun, J. Introducing EzAAI: a pipeline for high throughput calculations of prokaryotic average amino acid identity. *J Microbiol* **59**, 476–480 (2021). <https://doi.org/10.1007/s12275-021-1154-0>

18	Chen, S. F. Ultrafast one-pass FASTQ data preprocessing, quality control, and deduplication using fastp. *Imeta* **2**, e107 (2023). <https://doi.org/10.1002/imt2.107>

19	Langmead, B., Trapnell, C., Pop, M. & Salzberg, S. L. Ultrafast and memory-efficient alignment of short DNA sequences to the human genome. *Genome Biol* **10**, R25 (2009). <https://doi.org/10.1186/gb-2009-10-3-r25>

20	Anders, S., Pyl, P. T. & Huber, W. HTSeq-a Python framework to work with high-throughput sequencing data. *Bioinformatics* **31**, 166–169 (2015). <https://doi.org/10.1093/bioinformatics/btu638>

21	Guo, X. Y., Wu, K. C., Dong, C. Z. et al. . Paraburkholderia flagellata sp. nov. and Paraburkholderia adhaesiva sp. nov., two novel species isolated from forest soil in Dinghushan Biosphere Reserve in Guangdong, China. *Anton Leeuw Int J G* **116**, 1023–1035 (2023). <https://doi.org/10.1007/s10482-023-01867-4>

22	Chen, W. M., de Faria, S. M., Straliotto, R. et al. Proof that Burkholderia Strains Form Effective Symbioses with Legumes: a Study of Novel Mimosa-Nodulating Strains from South America. *Appl Environ Microb* **71**, 7461–7471 (2005). <https://doi.org/10.1128/Aem.71.11.7461-7471.2005>

23	Lu, C. H. *et al.* Ralstonia chuxiongensis sp. nov., Ralstonia mojiangensis sp. nov., and Ralstonia soli sp. nov., isolated from tobacco fields, are three novel species in the family Burkholderiaceae. *Front Microbiol* **14**, 1179087 (2023). <https://doi.org/10.3389/fmicb.2023.1179087>

24	Dahal, R. H., Chaudhary, D. K., Kim, D. U. & Kim, J. Zoogloea dura sp. nov., a N2-fixing bacterium isolated from forest soil and emendation of the genus Zoogloea and the species Zoogloea oryzae and Zoogloea ramigera *Int J Syst Evol Micr* **70**, 5312–5318 (2020). <https://doi.org/10.1099/ijsem.0.004416>

25	Ngo, H. T. T., Kim, H., Trinh, H. & Yi, T. H. Aquitalea aquatilis sp. nov., isolated from Jungwon waterfall. *Int J Syst Evol Micr* **70**, 4903–4907 (2020). <https://doi.org/10.1099/ijsem.0.004351>

26	Li, Y., Clough, T. J., Moinet, G. Y. K. et al. . Emissions of nitrous oxide, dinitrogen and carbon dioxide from three soils amended with carbon substrates under varying soil matric potentials. *Eur J Soil Sci* **72**, 2261–2275 (2021). <https://doi.org/10.1111/ejss.13124>

27	Shaaban, M., Peng, Q., Lin, S. et al. Nitrous oxide emission from two acidic soils as affected by dolomite application. *Soil Res* **52**, 841–848 (2014). <https://doi.org/10.1071/SR14129>

28	Wu, H. T., Hao, X. H., Xu, P. et al. . CO2 and N2O emissions in response to dolomite application are moisture dependent in an acidic paddy soil. *J Soil Sediment* **20**, 3136–3147 (2020). <https://doi.org/10.1007/s11368-020-02652-w>

29	Patel, S. & Gupta, R. S. A phylogenomic and comparative genomic framework for resolving the polyphyly of the genus Bacillus: Proposal for six new genera of Bacillus species, Peribacillus gen. nov., Cytobacillus gen. nov., Mesobacillus gen. nov., Neobacillus gen. nov., Metabacillus gen. nov. and Alkalihalobacillus gen. nov. *Int J Syst Evol Micr* **70**, 406–438 (2020). <https://doi.org/10.1099/ijsem.0.003775>

30	Kämpfer, P. *et al.* Pseudoneobacillus rhizosphaerae gen. nov., sp. nov., isolated from maize root rhizosphere. *Int J Syst Evol Micr* **72**, 005367 (2022). <https://doi.org/10.1099/ijsem.0.005367>

31	van den Heuvel, R. N., van der Biezen, E., Jetten, M. S. et al. Denitrification at pH 4 by a soil-derived Rhodanobacter-dominated community. *Environ Microbiol* **12**, 3264–3271 (2010). <https://doi.org/10.1111/j.1462-2920.2010.02301.x>

32	Lycus, P., Bothun, K. L., Bergaust, L. et al. Phenotypic and genotypic richness of denitrifiers revealed by a novel isolation strategy. *The ISME Journal* **11**, 2219–2232 (2017). <https://doi.org/10.1038/ismej.2017.82>

33	He, G. *et al.* Sustained bacterial N2O reduction at acidic pH. *Nature Communications* **15**, 4092 (2024). <https://doi.org/10.1038/s41467-024-48236-x>

34	Royalty, T. M. & Steen, A. D. Theoretical and Simulation-Based Investigation of the Relationship between Sequencing Effort, Microbial Community Richness, and Diversity in Binning Metagenome-Assembled Genomes. *Msystems* **4**, e00384–00319 (2019). <https://doi.org/10.1128/mSystems.00384-19>

35	Oshiki, M., Toyama, Y., Suenaga, T. et al. . N(2)O Reduction by Gemmatimonas aurantiaca and Potential Involvement of Gemmatimonadetes Bacteria in N(2)O Reduction in Agricultural Soils. *Microbes Environ* **37**, ME21090 (2022). <https://doi.org/10.1264/jsme2.ME21090>

36	Zhu, H. H., Guo, J. H., Chen, M. B., Feng, G. D. & Yao, Q. Burkholderia dabaoshanensis sp. nov., a Heavy-Metal-Tolerant Bacteria Isolated from Dabaoshan Mining Area Soil in China. *Plos One* **7**, e50225 (2012). <https://doi.org/10.1371/journal.pone.0050225>

37	Yoo, S. H. *et al.* Burkholderia soli sp nov., isolated from soil cultivated with Korean ginseng. *Int J Syst Evol Micr* **57**, 122–125 (2007). <https://doi.org/10.1099/ijs.0.64471-0>

38	Ramoneda, J., Stallard-Olivera, E., Hoffert, M. et al. Building a genome-based understanding of bacterial pH preferences. *Sci Adv* **9**, eadf8998 (2023). <https://doi.org/10.1126/sciadv.adf8998>

39	Ivaldi, C., Foy, C., Castex, S. et al. . Isolation of new Paraburkholderia strains for polyhydroxybutyrate production. *Lett Appl Microbiol* **76**, ovad082 (2023). <https://doi.org/10.1093/lambio/ovad082>
